# Life finds a way: Integrative phylogenomics resolves an overlooked bivalve order with chromosome fusion and mitochondrial translational-code evolution

**DOI:** 10.64898/2026.08.14.744788

**Authors:** Yi-Tao Lin, Yi-Xuan Li, Xin-Yi Li, Ming Tao, Zhangli Hu, Jingjie Hu, Zhenmin Bao, Jian-Wen Qiu

## Abstract

Resolving deep phylogenetic relationships requires integrating multiple lines of evidence, as distinct evolutionary forces shape signals from different genomic markers. Here, we investigate the systematics of the controversial APPD lineage (Anomiidae, Placunidae, Plicatulidae, and, by inference, Dimyidae) within Pectinida *sensu lato* using phylogenomic, comparative genomic, transcriptomic, proteomic, and morphological approaches. Our analyses consistently recover APPD as a monophyletic lineage sister to Limida and Pectinoidea, divergent at ∼428 Mya. With three novel high-quality genomes, extensive progressive chromosomal fusions demonstrate a reduction in chromosome number of the APPD lineage (6–13), compared with an ancestral 20 molluscan linkage groups (MLGs). Accompanied by extensive intrachromosomal gene-order scrambling, we identify one functional centromere in *Placuna vitream* flanked by two vestigial centromeric remnants on a single chromosome, providing a potential resource for investigating centromere inactivation and neocentromere formation. Mitochondrial genomes of APPD lineage exhibit unprecedented plasticity in translational decoding: *Pododesmus* employs the invertebrate mitochondrial code; *Heteranomia* employs +1 translational frameshifting to bypass in-frame TAG codons, whereas in *Anomia*, *Enigmonia*, *Placuna*, and Plicatulidae, TAA is reassigned to tyrosine and confirmed by proteomic evidence, which supports mitochondrial frameshifting in APPD lineage and defines a novel translation table for bivalves. Integrating phylogenetic distinctiveness, deep divergence, extreme karyotypic restructuring, unique mitochondrial features, and morphological diagnosability, we elevate the APPD lineage into Anomiida ord. nov. This revision resolves long-standing uncertainties for Pectinida *sensu stricto* and Limida, and establishes the APPD lineage as a valuable system for investigating chromosome fusion, centromere evolution, codon reassignment, and translational recoding.

**Classification:** Biological Sciences; Evolution

**SIGNIFICANCE STATEMENT:** We have re-examined a controversial group of marine bivalves (Anomiidae, Placunidae, Plicatulidae, and Dimyidae). Our integrative approach shows that these animals split from scallops and their relatives more than 428 million years ago and have undergone drastic chromosomal fusions that reduced their chromosome number from 20 to as few as 6. Additionally, some species evolved unusual ways of reading their mitochondrial genetic code, either reassigning the stop codon to tyrosine or using +1 translational frameshifting to skip stop signals. The combination of deep evolutionary time and genomic divergence warrants recognizing them as a new order, Anomiida ord. nov. This work, as a case study, demonstrates how chromosome fusion and genetic code variation contribute to invertebrate diversity.

## INTRODUCTION

Large-scale chromosomal rearrangements, particularly fusions and fissions, are key drivers of genome evolution and are tightly linked to speciation rates, rendering them valuable phylogenetic markers in invertebrates (1, 2). In Bivalvia, however, karyotypic evolution is strikingly heterogeneous. The superfamily Pectinoidea retains a karyotype that closely mirrors the bilaterian ancestral configuration (2n = 38), complete with the oldest known homomorphic sex chromosomes (3–5). This same 2n = 38 complement is shared by most members of Venerida, Unionida, Nuculanida, Nuculida, Adapedonta, and Arcida, suggesting an ancient origin and persistence across multiple lineages (6–8). Yet this conservation is far from universal: other orders deviate considerably in chromosomal numbers, with Ostreida at 2n = 20, Pteriida at 28, Mytilida at 28 or 30, and Solemyida at 22, alongside sporadic intra-order exceptions (9–12). This stark contrast, with karyotypic stasis in some clades standing in sharp opposition to extensive rearrangement in others, raises a fundamental question concerning the reliability of chromosomal changes as phylogenetic markers for resolving ancient divergences among major bivalve lineages.

Morphological characters further complicate bivalve systematics. Shell morphology, traditionally used for taxonomic identification, presents a dual challenge. Some distantly related lineages frequently exhibit high homoplasy; for instance, gliding scallops have independently evolved streamlined shells and morphological convergence within Veneridae has contributed to taxonomic uncertainty (13, 14). Conversely, some closely related taxa exhibit striking morphological disparity, as documented in mytilid species that differ markedly in shell shape and other characters despite genetic proximity (15, 16). This dual nature of shell morphology renders morphology-based phylogenies unreliable for reflecting evolutionary relationships. Moreover, molecular studies using limited markers also yield discordant topologies. Within Pectinidae, phylogenies from mitochondrial protein-coding genes (PCGs) versus ribosomal RNA genes are incongruent, suggesting locus sampling affects inference even at the family level (17). More broadly, a well-documented mito-nuclear discordance exists in Bivalvia: mitochondrial markers support the monophyly of Pteriomorphia and Heterodonta, whereas nuclear markers support the monophyly of Heterodonta and Palaeoheterodonta (18). Collectively, these observations indicate that morphological, chromosomal, and limited molecular data are insufficient in isolation to resolve deep bivalve relationships; overcoming these obstacles will likely require integrating multiple lines of evidence within a unified phylogenetic framework.

Beyond these phylogenetic challenges, another layer of complexity lies in the evolution of the mitochondrial genetic code itself. Metazoan mitochondrial genomes were long considered conserved in gene content and organization (19). However, broader phylogenetic sampling has revealed many exceptions, with Mollusca particularly replete with departures from the canonical pattern. Molluscan mitogenomes exhibit extraordinary variation in size, architecture, rearrangements, gene duplications/losses, and even novel genes (20–22). Despite this genomic fluidity, all molluscs share the same invertebrate mitochondrial genetic code (code 5) (23, 24). Within this shared framework, translational flexibility is nonetheless considerable: protein-coding genes can initiate at a range of alternative start codons (ATA, ATT, ATC, GTG, and even TTG) in addition to the canonical ATG, and some genes employ incomplete stop codons (e.g., T or TA) that are presumably completed post-transcriptionally (25–27). Given such mitogenomic and codon-usage plasticity, the apparent conservation of the molluscan mitogenomic code may reflect limited taxonomic sampling, and the possibility of genetic code deviations, which have been observed in other metazoans, should not be dismissed without comprehensive examination (23, 28–30).

The order Pectinida *sensu lato*, including scallops and their allies, is morphologically and ecologically diverse (31). Within it, the families Anomiidae, Placunidae, Plicatulidae, and Dimyidae (hereafter the “APPD lineage”) occupy habitats from intertidal to bathyal zones and exhibit considerable morphological and ecological diversity (32). However, long-term systematic uncertainty exists for this lineage: morphological disparity across families, coupled with both convergence and divergence within Anomiidae, has rendered morphology-based classification particularly challenging, while early studies failed to recover a stable placement for APPD lineage (31, 33, 34). Although recent analyses have consistently placed APPD as the earliest diverging clade within Pectinida *sensu lato*, sister to all other pectinids and Limida (32, 35, 36), this topology fundamentally conflicts with the current taxonomic framework since Limida is nested within Pectinida *sensu lato*, rendering the traditional classification system unable to accommodate this phylogenetic relationship. Our preliminary survey of nuclear genomes from the APPD lineage (excluding Dimyidae due to specimen unavailability) unexpectedly revealed extensive and progressive chromosomal fusions, contrasting sharply with the otherwise conserved karyotype of most pectinids; furthermore, mitochondrial genomes of most examined APPD species contained numerous in-frame TAA or TAG codons, directly challenging the assumed universal conservation of the mitochondrial genetic code in bivalves. These unexpected findings make the APPD lineage a valuable testing ground for addressing the broader questions raised above. Here, we integrate phylogenomic, comparative genomic, and proteomic approaches to reconstruct chromosomal evolution and investigate the molecular mechanisms underlying in-frame stop-codon circumvention. Our results uncover a previously underappreciated trajectory of large-scale chromosomal fusion, document the first instance of mitochondrial genetic code variation within Bivalvia, and provide a robust framework for a fundamental revision of the higher-level classification of Pectinida *sensu lato*.

## RESULTS & DISCUSSION

### Multi-source data supports a robust phylogenetic framework

We assembled chromosome-level genomes for *Anomia chinensis*, *Placuna vitream*, and *Plicatula muricata* (Fig. S1; Table 1 & S5). Genome sizes varied substantially across the three species, with *A. chinensis* (530.9 Mb, 7 chromosomes) and *P. vitream* (548.9 Mb, 8 chromosomes) being considerably smaller than *P. muricata* (2.21 Gb, 13 chromosomes). BUSCO completeness was high across all genomes (96.7%, 96.8%, and 95.9% for *A. chinensis*, *P. vitream*, and *P. muricata*, respectively; Table 1), and functional annotation was achieved for 84.0–92.2% of predicted genes (Table 1).

**Table 1.** Characteristics of three nuclear genomes assembled in this study.

| Item | *Anomia chinensis* | *Placuna vitream* | *Plicatula muricata* |
| --- | --- | --- | --- |
| Total length (Mb) | 530.9 | 548.9 | 2,207.8 |
| No. of chromosome | 7 | 8 | 13 |
| GC content (%) | 32 | 33 | 37 |
| N50 (Mb) | 94.7 | 83.5 | 214.9 |
| Genome coverage (X) | 153.1 | 45.5 | 33.4 |
| Mapping rate (%) | 99.9 | 99.8 | 99.9 |
| Assembly BUSCO Metazoa_odb10 | C:96.7%; S: 96.5%; D: 0.2%;<br>F: 0.5%; M: 2.8% | C:96.8%; S: 94.8%; D: 2.0%;<br>F: 0.4%; M: 2.8% | C:95.9%; S: 94.7%; D: 1.3%;<br>F: 0.4%; M: 3.7% |
| Protein-coding genes | 22,896 | 22,292 | 27,966 |
| Average length (AA) | 434.9 | 440.4 | 413.7 |
| With annotation (%) | 86.8% | 84.0% | 92.2% |
| Protein BUSCO Metazoa_odb10 | C:97.5%; S:97.0%; D:0.5%;<br>F:0.8%; M:1.7% | C:97.1%; S:95.2%; D:1.9%;<br>F:0.8%; M:2.1% | C:94.9%; S:94.3%; D:0.5%;<br>F:0.8%; M:4.3% |

Complete BUSCOs (C); Complete and single-copy BUSCOs (S); Complete and duplicated BUSCOs (D); Fragmented BUSCOs (F); Missing BUSCOs (M).

Our combined analyses of nuclear genomic single-copy orthologs, mitochondrial genomes, and shell morphological characters consistently recover the APPD lineage as an independent monophyletic group (Fig. 1a & 2; Table S1). Nuclear and mitochondrial phylogenies consistently recover this lineage as sister to the Limida and Pectinoidea (Fig. 1a & 2a; Table S2), which resolved the monophyly of these three clades. For the morphological component, the maximum parsimony tree based on discrete shell characters (Fig. 2b) recovered all families (including Dimyidae) as monophyletic, with strong support (bootstrap values of 0.94–1.00), and yielded an overall topology largely congruent with the molecular phylogenies at the family and ordinal levels. One topological discrepancy was observed: *Pododesmus* was resolved within Anomiidae in the morphological tree, consistent with its traditional taxonomic placement, rather than forming a sister group to Plicatulidae as in the molecular phylogeny (Fig. 1a & 2). This may reflect morphological homoplasy within Anomiidae, where shell characters often exhibit plasticity and overlapping diagnostic traits, a recurring challenge in pteriomorphian systematics (32, 37). Nevertheless, the overall congruence across independent datasets provides robust support for the major phylogenetic conclusions and underscores the distinctiveness of the APPD lineage. Importantly, the APPD lineage and Limida are recovered as monophyletic lineages in ((APPD + (Limida + Pectinoidea)), which falls outside the long-standing controversial topology — APPD and Limida nested within Pectinida *sensu lato*. These results align with recent phylogenetic studies that consistently recover the APPD lineage as the earliest-diverging clade among three controversial clades (32, 35, 36). Although genomic resources of Dimyidae were unavailable for comparison, their placement as the sister group of Plicatulidae within the APPD lineage is supported by both morphological evidence (Fig. 2b; Table S3; Supplementary Information) and previous molecular phylogenetic analyses based on single-gene markers (32, 35). Together with phenotypic analyses that place Dimyidae between ostreoids and pectinoids and Triassic fossil evidence that excludes their close ties to Ostreidae, these observations support the inclusion of Dimyidae within the APPD lineage (38, 39).

**Fig. 1.**
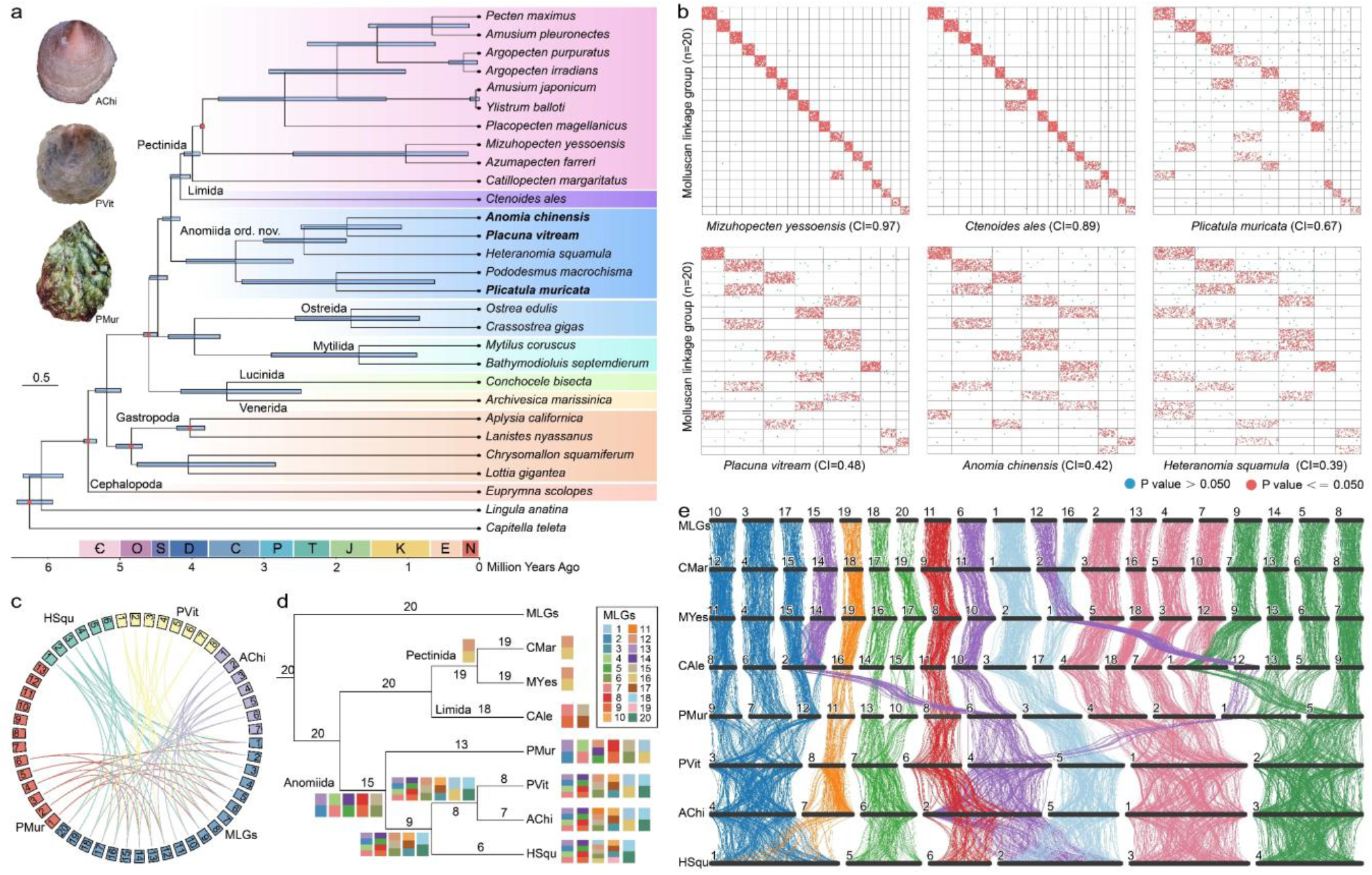
Phylogenomic relationships, chromosomal fusion scenario, and synteny. (a) Maximum-likelihood phylogenomic tree with divergence time estimates. The tree was constructed using 714 single-copy orthogroups (at least 80% taxon occupancy), with an annelid and a brachiopod as outgroups. Bootstrap support is 100 at all nodes. Six calibration points (red dots) based on fossils were applied (Table S4). Estimated divergence times are shown as blue bars, with lengths indicating 95% confidence intervals. Genome information is provided in Table S2. Different orders are shaded in distinct background colors. Specimen photographs of the three newly assembled species are shown: *Anomia chinensis* (2 cm), *Placuna vitream* (7 cm), and *Plicatula muricata* (6 cm). (b) Macrosynteny dot plots showing the retention levels of the ancestral molluscan linkage groups (MLGs, n=20) across different species, measured as the conserved index (CI). Each dot represents a pair of homologous genes. (c) Circular synteny plot illustrating the fusion relationships of MLGs in the selected three anomiids. Colors distinguish different species and chromosome numbers are labeled. (d) Chromosomal fusion scenario of MLGs within Anomiida ord. nov., Pectinida *sensu stricto*, and Limida. Differently colored rectangles represent each MLG, and stacked rectangles indicate fused chromosomes. The inferred chromosome number at each node is given. (e) Microsynteny among representative species of Anomiida ord. nov., Pectinida *sensu stricto*, and Limida. Each black horizontal bar represents a chromosome; bar lengths reflect relative chromosome sizes within each species and are not comparable across species. Chromosomes sharing the same fusion history are connected by color-coded ribbons. AChi, *Anomia chinensis*; CMar, *Catillopecten margaritatus*; CAle, *Ctenoides ales*; HSqu, *Heteranomia squamula*; MYes, *Mizuhopecten yessoensis*; PMur, *Plicatula muricata*; PVit, *Placuna vitream*.

**Fig. 2.**
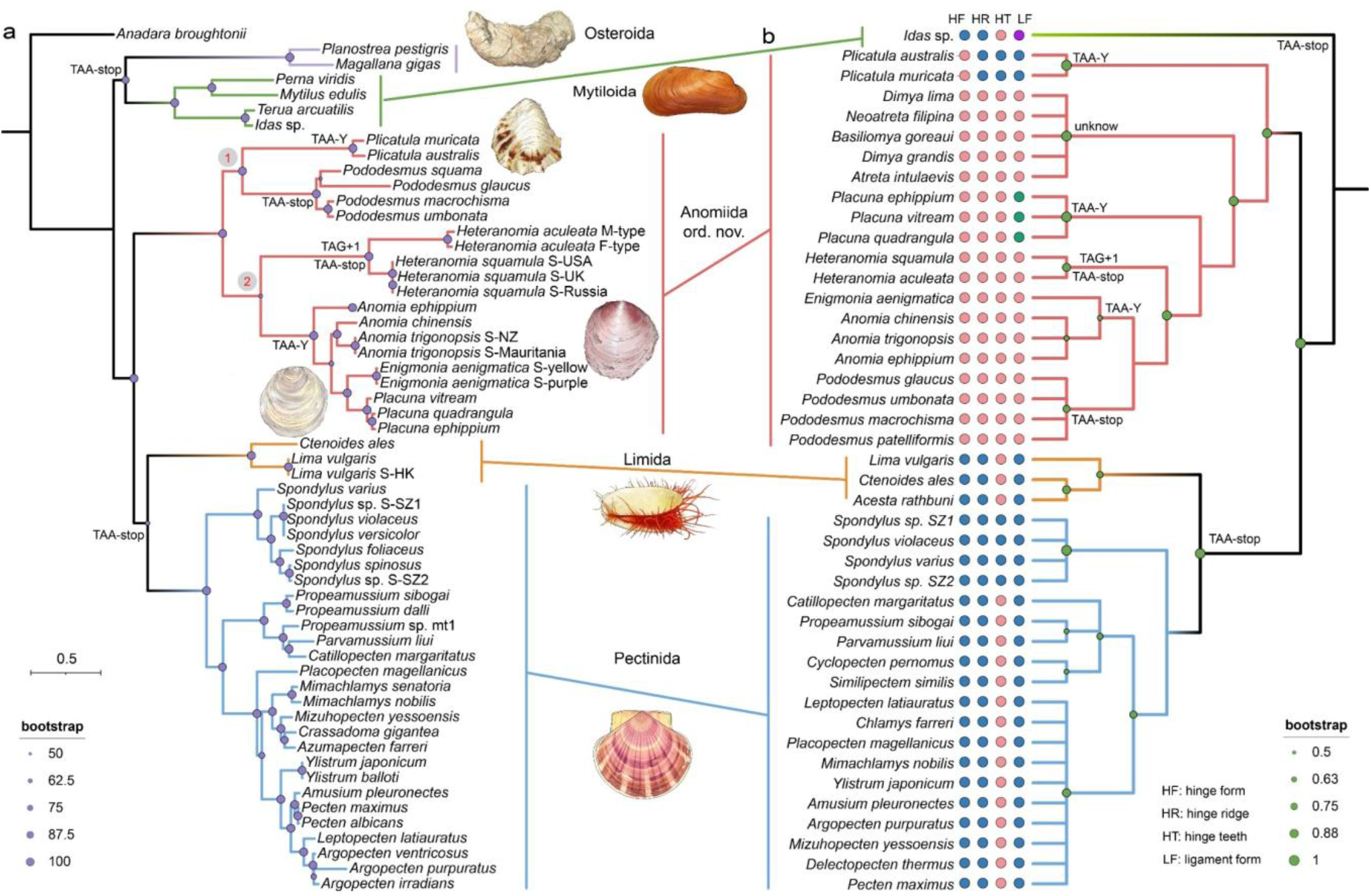
Phylogenetic relationships of the APP lineage and morphological corroboration. (a) Maximum likelihood (ML) tree inferred from concatenated mitochondrial PCGs and rRNA genes, with *Anadara broughtonii* (Arcida) as the outgroup. The tree shows the phylogenetic relationships among Ostreida, Mytilida, Limida, Pectinida *sensu stricto*, and the newly recognized Anomiida ord. nov. Nodes 1 and 2 indicate two independent origins of TAA=Tyr reassignment. Different orders are color-coded. Mitochondrial genome configurations are annotated at key nodes. (b) Maximum parsimony (MP) tree based on 21 discrete shell characters, with a Mytiloida species as the outgroup. Morphological data for Dimyidae were obtained from the literature (85, 86). Key hinge-related characters distinguishing lineages are illustrated. Orders are distinguished using colors as in (a). The topology is largely congruent with molecular phylogenies, except that *Pododesmus* falls within Anomiidae rather than as sister to Plicatulidae.

Time-calibrated phylogenetic analyses based on genomic data further revealed the ancient, independent evolutionary history of the APPD lineage (Fig. 1a; Table S4). The estimated divergence time between the APPD lineage and the Limida and Pectinoidea clade is approximately 428.57 million years ago (MYA). This date is not only earlier than the divergences of other pectinidan families, but also predates the ordinal-level splits between Mytilida and Ostreida (approximately 395.87 MYA) and between Venerida and Lucinida (approximately 351.12 MYA). In bivalves, time-calibrated phylogenies have increasingly been employed to reassess higher-level classifications, with deep divergence times often correlating with ordinal-level distinctions (40). The estimated divergence of the APPD lineage, comparable to or older than the ordinal splits between Mytilida and Ostreida and between Venerida and Lucinida, provides a chronostratigraphic rationale consistent with these practices. The temporal depth of this divergence, coupled with the topological distinctiveness established above, supports the distinctiveness of the APPD lineage.

### Karyotype Reduction and Progressive Chromosomal Fusions

With established the phylogenetic position of the APPD lineage, we further investigated whether this distinctiveness is reflected in its nuclear genome architecture. The APPD lineage species exhibited karyotypic divergence from both Pectinidae (n=19), Limida (n=18), and the inferred ancestral karyotype of molluscs (20 molluscan linkage groups, MLGs) (41), with *P. muricata*, *P. vitream*, *A. chinensis*, and *Heteranomia squamula* possessing n=13, 8, 7, and 6, respectively (Fig. 1b-e; Table 1 & S2). Macrosynteny analyses revealed extensive chromosomal homology across the examined genomes, indicating that all four species have undergone substantial fusion events relative to the inferred ancestral state (Fig. 1b & S2). Using MLGs as a reference, we inferred a parsimony-based scenario for ancestral karyotypes and sequential fusion events along the genome-wide phylogeny (Fig. 1c-d; Table 2). The typical scallop lineage (*Mizuhopecten yessoensis* and *Catillopecten margaritatus*) retained a karyotype close to the ancestral complement, with one single fusion event (MLG12+MLG16). Limida showed two independent fusions distinct from those in scallops, suggesting that the most recent common ancestors of both lineages likely maintained the full set of MLGs. In contrast, the common ancestor of the four APPD species already possessed a reduced karyotype of 15 chromosomes, resulting from five independent fusions (Fig. 1d). Subsequent lineage-specific fusion events further reduced chromosome numbers, with the number of retained original MLGs decreasing progressively from 7 (CI = 0.67) in *P. muricata* (n = 13), to 2 (CI = 0.48) in *P. vitream* (n = 8), and to only 1 (CI = 0.42–0.39) in both *A. chinensis* (n = 7) and *H. squamula* (n = 6) (Fig. 1b; Table 2). Moreover, our microsynteny analysis further reveals that fused chromosomes have also undergone extensive gene-order scrambling (Fig. 1e). Together, these lines of evidence demonstrate that the fusion events in the APPD lineage were followed by extensive inter-chromosomal rearrangements that disrupted ancestral linkages (Fig. 1e), not as simple Robertsonian fusions with typically conserved gene order within fused chromosomes (42). This degree of genomic restructuring, as inferred from 20 MLGs to a heavily rearranged karyotype of as few as 6 chromosomes, represents an extreme departure from the stable karyotype evolution inferred for most bivalve lineages and underscores the unique evolutionary trajectory of this clade.

**Table 2.** Inferred chromosomal fusion events in the APPD lineage and related species (based on parsimony reconstruction from extant genomes).

| Lineage / Species | Fusion events | Number of chromosomes | Notes |
| --- | --- | --- | --- |
| Scallop lineage (*Catillopecten margaritatus*, *Mizuhopecten yessoensis*) | MLG12+MLG16 | 19 to 18 | Single fusion, ancestral karyotype largely retained |
| Limida (*Ctenoides ales*) | MLG7+MLG9; MLG15+MLG17 | 20 to 18 | Two independent fusions distinct from scallops |
| Common ancestor of *Plicatula muricata*, *Placuna vitream*, *Anomia chinensis*, *Heteranomia squamula* | MLG2+MLG13; MLG4+MLG7; MLG5+MLG14; MLG8+MLG9; MLG6+MLG15 | 20 to 15 | Five independent fusions at the base of the APPD lineage |
| *Plicatula muricata* (additional fusions) | MLG12 fused with MLG5+MLG14; MLG1 fused with MLG16 | 15 to 13 | Lineage-specific secondary fusions |
| Ancestor of *Placuna vitream*, *Anomia chinensis*, *Heteranomia squamula* | MLG2+MLG13+MLG4+MLG7; MLG5+MLG14+MLG8+MLG9; MLG12+MLG6+MLG15; MLG3+MLG10+MLG17; MLG18+MLG20 | 15 to 9 | Four secondary fusions and one independent fusion |
| Ancestor of *Placuna vitream* and *Anomia chinensis* | MLG1+MLG16 | 9 to 8 | Single fusion after divergence from *Heteranomia squamula* |
| *Placuna vitream* | None | 8 | Karyotype retained from its ancestor |
| *Anomia chinensis* | MLG11 fused with MLG6+MLG12+MLG15 | 8 to 7 | Additional fusion specific to *Anomia chinensis* |
| *Heteranomia squamula* | (MLG12+MLG6+MLG15) fused with (MLG1+MLG16); (MLG3+MLG10+MLG17) fused with MLG19 | 9 to 6 | Two fusions leading to highly reduced karyotype |

The evolutionary significance of chromosomal fusions extends beyond their immediate effects on reproduction; they also provide natural experiments for investigating centromere fate, post-fusion genomic remodeling, and the interplay between karyotype change and speciation (43, 44). Within molluscs, compelling examples of chromosomal fusion have emerged across multiple classes, indicating that recurrent fusion events are not restricted to a single lineage. In Polyplacophora, recent comparative genomic analyses of chitons uncovered surprisingly dynamic karyotype evolution, including independent fusion events that reduced chromosome numbers in distantly related lineages (41). In Gastropoda, chromosomal fusion events have also been documented in several lineages, often associated with lineage diversification and adaptive radiation (45, 46). Even within Pectinida *sensu stricto*, where most families exhibit a conserved haploid number of n=19, an exception has been noted in the bay scallop *Argopecten irradians* (n=16) with limited chromosomal fusions (47, 48). However, beyond this isolated case, large-scale, progressive chromosomal fusions have not been documented previously (3, 5). In contrast, the APPD lineage examined here exhibits extensive and successive chromosomal fusions (Fig. 1), representing a remarkable departure from the otherwise stable karyotype evolution. This discovery positions the APPD lineage as a potential example for investigating the mechanisms and evolutionary consequences of karyotype restructuring.

Additionally, the correlation between the nodes of these fusion events and the divergence of APPD lineages raises the intriguing possibility that karyotype restructuring may have played a causal or reinforcing role in speciation. Chromosomal fusions can promote reproductive isolation by disrupting meiotic pairing in hybrids, thereby contributing to speciation (42, 49). The progressive nature of these fusions, each occurring after lineage divergence, raises the possibility that karyotype evolution in the APPD lineage could involve a process akin to a “fusion ratchet”, whereby each fusion event might incrementally reduce the probability of successful hybridization with ancestral populations. This pattern parallels a chromosomal speciation model proposed for mammals, where Robertsonian fusions are associated with rapid radiation in some groups (50, 51). However, testing this hypothesis will require experimental data on reproductive isolation and hybrid fertility in future studies. Generally, the APPD lineage presents a particularly tractable system for investigating this phenomenon in invertebrates, as the time and order of fusions can be reconstructed from comparative synteny analysis. Moreover, the extreme karyotypic reduction observed in these species represents one of the most dramatic examples of chromosomal fusion documented in any animal lineage. This positions the APPD lineage as a valuable example for investigating the functional consequences of karyotype reduction, including effects on gene expression, recombination rates, and the physical architecture of the nucleus (52, 53).

### Putative vestigial centromeres associated with chromosomal fusion

Centromeres are usually low in tandem repeat diversity compared to other genomic regions. In *P. vitream*, we identified a distinct centromeric region on chr7, spanning 18.35–21.43 Mb, putatively designated as the candidate functional centromere (FC), which is characterized by the massive occupation of only three tandem repeats (Fig. 3 & S3). Synteny analysis revealed that chr7 originated from an ancient chromosomal fusion involving two ancestral chromosomes, MLG18 and MLG20, traceable to the common ancestor of *P. vitream*, *A. chinensis*, and *H. squamula* (Fig. 1c-e). Intriguingly, two additional regions with markedly reduced tandem repeat diversity were detected around 10–12 Mb and 32–36 Mb, flanking the FC on opposite sides (Fig. 3a-b). We identified these regions as potential vestigial centromeres (VC1 and VC2), remnants of ancestral centromeres that have degenerated following the fusion event. Sequence similarity analyses showed that only a limited fraction of VC1 and VC2 share homology with FC, with VC2 exhibiting a greater extent of homology than VC1 (Fig. 3c-d). This pattern is consistent with two alternative scenarios: FC may have originated from the centromere of the same ancestral chromosome that gave rise to VC2, followed by subsequent divergence, or the differential retention of sequence similarity could reflect distinct rates of degeneration between VC1 and VC2. Nevertheless, sliding window GC content analysis revealed a striking compositional contrast among these regions: FC displays extremely low GC content ranging from 22.5% to 25.0%, whereas VC1 and VC2 exhibit substantially higher GC levels between 27.5% and 40.0% (Fig. 3e). This GC divergence may reflect distinct evolutionary trajectories following the fusion event, consistent with the notion that functional centromeres are subject to specific evolutionary constraints, potentially involving centromere-specific histones or chromatin modifications.

**Fig. 3.**
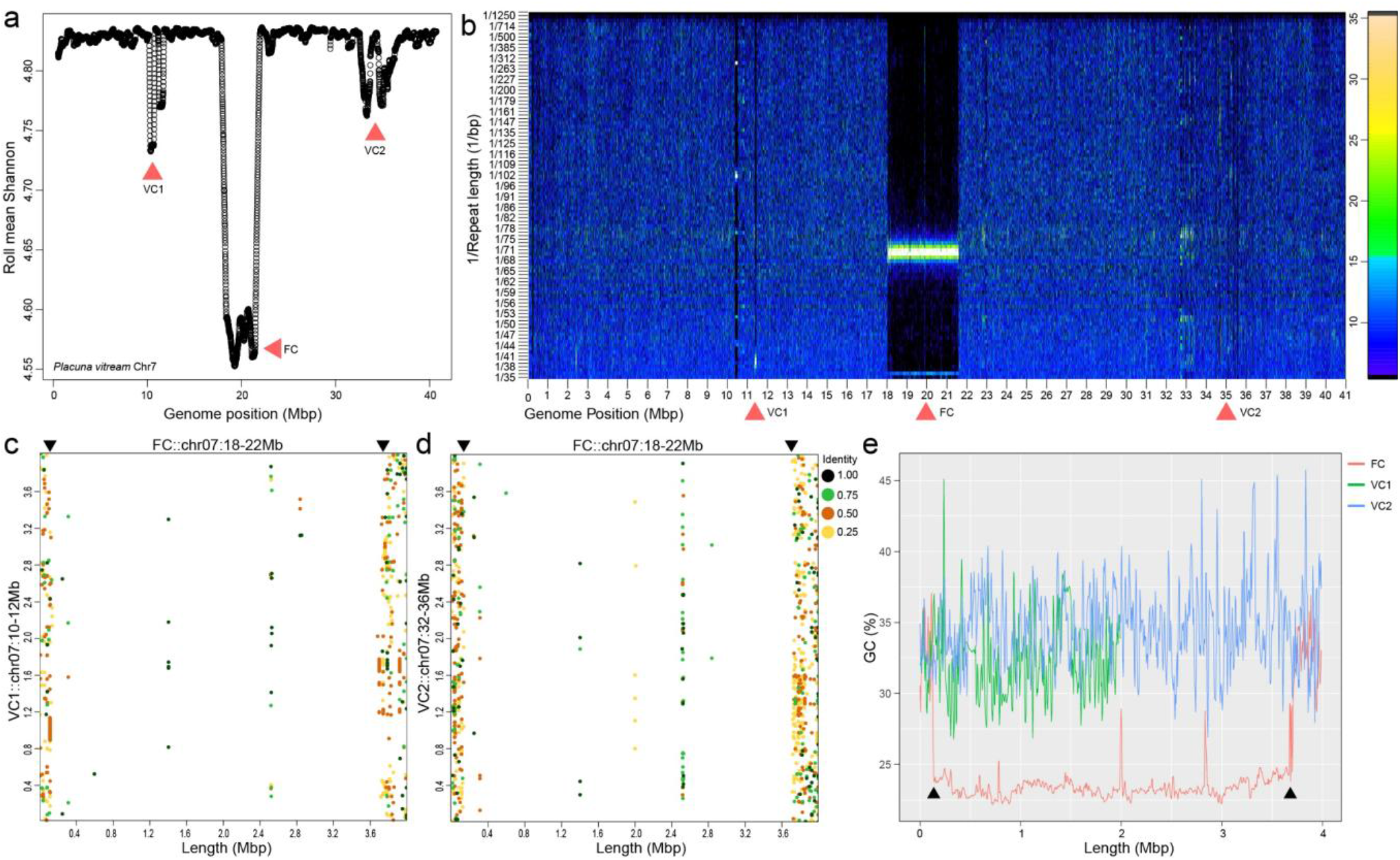
Identification of functional and vestigial centromeres on chr7 of *Placuna vitream*. (a) Rolling means of Shannon diversity index (window size = 100 bp) along chromosome 7. A prominent trough (FC, functional centromere) with exceptionally low sequence diversity is flanked by two shallower troughs (VC1 and VC2, vestigial centromeres), indicating reduced diversity at these loci. (b) Distribution of a 35-bp satellite repeat family along chromosome 7. A bright horizontal band at ∼1/68–1/72 corresponds to the FC, reflecting extremely low repeat diversity. Two additional regions of moderately reduced diversity coincide with VC1 and VC2. (c–d) Dot-plot comparisons of sequence similarity among the three centromeric regions: FC vs. VC1 (c) and FC vs. VC2 (d). (e) Sliding window GC content (window = 10 kb, step = 5 kb) across the three centromeric regions. FC exhibits a markedly lower GC content compared to VC1 and VC2.

The configuration observed in *P. vitream*, one functional centromere accompanied by two vestigial remnants, represents a clear example of centromere inactivation and retention following chromosomal fusion documented in molluscs. Comparable cases have been described in mammals, such as the vestigial centromeres retained on human chromosome 2 following the fusion of two ancestral ape chromosomes (54, 55). In that instance, the vestigial centromere is detectable primarily by molecular cytogenetic methods such as fluorescence in situ hybridization (FISH) with centromeric satellite probes (56). In *P. vitream*, by contrast, the vestigial centromeres are distinguished by their unique tandem repeat composition and, more strikingly, by their markedly elevated GC content relative to the functional centromere (57, 58). This compositional divergence may reflect distinct evolutionary trajectories following the fusion event: the functional centromere remains AT-rich due to centromere-specific chromatin dynamics or recombination suppression, while the vestigial centromeres, no longer under such constraints, have relaxed toward the base composition of surrounding euchromatin (59). Moreover, the presence of two vestigial centromeres on the same chromosome provides a unique opportunity to study centromere inactivation and formation. The greater sequence similarity between VC2 and FC relative to VC1 raises the possibility that FC may have been derived from the centromere of the same ancestral chromosome that gave rise to VC2, with subsequent divergence (58, 60). Alternatively, FC may represent a neocentromere that arose after the fusion event, a phenomenon documented in human clinical cases and some evolutionary contexts (61, 62). The *P. vitream*, with its exceptionally clear structural architecture, provides an unparalleled natural experiment for investigating the mechanisms of centromere inactivation and the genomic features that distinguish functional from vestigial centromeres. Nevertheless, definitive confirmation of centromere functionality and vestigial status— and discrimination between the alternative scenarios of centromere derivation versus neocentromere formation—will require experimental approaches, such as CENH3 chromatin immunoprecipitation (ChIP-seq) or immunofluorescence localization, to directly assess the presence of active centromeric histone variants at these loci. Collectively, these findings document an ancient chromosomal fusion event preserved as a mosaic of one functional and two vestigial centromeres on a single chromosome, revealing the long-term genomic consequences of karyotypic restructuring in this lineage. Along with the progressive fusion events documented above, these observations highlight the astounding extent of genomic rearrangement.

### Mitogenomic Diversity and Translational Adaptations

#### Three mitochondrial configurations

Annotation of mitochondrial genomes from the APPD lineage revealed widespread in-frame stop codons (TAA and TAG) within PCGs of most species analyzed (Fig. 4a & S4-7; Table S6, Supplementary Information), contradicting the traditional view of a strictly conserved bivalve mitochondrial genetic code (63, 64). These observations imply that these marine bivalves likely evolved specific mechanisms to mitigate the potentially deleterious effects of in-frame stop codons (65). Protein homology-based predictions revealed three mitochondrial configurations (Fig. 4b; Table S7): (1) a conventional configuration (*Pododesmus*) employing translation code 5; (2) *Heteranomia* also using code 5 but harboring in-frame TAG codons; and (3) a lineage encompassing *Anomia*, *Enigmonia*, Placunidae, and Plicatulidae where TAA is reassigned to tyrosine while all other codons follow code 5. As this recoding pattern (TAA=Tyr) is distinct from all known genetic codes, including translation table 33 for Cephalodiscidae, which shares TAA=Tyr but differs in other assignments, we designate it as translation code 34 (Fig. 4c; Table S7).

**Fig. 4.**
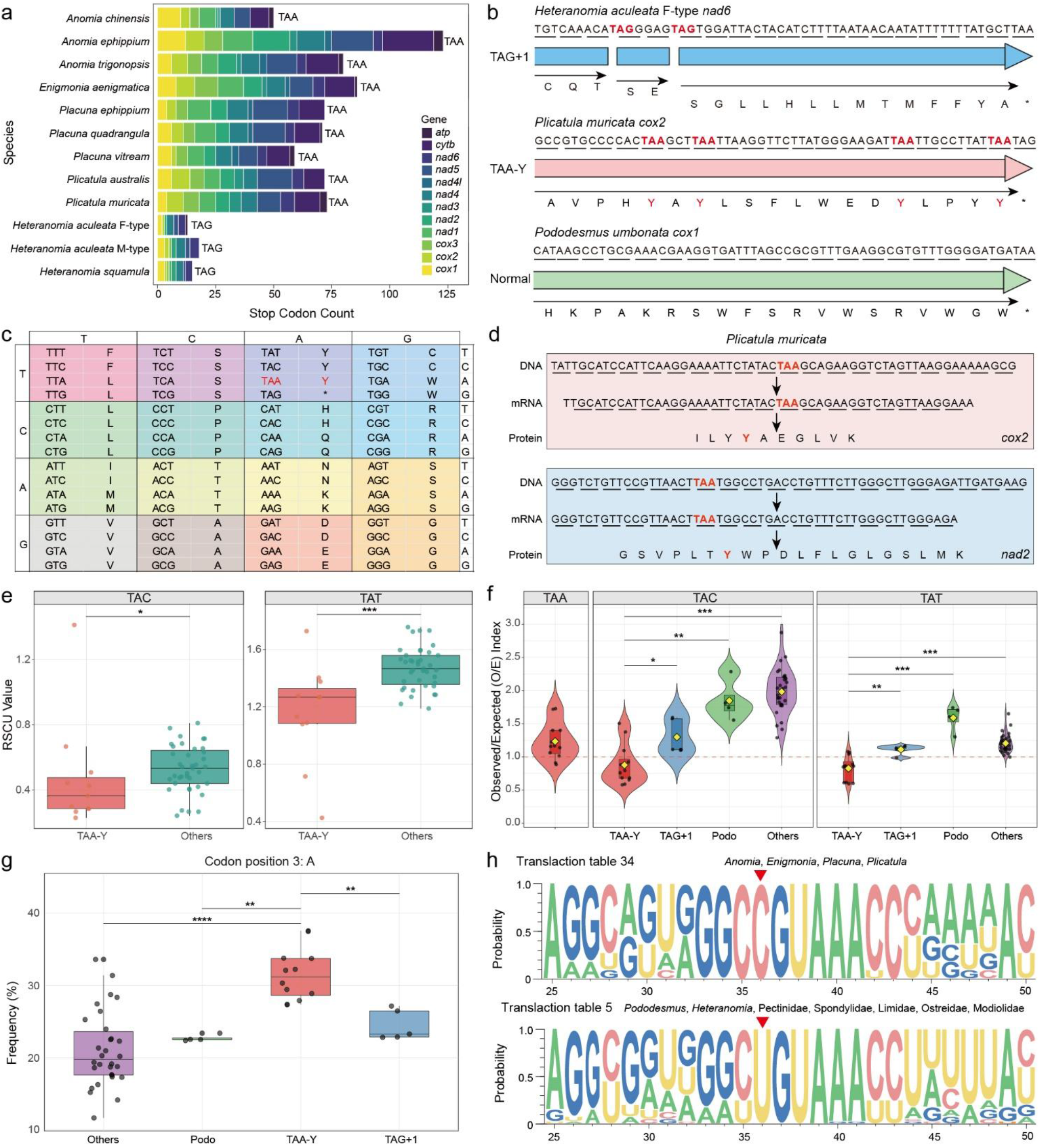
Mitochondrial genetic code variations and evidence for TAA=Tyr reassignment in the APP lineage. (a) Number of in-frame stop codons (TAA or TAG) identified within protein-coding genes (PCGs) of mitochondrial genomes across the examined species. (b) Three distinct mitochondrial genome configurations observed. (c) The newly proposed mitochondrial genetic code (translation table 34) featuring TAA reassigned to Tyr. (d) Genomic, transcriptomic, and proteomic evidence for TAA=Tyr reassignment in *Plicatula muricata*. All peptides detected from mitochondrial proteomics are listed in Table S8. (e) Comparison of relative synonymous codon usage (RSCU) values for TAC and TAA between the TAA=Tyr lineage and other groups. Detailed RSCU results for all examined species are shown in Fig. S9. (f) Observed/expected (O/E) ratios for TAA, TAC, and TAT codons across lineages with different mitochondrial genome configurations. (g) Frequency of adenine (A) at the third codon position in PCGs across lineages with different mitochondrial genome configurations. Base frequencies at all codon positions for these groups are displayed in Fig. S11. (h) Sequence logo of *trnY* (tRNA-Tyr) showing a U36C substitution in the TAA=Tyr lineage compared to other groups. Multiple sequence alignment and predicted secondary structure of *trnY* for all examined species are provided in Fig. S12-14. *: *p* < 0.05, **: *p* < 0.01, ***: *p* < 0.001.

#### Translational frameshifting

Through comparative reading frame analysis, we hypothesized that *Heteranomia* circumvents in-frame TAG codons via a +1 translational frameshift: this mechanism bypasses the thymidine (T) residue of each TAG codon, allowing translation to resume in the next reading frame (Fig. 4b). This strategy is exemplified in the *nad4* and *nad6* genes of *H. aculeata*—each containing four in-frame TAG codons with four corresponding frameshifts— enabling *Heteranomia* to assemble functional respiratory complexes with TAA as the sole termination codon. Programmed +1 translational frameshifting, though well-characterized in viruses, bacteria, ciliates, and yeast via diverse mechanisms including ribosome stalling and translocation-dependent shifts (66–71), is rarely documented in metazoan mitochondria (72, 73). A classic example in animals is the mammalian antizyme gene, where a downstream pseudoknot directs the frameshift (74, 75). The mechanism in *Heteranomia* remains to be determined, but the highly conserved sequence context flanking all TAG codons, a run of adenines upstream and a purine-rich motif downstream, suggests a possible rare codon-induced stalling mechanism (74, 76). However, direct evidence for +1 frameshifting (e.g., ribosome profiling or in vitro translation assays) is currently lacking, and alternative mechanisms such as translational bypassing or stop-codon readthrough cannot be excluded. The invariant AGN codon immediately following each frameshift suggests potential functional significance, but whether this sequence acts as a cis-acting signal or the encoded serine serves a structural role awaits experimental testing. Additionally, TAA remains the sole termination signal, indicating that frameshifting bypasses TAG without interfering with TAA termination, a precise discrimination likely involving ribosome–release factor interactions (72). Observed only in *Heteranomia*, this mechanism may represent either a transient evolutionary state or a stable alternative strategy, with frameshifting rather than codon reassignment reflecting lineage-specific differences in mutation pressure, genetic drift, or translational apparatus architecture.

#### Codon reassignment and tRNA evolution

The reassignment of TAA to encode Tyr represents the major evolutionary trajectory within the APPD lineage (Fig. 4a). In addition to genomic sequence evidence (Fig. S8), transcriptome analysis of *Plicatula muricata* revealed in-frame TAA codons in *cox2* and *nad2* transcripts (Fig. 4d), and mitochondrial proteomics further recovered peptides spanning these TAA-encoding regions (Table S8), providing direct evidence that TAA encodes tyrosine during mitochondrial translation (Fig. 4d). Codon usage patterns lend additional support to this conclusion: RSCU analysis showed significantly reduced TAC/TAT usage in TAA=Tyr lineages compared to other lineages (Fig. 4e & S9), and O/E ratio analysis confirmed that TAA has functionally displaced a portion of TAT/TAC usage (Fig. 4f, Supplementary Information). Furthermore, analysis of nucleotide composition revealed significantly elevated A content at the third codon position in TAA=Tyr lineages, accompanied by a corresponding reduction in G (Fig. 4g & Fig. S10-11, Supplementary Information), suggesting that A bias may be an important contributing factor to the appearance of in-frame TAA codons, thereby creating selective pressure for TAA reassignment. Additionally, codon reassignment is typically associated with changes in tRNA. In all TAA=Tyr lineages, we identified a consistent U36C substitution in the anticodon loop of *trnY* (Fig. 4h & S12–S14), at a position critical for maintaining the structural integrity of the anticodon loop. This substitution is present exclusively in the TAA=Tyr lineages and is absent in all species retaining the canonical code 5 (Fig. S12–S14). It likely weakens the discriminatory stringency of the GUA anticodon against third-position mismatches, thereby enabling trnY to recognize TAA as tyrosine (30, 77). The independent TAA reassignment in the hemichordate Cephalodiscidae also involves a U- to-C substitution at the analogous position in the *trnY* anticodon loop (30). The U36C substitution provides an ambiguous intermediate that allows simultaneous recognition of TAC, TAT, and TAA; if such ambiguous recognition was initially tolerated, selective pressure from abundant TAA codons would have favored its fixation, eventually converting TAA into a genuine tyrosine codon (78). This scenario is consistent with a hybrid model in which mutational pressure (A bias) created the need for reassignment, while a tRNA anticodon loop mutation provided the enabling change (79, 80). Overall, different groups within the APPD lineage have adopted radically different strategies in response to distinct in-frame stop codons: *Heteranomia* evolved a +1 translational frameshift to bypass TAG codons, whereas *Anomia*/*Enigmonia*/*Placuna* and Plicatulidae independently converged on TAA reassignment to tyrosine. The confirmation of TAA reassignment to tyrosine represents the first documented case of mitochondrial genetic code variation within Bivalvia, while the +1 translational frameshifting of TAG in *Heteranomia* represents a complementary translation-level adaptation. These APPD-specific recoding events, however, are not without precedent in the broader tree of life, with similar phenomena having emerged independently across multiple phyla. In the hemichordate Cephalodiscidae, UAA is reassigned to tyrosine (30); in nematode mitochondria, UAA has also been reported to be recoded from a stop codon to tyrosine (29), whereas the earlier claim for platyhelminths (81) has been questioned by subsequent studies (82). In fungi, mitochondrial genetic code reassignments represent an ongoing process occurring in multiple parallel evolutionary steps; Saccharomycotina yeasts, for instance, have undergone CUG codon reassignments in three independent evolutionary events (83). In the mitochondria of Labyrinthulea, UAG is recoded from termination to tyrosine; in certain thraustochytrids, both TAA and TAG are recoded to tyrosine (84). These parallel events across major taxonomic groups, spanning metazoans, fungi, and protists, demonstrate that stop-to-sense codon reassignment is a recurrent convergent strategy in mitochondrial genetic code evolution. This phenomenon underscores the evolutionary plasticity of the genetic code across distantly related lineages, and further highlights the molecular distinctiveness of the APPD lineage.

### Synthesis: An independent ordinal-level lineage

Synthesizing the multiple lines of evidence from phylogeny, karyotype evolution, and mitochondrial genetic code, the APPD lineage exhibits an independent evolutionary trajectory that is distinctly different from other pectinids. Our phylogenomic and morphological analyses consistently recovered the APPD lineage as a distinct clade, separate from the remaining members of pectinids and Limida as ((APPD + (Limida + Pectinoidea)). This topological arrangement, together with the estimated divergence time of approximately 428 MYA—predating or comparable to the ordinal splits between Mytilida and Ostreida and between Venerida and Lucinida—indicates that this lineage has followed an independent evolutionary trajectory for a substantial period. This conclusion is further supported by extensive karyotypic restructuring involving progressive chromosomal fusions from 20 MLGs to as few as 6 chromosomes, and unique mitochondrial features including TAA=Tyr reassignment and +1 translational frameshifting. Collectively, these lines of evidence demonstrate that the APPD lineage has diverged as an independent ordinal-level clade, rendering its current placement within Pectinida *sensu lato* taxonomically unjustified.

Beyond molecular evidence, comparative morphology provides independent support for this distinction. The APPD lineage can be reliably distinguished from other pectinidan families by key shell characters, particularly differences in hinge form, hinge ridge morphology, hinge teeth configuration, and ligament structure. Auricles and the ctenolium are usually absent, and the ligament is diversified—appearing as a single drop shape, inverted V-shape, or divided into inner and outer parts (see Diagnosis below). Although Dimyidae specimens were unavailable for genomic assembly, multiple lines of evidence support their placement within the APPD lineage (details in Supplementary Information). Multi-gene and phylogenomic analyses consistently recover Dimyidae, Plicatulidae, and Anomioidea as a monophyletic clade representing the earliest-diverging lineage within Pectinida *sensu lato*, with Dimyidae as sister to Plicatulidae (32, 35, 36). Morphological examination places Dimyidae between ostreoids and pectinoids but closer to the latter (38), while shell microstructural and conchological evidence refutes a close relationship with Ostreidae (39). Our comparative morphological analysis further demonstrates that Dimyidae shares key characters with Plicatulidae and other APPD members, including a transverse primary ligament overarched by mantle margins, crura-like hinge teeth, absence of auricles, and cementation by the right valve—a combination of features not observed in other pectinidan families. The congruence between molecular phylogenetics, paleontological evidence, and morphological characters supports the placement of Dimyidae within the APPD lineage and justifies its provisional inclusion in the new order Anomiida. Future phylogenomic analyses incorporating Dimyidae will further test the robustness of this taxonomic hypothesis.

### Taxonomy

#### Order

Anomiida ord. nov.

#### Type family

Anomiidae Rafinesque, 1815

#### Constituent families

Anomiidae Rafinesque, 1815 (*Anomia* Linnaeus, 1758; *Enigmonia* Iredale, 1918; *Heteranomia* Winckworth, 1922; *Isomonia* Dautzenberg & Fischer, 1897; *Monia* Gray, 1850; *Patro* Gray, 1850; *Placunanomia* Broderip, 1832; *Pododesmus* Philippi, 1837; *Tedinia* Gray, 1853; fossil: *Carolia* Cantraine, 1838; *Paranomia* Conrad, 1860; *Placunopsis* Morris & Lycett, 1853; *Wakullina* Dall, 1896); Dimyidae Fischer, 1886 (provisional) (*Basiliomya* Bayer, 1971; *Dimya* Rouault, 1850; *Dimyella* Moore, 1970; *Neoatreta* Waller, 2012; fossil: *Atreta* Etallon, 1862; *Diploschiza* Conrad, 1866); Placunidae Rafinesque, 1815 (*Placuna* Lightfoot, 1786); Plicatulidae Gray, 1854 (*Plicatula* Lamarck, 1801; fossil: *Eoplicatula* Carter, 1990; *Harpax* Parkinson, 1811; *Pseudoplacunopsis* Bittner, 1895).

#### Etymology

Derived from the type family Anomiidae.

#### Distribution

Marine habitats from intertidal to shelf depths.

#### Diagnosis

Bivalves usually sessile, without auricle and ctenolium; hinge short or degraded, with hinge ridge and denticulate teeth present in most lineages but absent in Anomiidae; ligament diversified: single drop-shaped (Plicatulidae), inverted V-shaped (Placunidae), or divided into inner and outer parts (Dimyidae) (85, 86), versus consistently alivincular, drop-shaped to triangular resilium in Pectinida *sensu stricto* and Limida; pallial line generally clear, adductor muscle scars one to three (versus strictly monomyarian in Pectinida *sensu stricto* and Limida); foot weak or completely degraded (except in secondarily motile *Enigmonia*); pallial eyes usually reduced or absent.

### Remarks

The order Anomiida ord. nov. (APPD lineage) has traditionally been placed within Pectinida *sensu lato* and considered closely related to scallops (87). Recent molecular phylogenomic analyses provide robust support for the separation of the APPD lineage from other pectinids. Specifically, these families form a well-supported clade that is sister to the other families, including Propeamussiidae, Pectinidae, Spondylidae, and Entoliidae (31, 32, 35, 36). However, Anomiida exhibits significant disparities in lifestyle compared to typical members of Pectinida *sensu stricto*. The latter encompasses a diverse range of habits: while many families (e.g., Pectinidae, Propeamussiidae, Entoliidae) are free-living or byssally attached, with numerous species capable of swimming, Spondylidae cement to hard substrates by their right valve (88–90). In contrast, Anomiida are predominantly sessile, with most members cementing their right valve to hard substrates (85, 90). Exceptions include the anomiid *Enigmonia aenigmatica*, with the ability to crawl on mangrove leaves and stems (representing a secondary reacquisition of mobility), and species of *Placuna*, which lose their byssal apparatus in adulthood and adopt a free-living yet immobile existence on soft substrates (90, 91).

Morphologically, Anomiida differ substantially from typical scallops in overall shell shape, lacking the characteristic fan-shaped form with prominent auricles (92). They can be reliably distinguished from Pectinida *sensu stricto* and Limida by fundamental differences in hinge architecture, ligament configuration, and adductor muscle scar patterns (85, 90). Auricles and the ctenolium are usually absent, and the hinge is typically short or degraded, though a hinge ridge and denticulate teeth (composed of aragonite) are present in most groups, including plicatulids, placunids, and dimyids (85, 93). The ligament is diversified, appearing as a drop-shaped structure, inverted V-shape, or divided into discrete inner and outer parts (90). The pallial line is generally clear, and the number of adductor muscle scars varies from one to three, reflecting differences in muscular organization (32, 85). The foot is weak or completely degraded in adults, correlating with sessile habits, except in the motile *Enigmonia* (90). In contrast, most members of Pectinida *sensu stricto* and Limida exhibit a more conservative, fan-shaped shell, characterized by a long, linear hinge and prominent auricles distinctly separated from the shell disc (92). Furthermore, the pectinid ligament (resilium) is consistently alivincular, with a well-defined, drop-shaped to triangular inner ligament layer (89, 94). A byssal notch is present in the right valve of many species, serving as an opening for the byssus during early ontogeny (85, 89, 95), and the monomyarian adductor scar is invariably single (96). Moreover, the ctenidial structure in Pectinida *sensu stricto* is typically plicate with well-developed interlamellar junctions, and many pectinids possess elaborate pallial eyes on the mantle margins; these features are reduced or absent in most Anomiida (89, 97–99). It should be emphasized that the morphological diagnosis rests on a combination of characters rather than any single autapomorphy. For example, while reduced auricles occur sporadically in some Pectinidae (e.g., Propeamussiidae), they co-occur there with a well-developed byssal notch and a strictly alivincular ligament, the features not found in Anomiida. Similarly, the loss of a ctenolium is observed in other pteriomorphians, but never in conjunction with the diverse ligament configurations and variable adductor muscle scar numbers documented here. This character combination provides a robust morphological basis for distinguishing Anomiida from all other bivalve orders.

Within Anomiida ord. nov., Anomiidae exhibits a short or degraded hinge with absent hinge teeth and a variably calcified byssus; a definitive feature is the presence of a central byssal notch in the right valve (90). However, in some species such as *Placunanomia*, the byssal notch can become sealed by the calcified byssal plug (90). Placunidae possess a reduced hinge structure, but distinct hinge teeth are evident in some species; more importantly, placunids exhibit a characteristic inverted V-shaped ridge and a primary ligament that provides the opening thrust, complemented by a secondary periostracal ligament that maintains valve alignment (90, 100). This secondary ligament formation is a key innovation enabling the free-living yet immobile existence of placunids on soft substrates (91). Additionally, both Plicatulidae and Dimyidae are cemented by the right valve and retain strong, crura-like hinge teeth that articulate with corresponding sockets (39, 89, 101). However, the ligament in Plicatulidae is profoundly different, with an internal, transversely compressed, hoop-like structure overarched by the mantle margins (89, 101). In Dimyidae, the alivincular ligament is situated on a more prominent, often calcitic, hinge plate and is accompanied by a secondary periostracal ligament (85, 86, 101). Additionally, the retention of an anterior adductor muscle in Dimyidae, a primitive feature lost in other monomyarians, underscores their early divergence within Pteriomorphia (86, 89, 101). Nevertheless, further morphological and molecular sampling of additional anomiid genera and critical fossil taxa will be essential to test the stability of this emerging classification and fully resolve the evolutionary history of this lineage.

### Limitations of this study

The classification of the FC and VCs on *P. vitream* chromosome 7 as functional or vestigial is based on sequence compositional criteria; in situ functional assays (e.g., CENH3 localization) will be essential to validate these assignments and to resolve the evolutionary origins of the FC. Despite extensive efforts to collect representative specimens, we failed to acquire Dimyidae specimens due to their rarity. While we provisionally place Dimyidae within Anomiida based on morphological affinities and previous phylogenetic studies, molecular validation, including mitochondrial and nuclear genomic analyses, is required to confirm this placement and to determine whether Dimyidae shares the TAA=Tyr genetic code and chromosomal fusions documented in other anomiid families. The precise mechanism underlying the +1 translational frameshift in *Heteranomia* also remains uncharacterized, as only dried specimens were available, precluding transcriptomic or functional assays. Elucidating this recoding mechanism will ultimately require fresh material and approaches such as ribosome profiling or heterologous expression. Similarly, experimental validation of the TAA=Tyr reassignment mechanism awaits suitable models. Although the correlation between the U36C substitution and TAA reassignment is compelling, definitive demonstration of causality will require experimental validation. Specifically, in vitro aminoacylation assays using purified wild-type and mutant tRNA-Tyr variants are further required to ideally test their ability to recognize TAA codons; alternatively, a heterologous expression system expressing the mutant tRNA could assess whether TAA is translated as tyrosine in vivo. Additionally, the proposed ‘ambiguous intermediate’ model, in which the U36C substitution allows simultaneous recognition of TAC, TAT, and TAA before the fixation of reassignment, remains theoretical and would benefit from evolutionary simulations or biochemical binding assays to determine the relative affinities of the mutant tRNA for each cognate codon. Such experiments are beyond the scope of the present study but represent high priorities for future investigation.

## CONCLUSION

The APPD lineage, comprising Anomiidae, Placunidae, Plicatulidae, and, by morphological inference, Dimyidae, represents an extraordinary case of genomic innovation that expands our understanding of the plasticity of bivalve genomes. Through an integrative approach combining phylogenomics, comparative genomics, and proteomics, we have uncovered three distinct configurations of mitochondrial genetic code evolution, documented a rare +1 translational frameshift mechanism, and revealed a pattern of progressive chromosomal fusion accompanied by the preservation of vestigial centromeres. We therefore propose the new order Anomiida ord. nov. These findings collectively resolve long-standing systematic uncertainties within Pectinida *sensu lato* and provide compelling evidence that this lineage constitutes a deeply divergent clade warranting ordinal-level recognition. Beyond the taxonomic significance, this work establishes the APPD lineage as a valuable model for investigating fundamental principles of genome evolution, ranging from codon reassignment and translational recoding to karyotype restructuring and centromere dynamics. The combination of multiple exceptional genomic features within a single lineage offers unique opportunities for future functional studies and expands our understanding of the remarkable plasticity of molecular evolution across the tree of life.

## DATA AVAILABILITY

All raw sequencing data generated during this study, including genomic Illumina and PacBio HiFi reads, transcriptome sequences, and nuclear genome assemblies, have been deposited in the NCBI Sequence Read Archive (SRA) under the BioProject accession number PRJNA1198682. The functional annotations of the assemblies and mitochondrial genomes are available in Figshare under the DOI: 10.6084/m9.figshare.33250323. The mass spectrometry proteomics data (LS/MS) have been deposited in the ProteomeXchange Consortium via the PRIDE partner repository with the dataset identifier [PXD identifier pending review].

## ACKNOWLEDGMENTS AND FUNDING SOURCE

This study was supported by Research Grants Council of Hong Kong (General Research Fund: 12101021 and 12102222), the Collaborative Research Fund (C2013-22GF), the Hainan Province Science and Technology Special Fund (SOLZSKY2026011), the Hainan Province Science and Technology Talent Innovation Project (KJRC2023A02), National Natural Science Foundation of China (32303034, 32273118), and Shenzhen Science and Technology Program (CJGJZD2025090291534011). We thank Mr. Siwei Liu (Littoral ENvironnement et Sociétés), Mr. Haodong Liu (Qingdao Shell Museum), Mr. Xing-Xiao Wang (Capital Normal University), Dr. Yanjie Zhang (Hainan University), and Mr. Ge Guo (Beijing Renchao Fen Dongba School) for their assistance in sample collection.

## AUTHOR CONTRIBUTIONS

JWQ conceived and designed the project. YTL, YXL, and MT collected the samples. YTL, YXL, and XYL conducted experiments. YTL and YXL performed data analyses. ZH, JH, and ZB provided critical comments. YTL drafted the manuscript. All authors contributed to the revision and approved the submitted version.

## DECLARATION OF INTERESTS

The authors declare no competing interests.

## METHOD DETAILS

### Sample collection and treatment

Details of specimen collection are provided in Table S1. For mitogenomic sequencing, specimens that had been dry-preserved at room temperature were dissected; the dry tissues were softened and then preserved in 70% ethanol. Mantle tissues from fresh specimens were directly preserved in 100% ethanol for DNA extraction. Additionally, several individuals of *Anomia chinensis*, *Placuna vitream*, and *Plicatula muricata* were dissected to obtain tissues for DNA, RNA, and protein extraction for genomic assembly and RNA sequencing.

### DNA extraction and genomic sequencing

Specimens with air-dried tissues were softened using 50% ethanol. Genomic DNA was extracted from the soft tissues using the CTAB method (102). DNA quality was examined using agarose gel (1.0%) electrophoresis, and the quantity was determined using a NanoDrop ND-1000 spectrophotometer (Thermo Scientific, USA). The DNA library was constructed with the NEBNext Ultra DNA Library Prep Kit for Illumina (NEB, United States) with an insert size of 350 bp. Genomic sequencing was conducted on an Illumina NovaSeq 6000 sequencer (Illumina, USA) in Novogene (Tianjin, China) to generate 150-bp paired-end reads. PacBio HiFi sequencing was used to generate draft genome assemblies. The DNA libraries of the mantle tissues of *Anomia chinensis*, *P. vitream*, and *P. muricata*, respectively, were constructed using the SMRTbell Express Template Prep Kit 2.0, following the manufacturer’s protocol. Each library was sequenced for one SMRT Cell sequencing run on a PacBio Revio sequencing plate in Novogene (Tianjin, China). High-throughput chromosome conformation capture (Hi-C) sequencing was used to scaffold the draft assemblies to chromosomal-level genomes. The gill tissues and shells were removed from the specimens of *A. chinensis*, *P. vitream*, and *P. muricata*, then the remnants were used for library construction with an insert size of 350 bp using the TruSeq Nano DNA HT Sample Preparation Kit (Illumina, USA) following the manufacturer’s protocol. Then, the libraries were sequenced on an Illumina NovaSeq 6000 sequencer (Illumina, USA) to produce 150 bp paired-end reads in Novogene (Tianjin, China).

### RNA library construction and sequencing

The total RNA of each tissue, including adductor muscle, digestive gland, foot, gill, mantle, and gonad from three individuals of *A. chinensis*, *P. vitream*, and *P. muricata*, was extracted using the Trizol reagent (TAKARA, Japan). The RNA quality and quantity were examined as described for DNA samples above. Then, the ribosomal RNA was removed from the total RNA using a NEBNext Ultra RNA Library Prep Kit for Illumina (NEB, USA), and the RNA molecules were fragmented into 250-300 bp and reverse-transcribed into cDNA. The constructed libraries were paired-end sequenced on an Illumina NovaSeq 6000 sequencer (Illumina, USA) to produce 150 bp paired reads.

### Mitogenome assembly, annotation, and genetic codon prediction

Raw Illumina reads were filtered using Trimmomatic v0.39 (103) to remove the adapter and low-quality reads with the following settings: LEADING = 15, TRAILING = 15, SLIDINGWINDOW = 4:20, MINLEN = 40. We used two approaches to assemble the mitogenomes to avoid assembly errors, including SPAdes v3.14.1 (104) and NOVOPlasty v3.2 (105). In detail, the clean reads were *de novo* assembled using SPAdes with *k-*mer sizes 21, 33, 55, 77, 99, and 127. The mitogenomes were identified by conducting BLASTn using BLAST v2.11.0+ (106) to search against the mitochondrial genomes or gene fragments of available phylogenetically close species from NCBI (https://www.ncbi.nlm.nih.gov/), with an E-value of 1e-10. In addition, the clean reads were assembled using NOVOPlasty with the mitogenomes of available phylogenetically close species as the templates and their *cox1* sequences as the seeds. After that, MITOS2 in the Galaxy online server (https://usegalaxy.eu/) was used to annotate the mitogenome with the invertebrate mitogenomic codes. The annotated PCGs were manually examined and adjusted against the invertebrate mitogenomic code or the new one proposed by this study. Additionally, the OGDRAW in the Geseq server (107) was used to visualize the mitogenomic structure and genetic arrangement. The genetic codes of the mitogenomes in this study were predicted using Codetta v2.0 (108) with the following parameters: -m -e 1e-5 -r 0.7.

### Mitochondrial genome comparative analyses

#### Base composition and AT skew

Complete mitochondrial genome sequences and concatenated protein-coding gene (PCG) sequences were extracted from each assembly. Nucleotide composition (A, T, G, and C) was calculated using custom Python scripts based on Biopython (109). AT content was defined as (A+T)% and AT skew as (A−T)/(A+T). Differences among translation-mode groups (normal, TAA=Tyr, frameshift) were tested using the Kruskal-Wallis test, followed by pairwise Mann-Whitney U tests with Bonferroni correction.

#### Codon usage and relative synonymous codon usage (RSCU)

PCG sequences were translated according to the respective translation mode (code 5, TAA=Tyr reassignment, or +1 frameshift). Codon counts were obtained, and RSCU was calculated for each codon as the observed frequency divided by the expected frequency under equal usage of synonymous codons. An RSCU value of 1 indicates no bias, > 1 indicates preference, and < 1 indicates under-representation.

#### Observed/expected (O/E) index for tyrosine codons

The observed/expected (O/E) ratio was calculated to quantify the usage bias of tyrosine codons relative to the neutral expectation based on nucleotide composition (110). For each mitochondrial genome, the observed count (O) of each tyrosine codon (TAT, TAC, and TAA in TAA=Tyr lineages) was obtained from codon usage tables. The expected count (E) was calculated assuming independent base frequencies at each codon position: E=Fp1×Fp2×Fp3×Tc, where Fp1, Fp2, and Fp3 are the proportion of frequencies (0–1) of the corresponding nucleotides at the first, second, and third codon positions, and Tc is the total frequency of occurrence of codons. Pairwise comparisons of O/E ratios for TAT and TAC were performed between the TAA=Tyr group and each of the other three groups (TAG+1, *Pododesmus*, and other code 5). Normality was assessed with the Shapiro-Wilk test and variance homogeneity with Levene’s test. Depending on the results, either a two-sample t-test (with equal or unequal variance) or the Wilcoxon rank-sum test was used. Significance levels were set at *p* < 0.05, with additional levels at *p* < 0.01 and *p* < 0.001; *p* < 0.10 was considered a trend.

#### Codon-position nucleotide composition

The frequencies of A, T, C and G were tallied separately for each position in each codon of each PCG. The percentage of each base at each position was then calculated per species. Between-group differences were tested using Kruskal–Wallis tests followed by pairwise Wilcoxon tests with Bonferroni adjustment. Results were visualized as boxplots overlaid with jittered data points; significance indicators used asterisks (*: *p* < 0.05, **: *p* < 0.01, ***: *p* < 0.001) and “ns” for non-significant.

#### The *trnY* sequence alignment and logo diagrams

All *tRNA-Tyr* (*trnY*) gene sequences were extracted from the annotated mitogenomes and aligned using MAFFT v7.520 (111) with default parameters. The alignment was submitted to the WebLogo online server (https://weblogo.threeplusone.com/) to generate a sequence logo showing base conservation and information content at each position under default settings.

#### Gene order comparison

All genes (PCGs, tRNAs, rRNAs) were extracted from each mitochondrial assembly, sorted by their start position on the mitochondrial DNA, and concatenated into a linear gene order list. To quantify similarity between gene orders, a gene order similarity index (GOSI) was calculated as the length of the longest common subsequence of shared genes divided by the total number of genes in the smaller genome.

#### Protein sequence alignment

The amino acid sequences of *cox1* were extracted for all species and aligned using MAFFT v7.520 (111) with the L-INS-i strategy, followed by manual refinement. The alignment was used for subsequent phylogenetic reconstruction.

#### Mitogenomic Phylogeny

A concatenated dataset comprising all 13 PCGs and the two ribosomal RNA genes (*12S rRNA* and *16S rRNA*) was assembled from 57 mitochondrial genomes, with *Anadara broughtonii* from the order Arcida as the outgroup. Phylogenetic analyses were conducted using PhyloSuite v1.2.2 (112) with the following plug-in programs. Each gene was aligned individually with MAFFT v7.520 (111) under the “auto” option and the “Normal alignment” mode. Ambiguously aligned regions were removed using GBlocks v0.91b (113), and missing genes or alignment gaps were filled with “-”. ModelFinder (114) was used to select the best-fit partition model (Edge-unlinked) using the BIC criterion. Best-fit model according to BIC: mtInv+F+R5 for PCGs and Blosum62+F+I+G4 for rRNAs. Maximum likelihood phylogenies were inferred using IQ-TREE (115) under an Edge-linked partition model for 10000 ultrafast (116) bootstraps, as well as the Shimodaira-Hasegawa-like approximate likelihood-ratio test (117).

### Mitochondrial Proteome

To verify the reassignment of the TAA codon on the protein level, we performed mitochondrial proteomics using an individual of *P. muricata*. Mitochondria were isolated from the sample using the Mitochondria Isolation Kit for mammalian cells (Thermo Fisher, USA) according to the manufacturer’s protocol. Proteins were extracted with SDT lysis buffer (4% SDS, 100 mM Tris-HCl, pH 7.6), sonicated on ice, boiled for 15 min, and centrifuged at 14,000g for 15 min to remove debris. Protein concentration was determined using the BCA assay (Takara, Japan). Quality control was performed by separating 10 µg of each protein sample on a 12% SDS-PAGE gel, followed by Coomassie Blue staining. For in-solution digestion, protein samples were reduced with 40 mM TCEP and alkylated with 100 mM iodoacetamide. After boiling for 5 min, 250 mM hydroxypropyl-β-cyclodextrin and 100 mM ammonium bicarbonate were added, followed by overnight digestion with trypsin (Promega, China) at an enzyme-to-substrate ratio of 1:50 (37°C). Digestion was stopped by acidifying to pH < 2 with trifluoroacetic acid. The resulting peptides were desalted on C18 cartridges, dried in a vacuum concentrator, and reconstituted in 15 µL of 0.1% formic acid. Peptide quantification was performed using a NanoDrop ND-1000 spectrophotometer (Thermo Scientific, USA). Peptides were separated on a NanoElute chromatography system (Bruker, Germany) coupled to a timsTOF Pro mass spectrometer (Bruker, Germany) operated in PASEF mode. Separation was carried out on a 25 cm × 75 µm analytical column packed with 1.6 µm C18 beads (IonOpticks, Australia) at a flow rate of 300 nL/min, using a 60 min linear gradient of buffer B (0.1% formic acid in 100% acetonitrile). MS data were acquired over a mass range of 100–1700 m/z with an ion mobility range of 0.75–1.4 V·s/cm². PASEF settings included 10 MS/MS scans per cycle (total cycle time 1.16 s), an intensity threshold of 2,500, and collision energies of 20–59 eV.

Raw MS data were processed using MaxQuant v1.6.17.0 (118). A species-specific protein database was constructed from the *P. muricata* mitochondrial and nuclear genomes. Search parameters included trypsin/P with up to two missed cleavages; precursor mass tolerance of 10 ppm (main search) and 20 ppm (first search); fragment mass tolerance of 20 ppm. Carbamidomethylation of cysteine was set as a fixed modification; oxidation of methionine and N-terminal acetylation were set as variable modifications. Both peptide and protein false discovery rates (FDR) were set to ≤ 0.01. Label-free quantification (LFQ) was enabled to estimate protein abundance. To confirm TAA=Tyr reassignment, extracted ion chromatograms and MS/MS spectra of peptides containing in-frame TAA codons were manually inspected for the presence of tyrosine residues at the corresponding positions. All peptides reported were matched with 100% identities and were derived from mitochondrial-encoded proteins.

### Morphological phylogenetic analysis

A morphological matrix was assembled for 43 terminal taxa representing the major pteriomorphian lineages, including Ostreida, Mytilida, Limida, Pectinidae, Propeamussiidae, Spondylidae, Anomiidae, Placunidae, Plicatulidae, and Dimyidae. A mytiloid species (*Idas* sp.) was designated as the outgroup. The matrix comprised 21 discrete shell characters (Table S3), which were coded based on direct observation of voucher specimens and, for Dimyidae, from published descriptions and illustrations (85). All characters were treated as unordered and equally weighted. The analysis was conducted using the phangorn package (119). Heuristic searches were performed with the ratchet function (implementing the ratchet method) with 100000 iterations, 1000 random starting trees, and subtree pruning and regrafting (SPR) rearrangements (120, 121). Nodal support was evaluated by 100000 bootstrap pseudoreplicates using the bootstrap.phyDat function, with the same tree-search settings (122). The resulting tree was rooted with *Idas* sp. and visualized using ggtree (123).

### Genome Estimation and Assembly

The Illumina raw reads for each species were trimmed using Trimmomatic v0.39 (103) (quality score < 30, length < 40 bp). The clean data were used to generate a 21 *k*-mer histogram using Jellyfish v2.2.028 (124), and GenomeScope v2.0 (125) was used to estimate the genome sizes of *A. chinensis*, *P. vitream*, and *P. muricata*.

The *de novo* assembly of HiFi reads to generate draft genomes was performed using HiFiasm v0.18.9 (126). Possible alternative heterozygous contigs were eliminated from the draft genomes using Purge_Haplotigs v1.1.3 (127) with the parameters of l = 5, m = 70, and h = 180 based on the read-depth histogram generated by the pipeline. Then, the de-redundant draft genomes were assessed for assembly statistics using QUAST v5.2.0 (128) under the default settings. The completeness of the de-redundant draft genomes (scaffold-level assembly) was assessed by analyzing the Compleasm v0.2.6 (129) scores against the database Metazoa_odb10 under the default settings.

The raw reads of Hi-C sequencing were trimmed using Trimmomatic v0.39 (103) (quality score < 30, length < 40 bp), followed by identification of high-quality reads using HiC-Pro v2.10 (130) and removal of duplicated reads using the Juicer pipeline v1.5 (131) under the default settings. The genomic scaffolding was performed using the 3D *de novo* assembly pipeline (132) under the haploid genome model. The pseudo-chromosomal linkage groups were checked, and corrections were made in Juicebox v1.11.08 (133) to ensure that the scaffolds within the same pseudo-chromosomal linkage groups met the Hi-C linkage characteristics. The completeness of the final genome assemblies (chromosome-level) was assessed by using Compleasm v0.2.6 (129) scores against the database Metazoa_odb10 under the default settings. In addition, QUAST v5.2.0 (128) was used to assess the assembly statistics under the default settings.

### Gene model prediction and functional annotation

The final versions of the assemblies were soft masked using RepeatMasker v4.1.2 (http://www.repeatmasker.org/) against the repeat libraries of all model organisms in RepBase v20181026 (134) and the species-specific repeat libraries constructed using RepeatModeler v2.0.3 (135) under the default settings. MAKER v3.0 (136) was applied to predict gene models. *De novo* and genome-guided transcriptomes of the tissues were assembled to provide transcriptomic evidence. Adapters and low-quality reads (quality score < 20, length < 40 bp) of the RNA sequencing data from the adductor muscle, digestive gland, foot, gill, mantle, and gonad tissues of the three species were removed using Trimmomatic v0.39 (103) (quality score < 30, length < 40 bp). All clean reads of the six tissues were pooled and used for *de novo* assembly using Trinity v2.8.5 (137) under the default settings. Genome-guided assembly was performed with Trinity by aligning RNA sequencing data to the genome using HISAT v2.1.0 (138) under the default settings. These two transcriptomes were merged using the PASA pipeline v2.2.0 (139) following the authors’ instructions. Selected molluscan protein sequences (Table S2) were used as protein evidence. Augustus v3.1 (140) was used for the gene *de novo* prediction in the repeat-masked genome sequences. EVidenceModeler v1.1.1 (141) was used to integrate results from different gene predictors, including Augustus and Maker. The PASA pipeline v2.2.0 (139) was used to improve the EVM gene models by modifying gene structures and adding UTR annotations with the *de novo* assembled transcriptome. The gene models were then filtered using gFACs v1.1.2 (142) with the following parameters: min-exon-size 20, min-intron-size 20, min-CDS-size 150, rem-5prime-3prime-incompletes, rem-all-incompletes, unique-genes-only, and allowed-inframe-stop-codons 0. Then, the completeness of the final predicted gene models was assessed by using Compleasm v0.2.6 (129) against the Metazoa_odb10 database under the protein model. The predicted genes were functionally annotated using EggNOG-MAPPER v5.0 (143) under the default settings and the BLASTP model of Diamond v0.9.24 (144) targeting NR databases (accessed March 2025) with an E-value of 1e-10.

### Phylogenomics and divergent time estimation

Orthologous groups (OGs) among the selected genomes, including one annelid, one brachiopod, and 27 molluscs (Table S2 & S9), were inferred using Diamond v2.0.15 BLASTp implemented in OrthoFinder v2.5.5 (145), with the “more-sensitive” option and an E-value threshold of 1e-10. Only single-copy OGs with at least 80% taxonomic representation (22 species) in OGs, including 714 single-copy OGs, were used to construct the phylogenetic tree. The protein sequences were aligned using MAFFT v7.520 (111) under the “auto” strategy to align the single-copy OGs in the “Normal alignment mode”. Gblocks v0.91b (113) was applied to remove ambiguously aligned fragments in batches with missing protein or alignment gaps filled with “-”. The aligned sequences with missing sequences were concatenated. The maximum-likelihood (ML) analysis was conducted using IQ-TREE v2.1.3 (115) under the MFP option for model selection and then run for 1,000,000 ultrafast bootstraps. The divergence time estimation was conducted based on the ML tree constructed above using MCMCtree implemented in PAML v4.9h (146). Available fossil records or geological events in MCMCtree were used to constrain the corresponding nodes (146) (Table S4). The LG model was employed for each partition. The burn-in, sample frequency, and number of samples were set at 1 million, 1000, and 10000, respectively, and MCMC was run for 10 million generations.

### Macro- and micro-synteny analysis

To reconstruct the karyotype evolution of the focal species, we performed macrosynteny comparisons against the MLGs (41). Genome assemblies from this study and the selected bivalve species were analyzed using the PanSyn pipeline (147) with default parameters. Ancestral chromosome states were inferred by mapping the observed synteny blocks onto a phylogeny, and the fusion history was deduced by manually reverse inference from the karyotypes of extant species relative to the MLGs. Microsynteny analysis was carried out with MCScanX (148) under default settings, and the resulting synteny networks were visualized using NGenomeSyn v1.41 (149).

### Centromere identification and characterization

Putative centromeric regions were identified using RepeatOBserver V1 (150) under default settings to locate regions with low sequence complexity and high repeat density, indicative of centromeres. For *P. vitream* chr7, three regions were identified: a functional centromere (FC) and two vestigial centromeres (VC1 and VC2). The three loci were extracted as FASTA files using custom R scripts. Pairwise dot plots between FC, VC1, and VC2 were generated using D-Genies (151). A sliding window of 35 bp (step = 35 bp) and a perfect match threshold of 35 bp were applied to visualize sequence similarities. GC content along each region was calculated with a sliding window of 100 bp (step = 100 bp) using custom R scripts based on the Biostrings package (152).

## Notes

### Competing Interest Statement

The authors have declared no competing interest.

## REFERENCES

1. T. D. Lewin, I. J.-Y. Liao, Y.-J. Luo, Annelid comparative genomics and the evolution of massive lineage-specific genome rearrangement in bilaterians. Mol. Biol. Evol. 41, msae172 (2024).

2. K. Lucek, et al., The impact of chromosomal rearrangements in speciation: From micro- to macroevolution. Cold Spring Harb. Perspect. Biol. 15, a041447 (2023).

3. W. Han, et al., Ancient homomorphy of molluscan sex chromosomes sustained by reversible sex-biased genes and sex determiner translocation. *Nat*. Ecol. Evol. 6, 1891–1906 (2022).

4. S. Wang, et al., Scallop genome provides insights into evolution of bilaterian karyotype and development. *Nat*. Ecol. Evol. 1, 0120 (2017).

5. Y.-T. Lin, et al., Glass scallop genome reveals key adaptations to deep-sea environments and ectosymbiosis. Nat. Commun. 17, 4713 (2026).

6. M. G. Corni, M. Trentini, A chromosomic study of *Chamelea gallina* (L.) (Bivalvia, Veneridae). Ital. J. Zool. 53, 23–24 (1986).

7. H. Chen, et al., A new genus and two new species of freshwater mussels (Bivalvia, Unionidae) from Sichuan, China: Overlooked cryptic endemism in the upper Yangtze River Basin. Zoosystematics Evol. 101, 1459–1470 (2025).

8. H. Ieyama, Chromosome numbers of ten species in four families of Pteriomorphia (Bivalvia). Venus J. Malacol. Soc. Jpn. 33, 129–137 (2018).

9. A. N. M. Z. Iqbal, M. S. Khan, M. A. Navalgund, U. Goswami, Physical mapping of *18S rRNA* gene in green mussel *Perna viridis* – An indication of higher major rRNA gene clusters. *Russ*. J. Mar. Biol. 48, 195–201 (2022).

10. X. Li, et al., OysterDB: A genome database for Ostreidae. Mar. Biotechnol. 26, 827–834 (2024).

11. Q. Mu, et al., The chromosome-level genome assembly and annotation of the silver-lipped pearl oyster, *Pinctada maxima*. Sci. Data 12, 1301 (2025).

12. C. Zhou, et al., Chromosome-level genome assembly of the deep-sea solemyid bivalve *Acharax haimaensis*. Sci. Data 13, 559 (2026).

13. J. M. Serb, A. Alejandrino, E. Otárola-Castillo, D. C. Adams, Morphological convergence of shell shape in distantly related scallop species (Mollusca: Pectinidae). Zool. J. Linn. Soc. 163, 571–584 (2011).

14. P. M. Mikkelsen, R. Bieler, I. Kappner, T. A. Rawlings, Phylogeny of Veneroidea (Mollusca: Bivalvia) based on morphology and molecules. Zool. J. Linn. Soc. 148, 439–521 (2006).

15. N. I. Selin, E. E. Vekhova, Morphology of the bivalve mollusks *Crenomytilus grayanus* and *Mytilus coruscus* in relation to their spatial distribution in the upper subtidal zone. *Russ*. J. Mar. Biol. 28, 213–218 (2002).

16. T. V. Goto, H. B. Tamate, N. Hanzawa, Phylogenetic characterization of three morphs of mussels (Bivalvia, Mytilidae) inhabiting isolated marine environments in Palau Islands. Zoolog. Sci. 28, 568–579 (2011).

17. T. Malkócs, et al., Complex mitogenomic rearrangements within the Pectinidae (Mollusca: Bivalvia). BMC Ecol. Evol. 22, 29 (2022).

18. F. Plazzi, A. Formaggioni, M. Passamonti, Mito-nuclear coevolution and phylogenetic artifacts: The case of bivalve mollusks. Sci. Rep. 12, 8395 (2022).

19. C. Saccone, et al., Mitochondrial DNA in metazoa: Degree of freedom in a frozen event. Gene 286, 3–12 (2002).

20. F. Ghiselli, et al., Molluscan mitochondrial genomes break the rules. Philos. Trans. R. Soc. B Biol. Sci. 376, 20200159 (2021).

21. F. Plazzi, G. Puccio, M. Passamonti, Comparative large-scale mitogenomics evidences clade- specific evolutionary trends in mitochondrial DNAs of Bivalvia. Genome Biol. Evol. 8, 2544– 2564 (2016).

22. D. Guerra, et al., Evolution of sex-dependent mtDNA transmission in freshwater mussels (Bivalvia: Unionida). Mol. Phylogenet. Evol. 115, 605–608 (2017).

23. R. D. Knight, S. J. Freeland, L. F. Landweber, Rewiring the keyboard: Evolvability of the genetic code. Nat. Rev. Genet. 2, 49–58 (2001).

24. K. Watanabe, Unique features of animal mitochondrial translation systems: The non-universal genetic code, unusual features of the translational apparatus and their relevance to human mitochondrial diseases. Proc. Jpn. Acad. Ser. B 86, 11–39 (2010).

25. J. L. Boore, W. M. Brown, Complete DNA sequence of the mitochondrial genome of the black chiton, *Katharina tunicata*. Genetics 138, 423–443 (1994).

26. G. Wang, T. Xue, M. Chen, L. Guo, J. Li, Complete F-type mitochondrial genome of freshwater mussels *Unio douglasiae*. Mitochondrial DNA Part A 27, 4021–4022 (2016).

27. X. Shen, et al., The first mitochondrial genome of *Coelomactra antiquata* (Mollusca: Veneroida: Mactridae) from Guangxi (China) and potential molecular markers. Mitochondrial DNA Part A 27, 3642–3643 (2016).

28. Y. Shulgina, S. R. Eddy, A computational screen for alternative genetic codes in over 250,000 genomes. eLife 10, e71402 (2021).

29. J. E. Jacob, B. Vanholme, T. Van Leeuwen, G. Gheysen, A unique genetic code change in the mitochondrial genome of the parasitic nematode *Radopholus similis*. BMC Res. Notes 2, 192 (2009).

30. Y. Li, et al., Mitogenomics reveals a novel genetic code in Hemichordata. Genome Biol. Evol. 11, 29–40 (2019).

31. R. Bieler, et al., Investigating the Bivalve Tree of Life – an exemplar-based approach combining molecular and novel morphological characters. Invertebr. Syst. 28, 32–115 (2014).

32. Y. T. Lin, J. W. Qiu, Taxonomic revision of jingle shells: Resurrecting and reclassifying species of Anomiidae (Bivalvia: Pectinida). Ecol. Evol. 15, e72372 (2025).

33. D. J. Combosch, et al., A family-level tree of life for bivalves based on a Sanger-sequencing approach. Mol. Phylogenet. Evol. 107, 191–208 (2017).

34. S. Lemer, R. Bieler, G. Giribet, Resolving the relationships of clams and cockles: dense transcriptome sampling drastically improves the bivalve tree of life. Proc. R. Soc. B Biol. Sci. 286, 20182684 (2019).

35. Y.-T. Lin, J.-W. Qiu, Reassessment of Pectinida (Mollusca: Bivalvia) phylogenetic relationships and description of a new *Parvamussium* species. Zool. J. Linn. Soc. 206, zlaf200 (2026).

36. Y. X. Li, et al., Phylogenomics of Bivalvia using ultraconserved elements reveal new topologies for Pteriomorphia and Imparidentia. Syst. Biol. 74, 16–33 (2025).

37. T. R. Waller, Phylogeny of families in the Pectinoidea (Mollusca: Bivalvia): importance of the fossil record. Zool. J. Linn. Soc. 148, 313–342 (2006).

38. L. R. L. Simone, V. S. do Amaral, Phenotypic features of *Dimya cf. japonica* (Bivalvia, Dimyidae) from Niue Island (South Pacific) with accounts on its phylogeny and taxonomic relationships. Malacologia 64, 121–136 (2021).

39. M. Hautmann, Taxonomy and phylogeny of cementing Triassic bivalves (families Prospondylidae, Plicatulidae, Dimyidae and Ostreidae). Palaeontology 44, 339–373 (2001).

40. F. Plazzi, A. Ceregato, M. Taviani, M. Passamonti, A molecular phylogeny of bivalve mollusks: ancient radiations and divergences as revealed by mitochondrial genes. PLoS One 6, e27147 (2011).

41. J. D. Sigwart, Y. Li, Z. Chen, K. Vončina, J. Sun, Still waters run deep in large-scale genome rearrangements of morphologically conservative Polyplacophora. eLife 13, RP10254 (2025).

42. L. H. Rieseberg, Chromosomal rearrangements and speciation. Trends Ecol. Evol. 16, 351–358 (2001).

43. J. M. De Vos, H. Augustijnen, L. Bätscher, K. Lucek, Speciation through chromosomal fusion and fission in Lepidoptera. Philos. Trans. R. Soc. B Biol. Sci. 375, 20190539 (2020).

44. F. Cicconardi, et al., Chromosome fusion affects genetic diversity and evolutionary turnover of functional loci but consistently depends on chromosome size. Mol. Biol. Evol. 38, 4449–4462 (2021).

45. S. Farhat, M. V. Modica, N. Puillandre, Whole genome duplication and gene evolution in the hyperdiverse venomous gastropods. Mol. Biol. Evol. 40, msad171 (2023).

46. Y.-S. Wang, et al., Chromosome-level genome assemblies of two littorinid marine snails indicate genetic basis of intertidal adaptation and ancient karyotype evolved from bilaterian ancestors. GigaScience 13, giae072 (2024).

47. Y. Wang, X. Guo, Chromosomal rearrangement in Pectinidae revealed by rRNA genes and their implications for phylogeny. Mar. Biol. 144, 1119–1126 (2004).

48. D. Grouzdev, et al., Chromosome-level genome assembly of the bay scallop *Argopecten irradians*. Sci. Data 11, 1057 (2024).

49. A. Navarro, N. H. Barton, Chromosomal speciation and molecular divergence–Accelerated evolution in rearranged chromosomes. Science 300, 321–324 (2003).

50. D. Dumas, J. Catalan, J. Britton-davidian, Reduced recombination patterns in Robertsonian hybrids between chromosomal races of the house mouse: chiasma analyses. Heredity 114, 56– 64 (2015).

51. V. Merico, et al., Chromosomal speciation in mice: A cytogenetic analysis of recombination. Chromosome Res. 21, 523–533 (2013).

52. T. Cremer, C. Cremer, Chromosome territories, nuclear architecture and gene regulation in mammalian cells. Nat. Rev. Genet. 2, 292–301 (2001).

53. T. Misteli, Beyond the sequence: Cellular organization of genome function. Cell 128, 787–800 (2007).

54. J. J. Yunis, O. Prakash, The origin of man: A chromosomal pictorial legacy. Science 215, 1525– 1530 (1982).

55. J. W. Ijdo, A. Baldini, D. C. Ward, S. T. Reeders, R. A. Wells, Origin of human chromosome 2: An ancestral telomere-telomere fusion. Proc. Natl. Acad. Sci. U. S. A. 88, 9051–9055 (1991).

56. A. Nietzel, et al., A new multicolor-FISH approach for the characterization of marker chromosomes: Centromere-specific multicolor-FISH (cenM-FISH). Hum. Genet. 108, 199–204 (2001).

57. R. Bracewell, K. Chatla, M. J. Nalley, D. Bachtrog, Dynamic turnover of centromeres drives karyotype evolution in *Drosophila*. eLife 8, e49002 (2019).

58. G. Chiatante, G. Giannuzzi, F. M. Calabrese, E. E. Eichler, M. Ventura, Centromere destiny in dicentric chromosomes: New insights from the evolution of human chromosome 2 ancestral centromeric region. Mol. Biol. Evol. 34, 1669–1681 (2017).

59. P. B. Talbert, S. Henikoff, What makes a centromere? Exp. Cell Res. 389, 111895 (2020).

60. R. Avarello, A. Pedicini, A. Caiulo, O. Zuffardi, M. Fraccaro, Evidence for an ancestral alphoid domain on the long arm of human chromosome 2. Hum. Genet. 89, 247–249 (1992).

61. D. J. Amor, K. H. A. Choo, Neocentromeres: role in human disease, evolution, and centromere study. Am. J. Hum. Genet. 71, 695–714 (2002).

62. O. J. Marshall, A. C. Chueh, L. H. Wong, K. H. Choo, Neocentromeres: New insights into centromere structure, disease development, and karyotype evolution. Am. J. Hum. Genet. 82, 261–282 (2008).

63. S. Osawa, T. Ohama, T. H. Jukes, K. Watanabe, Evolution of the mitochondrial genetic code. I. Origin of AGR serine and stop codons in metazoan mitochondria. J. Mol. Evol. 29, 202–207 (1989).

64. K. Watanabe, S. I. Yokobori, tRNA modification and genetic code variations in animal mitochondria. J. Nucleic Acids 2011, 623095 (2011).

65. A. T. Ho, L. D. Hurst, Stop codon usage as a window into genome evolution: Mutation, selection, biased gene conversion and the TAG paradox. Genome Biol. Evol. 14, evac115 (2022).

66. C. H. Hill, I. Brierley, Structural and functional insights into viral programmed ribosomal frameshifting. Annu. Rev. Virol. 10, 217–242 (2023).

67. T. Jacks, et al., Characterization of ribosomal frameshifting in HIV-1 gag-pol expression. Nature 331, 280–283 (1988).

68. J. L. Jacobs, A. T. Belew, R. Rakauskaite, J. D. Dinman, Identification of functional, endogenous programmed−1 ribosomal frameshift signals in the genome of *Saccharomyces cerevisiae*. Nucleic Acids Res. 35, 165–174 (2006).

69. H. B. Gamper, I. Masuda, M. Frenkel-Morgenstern, Y.-M. Hou, Maintenance of protein synthesis reading frame by EF-P and m1G37-tRNA. Nat. Commun. 6, 7226 (2015).

70. O. L. Gurvich, P. V. Baranov, R. F. Gesteland, J. F. Atkins, Expression levels influence ribosomal frameshifting at the tandem rare arginine codons AGG_AGG and AGA_AGA in *Escherichia coli*. J. Bacteriol. 187, 4023–4032 (2005).

71. C. L. Simms, L. L. Yan, J. K. Qiu, H. S. Zaher, Ribosome collisions result in +1 frameshifting in the absence of no-go decay. Cell Rep. 28, 1679–1689 (2019).

72. J. F. Atkins, G. Loughran, P. R. Bhatt, A. E. Firth, P. V. Baranov, Ribosomal frameshifting and transcriptional slippage: From genetic steganography and cryptography to adventitious use. Nucleic Acids Res. 44, 7007–7078 (2016).

73. J. D. Dinman, Mechanisms and implications of programmed translational frameshifting. Wiley Interdiscip. Rev. RNA 3, 661–673 (2012).

74. S. Matsufuji, et al., Autoregulatory frameshifting in decoding mammalian ornithine decarboxylase antizyme. Cell 80, 51–60 (1995).

75. Y. Xiao, R. Wang, X. Han, W. Wang, A. Liang, The deficiency of hypusinated eIF5A decreases the putrescine/spermidine ratio and inhibits +1 programmed ribosomal frameshifting during the translation of Ty1 retrotransposon in *Saccharomyces cerevisiae*. Int. J. Mol. Sci. 25, 1766 (2024).

76. P. J. Farabaugh, Programmed translational frameshifting. Microbiol. Rev. 60, 103–134 (1996).

77. S. Blanchet, et al., Deciphering the reading of the genetic code by near-cognate tRNA. Proc. Natl. Acad. Sci. U. S. A. 115, 3018–3023 (2018).

78. S. Sengupta, P. G. Higgs, A unified model of codon reassignment in alternative genetic codes. Genetics 170, 831–840 (2005).

79. S. Osawa, T. H. Jukes, Codon reassignment (codon capture) in evolution. J. Mol. Evol. 28, 271– 278 (1989).

80. D. W. Schultz, M. Yarus, Transfer RNA mutation and the malleability of the genetic code. J. Mol. Biol. 235, 1377–1380 (1994).

81. Y. Bessho, T. Ohama, S. Osawa, Planarian mitochondria II. The unique genetic code as deduced from cytochrome c oxidase subunit I gene sequences. J. Mol. Evol. 34, 331–335 (1992).

82. M. J. Telford, E. A. Herniou, R. B. Russell, D. T. J. Littlewood, Changes in mitochondrial genetic codes as phylogenetic characters: Two examples from the flatworms. Proc. Natl. Acad. Sci. U. S. A. 97, 11359–11364 (2000).

83. A. C. Christinaki, et al., Mitogenomics and mitochondrial gene phylogeny decipher the evolution of *Saccharomycotina* yeasts. Genome Biol. Evol. 14, evac073 (2022).

84. D. Žihala, J. Salamonová, M. Eliáš, Evolution of the genetic code in the mitochondria of Labyrinthulea (Stramenopiles). Mol. Phylogenet. Evol. 152, 106908 (2020).

85. T. R. Waller, Morphology, phylogeny, and systematic revision of genera in the Dimyidae (Mollusca, Bivalvia, Pteriomorphia). J. Paleontol. 86, 829–851 (2012).

86. C. M. Yonge, On the Dimyidae (Mollusca: Bivalvia) with special reference to *Dimya corrugata* Hedley and *Basiliomya goreaui* Bayer. J. Molluscan Stud. 44, 357–375 (1978).

87. R. Bieler, J. G. Carter, E. V. Coan, Classification of Bivalve families. Malacologia 52, 113–133 (2010).

88. R. Bieler, P. M. Mikkelsen, Bivalvia – a look at the Branches. Zool. J. Linn. Soc. 148, 223–235 (2006).

89. C. M. Yonge, Functional morphology with particular reference to hinge and ligament in *Spondylus* and *Plicatula* and a discussion on relations within the superfamily Pectinacea (Mollusca: Bivalvia). Philos. Trans. R. Soc. Lond. B Biol. Sci. 267, 173–208 (1973).

90. C. M. Yonge, Form and evolution in the Anomiacea (Mollusca: Bivalvia)—*Pododesmus*, *Anomia*, *Patro*, *Enigmonia* (Anomiidae); *Placunanomia*, Placuna (Placunidae fam. nov.). Philos. Trans. R. Soc. Lond. B Biol. Sci. 276, 453–523 (1977).

91. C. M. Yonge, The monomyarian condition in the Lamellibranchia. Trans. R. Soc. Edinb. 62, 443–478 (1953).

92. P. M. Mikkelsen, R. Bieler, Seashells of southern Florida: Living marine mollusks of the Florida Keys and adjacent regions: Bivalves (Princeton University Press, 2021).

93. N. D. Newell, “Classification of Bivalvia” in Treatise on Invertebrate Paleontology, Part N, Mollusca 6, Bivalvia, R. C. Moore, Ed. (Geological Society of America and University of Kansas, 1969), pp. N205–N224.

94. M. Hautmann, Early Mesozoic evolution of alivincular bivalve ligaments and its implications for the timing of the “Mesozoic marine revolution.” Lethaia 37, 165–172 (2004).

95. R. Rose, T. Dix, Larval and juvenile development of the doughboy scallop, *Chlamys (Chlamys) asperrimus* (Lamarck) (Mollusca: Pectinidae). Mar. Freshw. Res. 35, 315–323 (1984).

96. P. D. Chantler, “Chapter 4 Scallop adductor muscles: Structure and function” in *Scallops: Biology, Ecology and Aquaculture*, (Elsevier, 2006), pp. 229–316.

97. D. Atkins, On the Ciliary Mechanisms and Interrelationships of Lamellibranchs.: PART V: Note on the Gills of *Amussium pleuronectes*. J. Cell Sci. s2-80, 321–329 (1938).

98. J. A. Audino, J. E. A. Marian, A. Wanninger, S. G. Lopes, Mantle margin morphogenesis in *Nodipecten nodosus* (Mollusca: Bivalvia): new insights into the development and the roles of bivalve pallial folds. BMC Dev. Biol. 15, 22 (2015).

99. P. J. Hayward, J. S. Ryland, The marine fauna of the British Isles and North-West Europe: 2. Molluscs to chordates (Oxford University Press, 1990).

100. Y.-T. Lin, Y.-X. Li, H.-X. Loke, X. Han, J.-W. Qiu, One becomes three: An integrative morphological and molecular analysis of the windowpane oyster *Placuna* (Bivalvia: Pectinida) reveals new species. Ecol. Evol. 14, e70260 (2024).

101. C. M. Yonge, The status of the Plicatulidae and the Dimyidae in relation to the superfamily Pectinacea (Mollusca: Bivalvia). J. Zool. 176, 545–553 (1975).

102. C. N. Stewart Jr., L. E. Via, A rapid CTAB DNA isolation technique useful for RAPD fingerprinting and other PCR applications. BioTechniques 14, 748–750 (1993).

103. A. M. Bolger, M. Lohse, B. Usadel, Trimmomatic: A flexible trimmer for Illumina sequence data. Bioinformatics 30, 2114–2120 (2014).

104. A. Bankevich, et al., SPAdes: A new genome assembly algorithm and its applications to single-cell sequencing. J. Comput. Biol. 19, 455–477 (2012).

105. N. Dierckxsens, P. Mardulyn, G. Smits, NOVOPlasty: *De novo* assembly of organelle genomes from whole genome data. Nucleic Acids Res. 45, e18 (2017).

106. C. Camacho, et al., BLAST+: Architecture and applications. BMC Bioinformatics 10, 421 (2009).

107. S. Greiner, P. Lehwark, R. Bock, OrganellarGenomeDRAW (OGDRAW) version 1.3.1: expanded toolkit for the graphical visualization of organellar genomes. Nucleic Acids Res. 47, W59–W64 (2019).

108. Y. Shulgina, S. R. Eddy, A computational screen for alternative genetic codes in over 250,000 genomes. eLife 10, e71402 (2021).

109. P. J. A. Cock, et al., Biopython: freely available Python tools for computational molecular biology and bioinformatics. Bioinformatics 25, 1422–1423 (2009).

110. S. Hussain, S. T. Rasool, A. H. Asif, A detailed analysis of synonymous codon usage in human bocavirus. Arch. Virol. 164, 335–347 (2019).

111. K. Katoh, D. M. Standley, MAFFT multiple sequence alignment software version 7: Improvements in performance and usability. Mol. Biol. Evol. 30, 772–780 (2013).

112. D. Zhang, et al., PhyloSuite: An integrated and scalable desktop platform for streamlined molecular sequence data management and evolutionary phylogenetics studies. Mol. Ecol. Resour. 20, 348–355 (2020).

113. G. Talavera, J. Castresana, Improvement of phylogenies after removing divergent and ambiguously aligned blocks from protein sequence alignments. Syst. Biol. 56, 564–577 (2007).

114. S. Kalyaanamoorthy, B. Q. Minh, T. K. F. Wong, A. von Haeseler, L. S. Jermiin, ModelFinder: Fast model selection for accurate phylogenetic estimates. Nat. Methods 14, 587– 589 (2017).

115. L.-T. Nguyen, H. A. Schmidt, A. von Haeseler, B. Q. Minh, IQ-TREE: A fast and effective stochastic algorithm for estimating maximum-likelihood phylogenies. Mol. Biol. Evol. 32, 268– 274 (2015).

116. B. Q. Minh, M. A. Nguyen, A. von Haeseler, Ultrafast approximation for phylogenetic bootstrap. Mol. Biol. Evol. 30, 1188–1195 (2013).

117. S. Guindon, et al., New algorithms and methods to estimate maximum-likelihood phylogenies: Assessing the performance of PhyML 3.0. Syst. Biol. 59, 307–321 (2010).

118. J. Cox, M. Mann, MaxQuant enables high peptide identification rates, individualized p.p.b.- range mass accuracies and proteome-wide protein quantification. Nat. Biotechnol. 26, 1367– 1372 (2008).

119. K. P. Schliep, phangorn: Phylogenetic analysis in R. Bioinformatics 27, 592–593 (2011).

120. J. Felsenstein, Inferring phylogenies (Sinauer Associates, 2004).

121. K. C. Nixon, The Parsimony Ratchet, a new method for rapid parsimony analysis. Cladistics 15, 407–414 (1999).

122. J. Felsenstein, Confidence limits on phylogenies: An approach using the bootstrap. Evolution 39, 783–791 (1985).

123. G. Yu, D. K. Smith, H. Zhu, Y. Guan, T. T. Lam, ggtree: An R package for visualization and annotation of phylogenetic trees with their covariates and other associated data. Methods Ecol. Evol. 8, 28–36 (2017).

124. G. Marçais, C. Kingsford, A fast, lock-free approach for efficient parallel counting of occurrences of k-mers. Bioinformatics 27, 764–770 (2011).

125. T. R. Ranallo-Benavidez, K. S. Jaron, M. C. Schatz, GenomeScope 2.0 and Smudgeplot for reference-free profiling of polyploid genomes. Nat. Commun. 11, 1432 (2020).

126. H. Cheng, G. T. Concepcion, X. Feng, H. Zhang, H. Li, Haplotype-resolved *de novo* assembly using phased assembly graphs with hifiasm. Nat. Methods 18, 170–175 (2021).

127. M. J. Roach, S. A. Schmidt, A. R. Borneman, Purge Haplotigs: Allelic contig reassignment for third-gen diploid genome assemblies. BMC Bioinformatics 19, 460 (2018).

128. A. Mikheenko, A. Prjibelski, V. Saveliev, D. Antipov, A. Gurevich, Versatile genome assembly evaluation with QUAST-LG. Bioinformatics 34, i142–i150 (2018).

129. N. Huang, H. Li, Compleasm: A faster and more accurate reimplementation of BUSCO. Bioinformatics 39, btad595 (2023).

130. N. Servant, et al., HiC-Pro: An optimized and flexible pipeline for Hi-C data processing. Genome Biol. 16, 259 (2015).

131. N. C. Durand, et al., Juicer provides a one-click system for analyzing loop-resolution Hi-C experiments. Cell Syst. 3, 95–98 (2016).

132. O. Dudchenko, et al., *De novo* assembly of the Aedes aegypti genome using Hi-C yields chromosome-length scaffolds. Science 356, 92–95 (2017).

133. N. C. Durand, et al., Juicebox provides a visualization system for Hi-C contact maps with unlimited zoom. Cell Syst. 3, 99–101 (2016).

134. W. Bao, K. K. Kojima, O. Kohany, Repbase Update, a database of repetitive elements in eukaryotic genomes. Mob. DNA 6, 11 (2015).

135. J. M. Flynn, et al., RepeatModeler2 for automated genomic discovery of transposable element families. Proc. Natl. Acad. Sci. U. S. A. 117, 9451–9457 (2020).

136. B. L. Cantarel, et al., MAKER: An easy-to-use annotation pipeline designed for emerging model organism genomes. Genome Res. 18, 188–196 (2008).

137. M. G. Grabherr, et al., Full-length transcriptome assembly from RNA-Seq data without a reference genome. Nat. Biotechnol. 29, 644–652 (2011).

138. D. Kim, B. Langmead, S. L. Salzberg, HISAT: A fast spliced aligner with low memory requirements. Nat. Methods 12, 357–360 (2015).

139. B. J. Haas, et al., Improving the Arabidopsis genome annotation using maximal transcript alignment assemblies. Nucleic Acids Res. 31, 5654–5666 (2003).

140. M. Stanke, B. Morgenstern, AUGUSTUS: A web server for gene prediction in eukaryotes that allows user-defined constraints. Nucleic Acids Res. 33, W465–W467 (2005).

141. B. J. Haas, et al., Automated eukaryotic gene structure annotation using EVidenceModeler and the program to assemble spliced alignments. Genome Biol. 9, R7 (2008).

142. M. Caballero, J. Wegrzyn, gFACs: Gene filtering, analysis, and conversion to unify genome annotations across alignment and gene prediction frameworks. Genomics Proteomics Bioinformatics 17, 305–310 (2019).

143. J. Huerta-Cepas, et al., eggNOG 5.0: A hierarchical, functionally and phylogenetically annotated orthology resource based on 5090 organisms and 2502 viruses. Nucleic Acids Res. 47, D309–D314 (2019).

144. B. Buchfink, C. Xie, D. H. Huson, Fast and sensitive protein alignment using DIAMOND. Nat. Methods 12, 59–60 (2015).

145. D. M. Emms, S. Kelly, OrthoFinder: Phylogenetic orthology inference for comparative genomics. Genome Biol. 20, 238 (2019).

146. Z. Yang, PAML 4: Phylogenetic analysis by maximum likelihood. Mol. Biol. Evol. 24, 1586–1591 (2007).

147. H. Yu, et al., Pan-evolutionary and regulatory genome architecture delineated by an integrated macro- and microsynteny approach. Nat. Protoc. 19, 1623–1678 (2024).

148. Y. Wang, et al., MCScanX: A toolkit for detection and evolutionary analysis of gene synteny and collinearity. Nucleic Acids Res. 40, e49 (2012).

149. W. He, Y. Zhang, Z. Li, J. Wang, NGenomeSyn: An easy-to-use and flexible tool for publication-ready visualization of syntenic relationships across multiple genomes. Bioinformatics 39, btad121 (2023).

150. C. Elphinstone, R. Elphinstone, M. Todesco, L. Rieseberg, RepeatOBserver: tandem repeat visualization and centromere detection. bioRxiv 2023.12.30.573697 (2023). 10.1101/2023.12.30.573697.

151. F. Cabanettes, C. Klopp, D-GENIES: dot plot large genomes in an interactive, efficient and simple way. PeerJ 6, e4958 (2018).

152. H. Pagès, P. Aboyoun, R. Gentleman, S. DebRoy, Biostrings: Efficient manipulation of biological strings. (2019).

