## Supplementary Information for "Life finds a way: Integrative phylogenomics resolves an overlooked bivalve order with chromosome fusion and mitochondrial translational-code evolution"

#### Mitochondrial Conservation and Divergence in the APP Lineage

Mitochondrial gene order is highly variable among mollusks, with bivalves exhibiting the most pronounced variability across all metazoan groups (1, 2). Within the infraclass Pteriomorphia, mitochondrial gene orders exhibit significant diversity, with 44 distinct patterns identified (excluding tRNA genes), underscoring the extraordinary lability of mitochondrial genome architecture within this subclass (3). The APP lineage (comprising Anomiidae, Placunidae, and Plicatulidae) exhibits a pattern that aligns with this broader molluscan variability: mitochondrial gene orders appear to be generally conserved within genera while displaying remarkable structural diversity among genera, forming a distinct hierarchical differentiation (Fig. S6-7). We found that the three species of *Placuna* share identical gene orders, indicating that the mitochondrial genome structure of the windowpane oysters has remained highly stable following speciation. The two-color morphs of *Enigmonia aenigmatica* show similarly minimal divergence (distance = 12), and the two *Plicatula* species exhibit a distance of only 25, suggesting that these genera have maintained conserved gene arrangements even across species boundaries. Within *Anomia*, the geographically separated populations of *Anomia trigonopsis* from New Zealand and Mauritania share identical gene orders, while *Anomia chinensis* shows a distance of 56 from *A. trigonopsis*, and *Anomia ephippium* exhibits distances of 82–84 from other *Anomia* species. Although these values indicate that *Anomia* has undergone more structural differentiation than other APP genera, the rearrangements remain limited, and the overall gene order synteny is still clearly recognizable. Additionally, the three geographically distant populations of *Heteranomia squamula* (Russia, UK, USA) exhibit distances of 0–25, further supporting the high intra-generic conservation of gene order. This pattern aligns with the broader observation in bivalves that gene sequences are highly variable among species, yet gene arrangements tend to be more consistent among closely related taxa

(1).

In striking contrast, inter-generic divergence in gene order is profound, with distances far exceeding intra-generic distances and each genus exhibiting distinctive structural features. The magnitude of this inter-generic divergence is comparable to the substantial rearrangements observed at the intra-familial level within Pectinidae, where scallops (*Argopecten irradians*, *Chlamys farreri*, *Mizuhopecten yessoensis*, and *Placopecten magellanicus*) exhibit substantial rearrangements despite belonging to the same family, a level of variation seldom observed in other metazoan or even molluscan classes (4, 5). Indeed, pectinids present extraordinary variance in mitochondrial genome size, structure, and content. At the scale of the Pectinidae, 15 sequence blocks are involved in mitogenome rearrangements, which behave as separate units, and different genome organization patterns can be observed even at the level of Tribes (6). Such extensive rearrangements occurring at the familial level in Pectinidae provide an informative context for interpreting the inter-generic divergences observed within the APP lineage. Additionally, we found that Spondylidae, a family within Pectinida, exhibits highly conserved mitochondrial gene orders. Spondylidae is a small group comprising a single genus (*Spondylus*) with approximately 70 living species, all of which are sessile, cementing their right valve to hard substrates. This presents an intriguing contrast to the variability seen in Pectinidae and within the APP lineage.

The observed pattern of generally conserved gene orders within genera alongside substantial inter-generic divergence reflects the differentiation also observed in morphology, ecology, and genetic code variation (TAA=Tyr, code 5, and +1 frameshifting) across these lineages. This suggests that the mitochondrial genome architecture of the APP lineage may have experienced a relatively punctuated mode of evolution at the generic level, wherein gene order rearrangements occurred primarily during ancestor divergence and subsequently remained stable (3, 4). In contrast, a more gradual accumulation of gene order changes has been observed in some other molluscan lineages; for example, in Vermetidae gastropods, extensive gene order changes have occurred at a scale unexpected for such an evolutionarily young group (7), suggesting that in some lineages, rearrangements can accumulate progressively over shorter evolutionary timescales. This pattern is consistent with observations in other bivalve groups, where gene rearrangement among families is particularly obvious, while gene order within families is relatively conservative, such as those of Adapedonta and Veneridae (1, 8, 9). Collectively, the APP lineage exemplifies a broader evolutionary tendency observed across Bivalvia, wherein mitochondrial gene order rearrangements occur predominantly at higher taxonomic levels yet remain relatively stable within genera.

The pronounced inter-generic divergence in mitochondrial gene order documented within the APP lineage raises the question of what mechanisms might underlie such extensive structural variation. Evidence from diverse molluscan lineages suggests that the tRNA gene family may play a particularly active role in driving mitochondrial genome rearrangements, with gene duplication, loss, isomerism, recruitment, and positional changes occurring frequently across multiple invertebrate groups (2, 10).

Duplications of tRNA genes are thought to be a major contributor to mitochondrial genome rearrangement, consistent with the tandem duplication-random loss (TDRL) model, and it is well established that many mollusks contain extra tRNAs beyond the minimal set of 22 essential for accommodating the super-wobble of mitochondrial translation (8). Our observation of lineage-specific tRNA duplications, such as the *trnT* and *trnA* duplications in *Heteranomia aculeata*, and the duplicated *trnM* and *trnW* genes in *Plicatula* and *Pododesmus*, provides evidence consistent with the TDRL model operating in the APP lineage. This model, first documented in *Crassostrea* oysters, proposes that genomic regions undergo duplication followed by differential loss of redundant copies, thereby generating hotspots of rearrangement (8). The presence of multiple, independently derived tRNA duplications across phylogenetically disparate APP genera suggests that TDRL may be a recurrent process in this clade, contributing to the structural diversity observed among genera.

**Doubly Uniparental Inheritance in *Heteranomia aculeata***

In addition to the +1 translational frameshifting mechanism described in the main text, our analyses revealed that the species *Heteranomia aculeata* exhibits doubly uniparental inheritance (DUI) of the mitogenome, a distinctive reproductive strategy in which female- and male-transmitted mitochondrial lineages are maintained separately (11). However, examination of specimens collected from three geographically distinct localities (Devon, UK; Massachusetts, USA; and the Barents Sea, Russia) revealed no evidence of DUI in the sole congener, *H. squamula*. DUI represents a notable exception to the typical maternal inheritance of mitochondrial DNA in metazoans (12). In bivalves, DUI manifests as the concomitant presence of two distinct mitochondrial lineages: the F-type, transmitted by females, and the M-type, transmitted exclusively through males (13, 14). This system has been documented in approximately 100 bivalve species across six orders (12, 15). In *H. aculeata*, based on our Illumina sequencing data, the F-type and M-type mitochondrial genomes (Fig. S5) are similar in size (22,400 bp and 22,996 bp, respectively) and GC content (34.05% and 33.89%, respectively). Read-depth analysis revealed a marked copy number disparity, with the F-type mitogenome maintained at approximately 12-fold higher abundance than its M-type counterpart. Sequence divergence between the two sex-associated mitogenomes is substantial, with only 85.10% nucleotide identity, a value comparable to the interspecific divergence observed between *H. aculeata* and *H. squamula* (80%). This marked divergence underscores the pronounced heteroplasmy and evolutionary independence of the two lineages. Such differentiation between sex-associated mitochondrial genomes is not without precedent: gene duplication and loss contribute to variation in mitochondrial genome content both among bivalve taxa and between male and female mitotypes within a single species (16). Similar patterns of extensive divergence have been observed in other DUI systems, such as in *Mytilus* mussels, where sequence divergence between sex-linked mitotypes can exceed 20% (17, 18). Therefore, DUI represents an additional dimension of mitogenomic complexity in Bivalvia (11, 13) .

Gene content and organization are largely conserved between these two mitogenomes, yet several distinctive features warrant attention. The F-type mitogenome lacks *trnR* and contains a severely truncated *cox3* gene retaining only a 5’-end fragment of approximately 800 bp; this remnant is tandemly duplicated and appears to represent non-functional pseudogenes (Fig. S5). The M-type mitogenome, by contrast, retains an intact *trnR* copy and a full-length *cox3*, but has lost *atp6* entirely (Fig. S5). A plausible interpretation is that these gene losses reflect sex-biased degenerative processes: the pseudogenization of *cox3* and loss of *trnR* in the F-type, and the loss of *atp6* in the M-type, may be tolerated in somatic tissues and the male germline, respectively, because each sex-specific mitochondrial lineage experiences different selective pressures independently. This phenomenon is reminiscent of, yet distinct from, the situation in *Mytilus* mussels, where *atp8* is pseudogenized in the M-type but retained in the F-type, suggesting a sex-specific pattern of gene degeneration rather than functional complementation (15). The gene-loss pattern observed in *H. aculeata* may reflect a similar sex-biased process, but involves different genes (*cox3*, *trnR*, *atp6*), indicating lineage-specific features of DUI-associated degradation. The maintenance of the M-type lineage ensures that an intact *cox3* copy is preserved within the species' gene pool, specifically for male germline function. However, because F- and M-type mitochondria are largely segregated in different tissues, this does not constitute a functional complementation in somatic cells. Instead, this pattern exemplifies how DUI permits the tolerance of gene loss in one sex-specific lineage without causing a species-wide fitness defect. To our knowledge, this represents the first such report within Pectinida. Ghiselli et al. (2013) further demonstrated that mitochondrial transcription in DUI species is lineage-specific, and that F- and M-type mtDNAs harbor similar levels of polymorphism but of different types, reflecting differential population sizes and selection efficiencies (13).

The obligate DUI in *H. aculeata* may represent an evolutionary dependency: while functional specialization might confer certain selective advantages (e.g., reduced replication burden), the loss of key genes in the F-type mitogenome renders the species irrevocably dependent on the retention of the M-type lineage. Should DUI fail, whether through loss of the M-type lineage during transmission or male sterility preventing M-type inheritance, individuals would likely be inviable due to respiratory chain dysfunction. The lack of DUI and the retention of a full set of mitochondrial genes in *H. squamula* further support the interpretation that the emergence of DUI in *H. aculeata* represents a unique and potentially irreversible event in its evolutionary history. The costs associated with this dependency are multifaceted. First, each generation must produce functional M-type mitochondria for sperm transmission, imposing strong selective pressure on male germline mitochondrial quality. Second, the nuclear genome must maintain compatibility with two divergent mitochondrial genomes; with sequence divergence approaching 15% between the two lineages, this dual compatibility imposes a substantial coevolutionary burden. Third, once DUI becomes obligate, the species loses the flexibility to respond to mitochondrial gene loss through alternative mechanisms. Nevertheless, this system may also confer benefits: functional specialization between the two mitochondrial genomes could permit independent optimization in female and male germlines, representing a potential selective force driving the repeated evolution of DUI systems in bivalves (12, 15).

#### Role of Directional Mutation Pressure in Mitogenomes

A fundamental question arising from our observations is why the TAA=Tyr reassignment documented in *Anomia*, *Enigmonia*, *Placuna*, and *Plicatula* did not also evolve in *Pododesmus* and *Heteranomia*. The differential outcomes among these closely related lineages provide a natural experiment for understanding the conditions that favor codon reassignment versus alternative translational recoding strategies. Unlike other APP lineages, *Pododesmus* completely lacks in-frame stop codons within its mitochondrial PCGs and retains the canonical translation table 5 without modification. *Heteranomia*, by contrast, contains numerous in-frame TAG codons but has evolved a +1 translational frameshift mechanism to circumvent them, rather than reassigning the genetic code.

The codon capture hypothesis posits that codon reassignment is facilitated by directional mutation pressure that reduces the frequency of a particular codon to the point of disappearance, thereby removing the deleterious consequences of its reassignment (19). Our comparative analysis of base composition across mitochondrial PCGs reveals a pattern consistent with this model. In TAA=Tyr lineages, adenine content at the third codon position is significantly elevated (mean = 31.52%, SD = 3.61%), whereas *Pododesmus* exhibits substantially lower A content at this position (mean = 22.71%, SD = 0.46%), and *Heteranomia* displays an intermediate value (mean = 24.52%, SD = 2.12%). This difference is critical because in-frame TAA codons arise when adenine occupies the third position, driven by directional mutation pressure, which increases the probability of TAA occurrence. The substantial A bias in TAA=Tyr lineages likely generated strong selective pressure by producing numerous in-frame TAA codons that would prematurely terminate translation. In *Pododesmus*, the absence of such pronounced A bias resulted in no in-frame TAA codons, thereby eliminating the selective pressure for TAA reassignment. This observation is broadly consistent with the codon capture hypothesis: the mutational pressure that creates the problem is likely a prerequisite for driving the evolutionary innovation that solves it.

*Heteranomia* presents an intermediate case. Despite a moderate A-bias at the third position (24.52%) that falls between *Pododesmus* (22.71%) and the TAA=Tyr lineages (31.52%), *Heteranomia* did not evolve TAA reassignment. This can be explained by its specific nucleotide composition at the second and third codon positions. The formation of in-frame TAA codons requires adenine at both the second and third positions; the formation of in-frame TAG codons requires adenine at the second position and guanine at the third. In *Heteranomia*, the second-position A content (mean = 21.23%) is somewhat higher than in TAA=Tyr lineages (mean = 18.77%), while the third-position G content (mean = 21.13%) is also elevated relative to TAA=Tyr lineages (mean = 17.71%). This nucleotide composition—characterized by high A at the second position combined with elevated G at the third—may have favored the accumulation of TAG codons over TAA codons. In contrast, TAA=Tyr lineages exhibit a more extreme third-position A bias (31.52%) coupled with lower third-position G content (17.71%).

It should be noted, however, that the codon capture hypothesis is not the only mechanism proposed to explain codon reassignment. Knight et al. (2001) demonstrated that low codon frequencies can be related to codon reassignment but appear to be neither necessary nor sufficient for reassignment to occur (20). The ambiguous intermediate hypothesis (21) proposes that a tRNA mutation allows a codon to be recognized by two different decoding molecules; if beneficial, the new assignment can become fixed. Sengupta and Higgs (2005) developed a unified framework incorporating both codon disappearance and ambiguous intermediate mechanisms, demonstrating that directional mutation pressure is one of several parameters influencing the likelihood of reassignment (22). The elevated A bias observed in TAA=Tyr lineages is therefore consistent with a codon capture-like process, but does not definitively prove it. The relative contributions of these mechanisms in driving the independent origins of TAA=Tyr reassignment in the APP lineage remain to be fully resolved. Nevertheless, the observation that *Pododesmus* lacks in-frame TAA codons entirely, coupled with the absence of the U36C tRNA substitution, provides strong correlative support for the hypothesis that both the generation of the problem (A bias producing TAA codons) and the molecular innovation necessary to solve it (tRNA anticodon loop modification) are required for codon reassignment to occur.

**Detailed Evaluation of the Reassignment Hypotheses**

Two major hypotheses have been proposed to explain codon reassignment (Knight et al., 2001). The codon capture hypothesis suggests that a codon must first disappear from the genome before it can be reassigned to a different meaning (19). The ambiguous intermediate hypothesis proposes that a tRNA mutation allows a codon to be recognized by two different decoding molecules; if beneficial, the new assignment can become fixed (21). These two hypotheses are not mutually exclusive, and codon reassignment may involve elements of both (23). In the APP lineage, the independent emergence of TAA=Tyr reassignment provides a compelling case to test these models. The elevated A content at the third codon position is consistent with the codon capture hypothesis: directional mutation pressure led to a biased nucleotide composition, potentially causing an overrepresentation of in frame TAA codons. Its elevated frequency in the lineage would have imposed strong selective pressure due to premature translation termination, potentially driving the reassignment of TAA to tyrosine even in the absence of complete codon loss.

**Phylogenetic Affinities of Dimyidae**

Phylogenomic analyses of the APP lineages for which genome data are available robustly place Anomiidae, Placunidae, and Plicatulidae as sister to the Limida + Pectinidae/Propeamussiidae clade, with divergence estimated at approximately 428 MYA (Fig. 1-2). Specimens of Dimyidae were unavailable for genomic sequencing due to specimen rarity. Nevertheless, multiple lines of morphological and molecular phylogenetic evidence support its placement within the APP lineage.

Multi-gene analyses have consistently recovered Dimyidae, Plicatulidae, and Anomioidea as a monophyletic clade representing the earliest diverging lineage within Pectinida, forming a sister group to all other pectinids and embedded limids (24). This topology is corroborated by independent phylogenomic analyses using ultraconserved elements, which recovered congruent relationships for Pteriomorphia (25). Simone and Do Amaral (2021) further demonstrated that dimyids share four synapomorphies with pectinoideans, placing them between ostreoideans and pectinoideans but closer to the latter, a placement consistent with the topology recovered by molecular phylogenies (26). Hautmann (2001) demonstrated that shell microstructure and conchological characters do not support a close relationship between Dimyidae and Ostreidae, refuting the traditional placement of dimyids within or near Ostreidae (27). Consequently, the available molecular evidence firmly supports the inclusion of Dimyidae within the APP lineage and, by extension, within the proposed new order Anomiida.

The morphological comparison below demonstrates that the APPD lineage (Anomiidae, Placunidae, Plicatulidae, and Dimyidae) constitutes a coherent group distinct from other Pectinida families, based on a unique combination of characters.

#### Ligament structure

The ligament provides one of the most informative characters for distinguishing major groups. In Pectinidae and Propeamussiidae, the massive rounded inner ligament layer is bounded by long stretches of anterior and posterior outer ligament layers, with the ligament maintaining a longitudinal disposition (28). In Spondylidae, the outer layers migrate centrally to unite on either side of the inner layer, with the primary ligament becoming transversely disposed (29). In Cyclochlamydidae, a small family of minute pectinoids separated from Propeamussiidae, the ligament is reduced, and the hinge is simplified, consistent with their diminutive size (30). By contrast, the APPD lineage exhibits a fundamentally different ligamentary configuration. In Plicatulidae and Dimyidae, the transverse primary ligament is overarched by the mantle margins, producing a continuous longitudinal external secondary ligament above the internal primary ligament (28). In Anomiidae, the ligament is reduced to a small, drop-shaped structure with a largely internal resilium (31). In Placunidae, the ligament takes the form of a characteristic inverted V-shaped resilium, complemented by a secondary periostracal ligament that maintains valve alignment (31). This diversity of ligament types within the APPD lineage—from the reduced resilium of Anomiidae to the inverted V-shape of Placunidae and the transverse primary ligament with overarching secondary ligament in Plicatulidae and Dimyidae—is unique among Pectinida and collectively distinguishes the APPD lineage from the more uniform ligament configurations of Pectinidae, Propeamussiidae, Cyclochlamydidae, and Spondylidae.

#### Hinge structure and dentition

Hinge dentition also varies markedly across Pectinida. Pectinidae, Propeamussiidae, and Cyclochlamydidae are characterized by an edentulous or simplified hinge, with valve alignment maintained primarily by the ligament (28, 30). Spondylidae, in contrast, possess well-developed hinge teeth and corresponding sockets that form ball-and-socket joints, a derived condition associated with cementation (29). The APPD lineage, by contrast, exhibits a characteristic pattern of hinge reduction and modification. Anomiidae possesses a short or degraded hinge with absent hinge teeth, while Placunidae also has a reduced hinge, although distinct hinge teeth are present in some species (31). Plicatulidae and Dimyidae share the most derived condition, with strong crura-like hinge teeth articulating with corresponding sockets (28, 29, 32, 33). In Plicatulidae, the hinge features a trigonal resilifer flanked by prominent secondary isodont teeth, with V-shaped hinge teeth that tightly fit between the valves (32). Li et al. (2026) further examined the iterative evolution of hinge teeth in cementing bivalve families including Plicatulidae, providing additional paleontological context for understanding hinge diversification within this clade (34). Despite this variation, all APPD families share a general trend toward hinge reduction or modification, contrasting with the edentulous hinge of Pectinidae and Propeamussidae, the simplified hinge of Cyclochlamydidae, and the ball-and-socket hinge of Spondylidae.

**Adductor musculature**

All families of Pectinida, with the sole exception of Dimyidae, are monomyarian, possessing a single adductor muscle (28, 29, 31, 32). Dimyidae uniquely retains an anterior adductor muscle, a primitive feature lost in all other monomyarian bivalves (35). This retention is particularly significant for understanding the evolutionary trajectory of the APPD lineage. Yonge (1975) proposed that the monomyarian condition in Plicatulidae likely arose from a dimyarian ancestor similar to Dimyidae, with the transition occurring following cementation rather than through byssal attachment (28). Under this scenario, the loss of the anterior adductor in Plicatulidae and other APP families represents a secondary event, whereas the uniform monomyarian condition in Pectinidae, Propeamussiidae, and Spondylidae reflects a distinct evolutionary pathway associated with byssal attachment and subsequent loss of the anterior adductor (28). Thus, within the APPD lineage, adductor musculature ranges from the primitive dimyarian condition in Dimyidae to the derived monomyarian condition in Anomiidae, Placunidae, and Plicatulidae, reflecting a reduction in adductor musculature that is unique among Pectinida, with Dimyidae retaining the ancestral dimyarian condition while all other families in the order are monomyarian. This difference—Dimyidae retaining the anterior adductor while all other Pectinida families have lost it—provides strong morphological evidence for the early divergence of Dimyidae and supports its placement within the APPD lineage rather than with other pectinidan families.

**Lifestyle and attachment modes**

Pectinidae and Propeamussiidae are predominantly free-living or byssally attached, with many species capable of swimming (28, 29). Cyclochlamydidae are minute, free-living bivalves often found in deeper waters, with a byssal notch present in some taxa (30). Spondylidae are cemented by the right valve to hard substrates (29). The APPD lineage, however, exhibits a more diverse array of attachment strategies than typically observed in other pectinidan families, ranging from byssal attachment and free-living immobility to cementation. Anomiidae attach via a calcified byssus passing through a hole in the right valve; remarkably, *Enigmonia* (Anomiidae) has even reacquired mobility, crawling on mangrove leaves and stems (31). Placunidae lose their byssal apparatus in adulthood and adopt a free-living yet immobile existence on soft substrates (31). Plicatulidae and Dimyidae, in contrast, are cemented by the right valve to hard substrates, a habit they share with Spondylidae (28, 29, 32). This diversity of attachment modes within the APPD lineage—ranging from byssal attachment to free-living immobility to cementation—contrasts sharply with the more uniform lifestyles of other pectinidan families. The shared occurrence of cementation in Plicatulidae and Dimyidae, coupled with their common ligamentary and hinge modifications, suggests that these two families represent a cemented subclade within the APPD lineage, while Anomiidae and Placunidae have evolved alternative attachment strategies.

**Shell outline and ornamentation**

Shell morphology further reinforces the distinction between the APPD lineage and other Pectinida. Pectinidae, Propeamussiidae, and Cyclochlamydidae typically possess fan-shaped shells with prominent auricles (28–30). Spondylidae species also retain auricles, though they may be reduced (29). In the APPD lineage, by contrast, auricles are consistently absent. Anomiidae, Placunidae, Plicatulidae, and Dimyidae all lack auricles, and their shells are generally oval, suborbicular, or irregular in shape, reflecting their sessile or cemented lifestyles (28, 31, 33, 35). This shared absence of auricles, combined with generally reduced or irregular shell outlines, clearly distinguishes the APPD lineage from the fan-shaped, auriculate shells that characterize Pectinidae, Propeamussiidae, Cyclochlamydidae, and Spondylidae.

The morphological characters summarized above—including distinctive ligament configurations, hinge reduction or modification, diverse attachment modes, absence of auricles, and shell microstructural evidence—collectively define the APPD lineage as a morphological unit distinct from the families of Pectinida. While each character may vary within the APPD lineage, the combination of these features is not observed in any other pectinidan family. It must be acknowledged, however, that morphological data alone cannot confirm whether Dimyidae shares the distinctive mitochondrial genomic features documented in other APP lineages. Future collection efforts targeting Dimyidae for genomic sequencing will be critical to resolving these questions. Nevertheless, the congruence between morphological, paleontological, and molecular phylogenetic data firmly supports the placement of Dimyidae within the APP lineage and its inclusion in the new order Anomiida, while distinguishing this clade from other families of Pectinida.

**Genome size estimation and assembly**

Based on the Illumina sequences generated from the mantle tissue, the genome of *Anomia chinensis* was estimated to be 491.77 Mb in length, with a heterogeneity of 1.54% and a duplication of 2.25% (Fig. S1a). The assembly size was slightly larger than predicted, with 530.9 Mb for the chromosomal-level assembly, with 7 chromosomes (Fig. S1d; Table 1). Mapping the PacBio HiFi reads to the final genome revealed a sequencing coverage of 153.1× and a mapping rate of 99.9%. Besides, 86.8% of predicted genes (19,874 of 22,896) were functionally annotated against public databases. The quality assessment showed comparable BUSCO completeness of the genome (96.7%) and proteins (97.5%) from the predicted gene models.

For *Placuna vitream*, the genome was estimated to be 464.68 Mb, with a heterogeneity of 0.85% and a duplication of 0.64% (Fig. S1b). The chromosomal-level assembly reached 548.9 Mb with 8 chromosomes (Fig. S1e; Table 1), with PacBio HiFi reads mapping at 45.5× coverage and 99.8% mapping rate. Functional annotation was achieved for 84.0% of predicted genes (18,726 of 22,292). BUSCO completeness was 96.8% for the genome and 97.1% for proteins.

For *Plicatula muricata*, the genome was estimated to be 1,726.28 Mb, with a heterogeneity of 2.96% and a duplication of 2.42% (Fig. S1c). The chromosomal-level assembly was larger than predicted, reaching 2,207.8 Mb with 13 chromosomes (Fig. S1f; Table 1). PacBio HiFi reads mapped at 33.4× coverage with a 99.9% mapping rate. Functional annotation was achieved for 92.2% of predicted genes (25,785 of 27,966). BUSCO completeness was 95.9% for the genome and 94.9% for proteins.

#### Transposable Element Landscape

Genome assemblies of *Anomia chinensis* (530.9 Mb), *Placuna vitream* (548.9 Mb), *Heteranomia squamula* (712.4 Mb), *Plicatula muricata* (2.21 Gb), and *Pododesmus macrochisma* (766.6 Mb) revealed substantial variation in genome size among APP lineages, with *Plicatula muricata* being approximately four times larger than *Anomia chinensis* and *Placuna vitream* (Table 1). Notably, no evidence of whole-genome duplication was detected in *P. muricata* (Fig. 1b), suggesting that TE proliferation likely played a major role in this expansion, although other mechanisms cannot be entirely excluded, such as the proliferation of non-TE repetitive sequences or assembly artifacts related to high heterozygosity. To investigate the potential contribution of transposable element (TE) proliferation to this genome size disparity, we characterized the TE landscape across these genomes.

#### Genome-wide Repeat Content

Repeat annotation revealed substantial variation in TE content across the APP genomes (Table S5). Among them, *Plicatula muricata* exhibited the highest TE content (68.60%), followed by *Heteranomia squamula* (56.62%), *Placuna vitream* (50.57%), *Anomia chinensis* (43.85%), and *Pododesmus macrochisma* (38.85%). Total interspersed repeats accounted for 63.67% of the *P. muricata* genome, substantially exceeding that of the other APP species (27.34% to 46.75%). These values are comparable to or exceed those reported for many other bivalve genomes, where TEs can represent a major source of genomic variation (36). A positive correlation between genome size and TE content has been observed across eukaryotes, and TEs are widely recognized as major contributors to genome size evolution (37–39). For example, the association between TE proliferation and genome expansion has been documented in diverse metazoan lineages, including rotifers, where transposon expansion has driven a doubling in genome size (40). Our data are consistent with this general pattern across lophotrochozoans (41, 42). The substantial TE content in *Plicatula muricata* is strongly associated with its genome expansion, suggesting that TE proliferation, rather than polyploidization, is the primary driver of this expansion.

#### LINEs as the Primary Driver of Genome Expansion

Among TE categories, LINEs were the most prominent class in *Plicatula muricata*, accounting for 10.22% of the genome (225.5 Mb), substantially higher than in *Anomia chinensis* (2.49%, 13.2 Mb), *Placuna vitream* (2.37%, 13.0 Mb), *Heteranomia squamula* (4.81%, 34.3 Mb), and *Pododesmus macrochisma* (3.15%, 24.2 Mb). This finding is consistent with recent comparative analyses showing that class I elements are highly dominant in bivalve genomes, with LINE elements being the most common retroposon group (36). LINEs are non-LTR retrotransposons that replicate via target-primed reverse transcription and can significantly influence genome structure through insertional mutagenesis and ectopic recombination (43). Bivalves host an exceptional diversity of transposons compared to other molluscs, and their LINE complement may follow a "stealth drivers" model of evolution, in which multiple diversified families survive and coexist for extended periods, potentially shaping both recent and early phases of bivalve genome evolution (36). The expansion of LINEs has been implicated as a major factor driving genome size variation in diverse animal lineages, including bdelloid rotifers where LINE proliferation has contributed substantially to genome expansion (44).

Among LINE subfamilies, RTE/Bov-B elements showed the most striking expansion in *Plicatula muricata*, accounting for 2.26% of the genome (49.9 Mb), compared to only 0.02% to 0.24% in *Anomia chinensis* and *Placuna vitream*, and 1.23% to 1.41% in *Pododesmus macrochisma* and *Heteranomia squamula*, respectively. RTE/Bov-B elements are known to be particularly active in bivalve genomes and have been implicated as a major factor influencing genome expansion in other invertebrate lineages (36, 45). L2/CR1/Rex elements, which are ancient LINE clades widespread across eukaryotic genomes and often associated with insertional mutagenesis (36, 46), also showed elevated representation in *Plicatula muricata* (0.66%, 14.5 Mb) compared to *Anomia chinensis* (0.05%) and *Placuna vitream* (0.18%). Conversely, R1/LOA/Jockey elements, which are abundant in *Anomia chinensis* (1.15%, 6.08 Mb), are markedly reduced in *P. muricata* (0.13%, 2.91 Mb), suggesting differential expansion of specific LINE lineages across APP lineages. Such lineage-specific LINE dynamics have been observed in other bivalve lineages, where different LINE families have undergone independent expansions in different species, reflecting lineage-specific transposon-host interactions (36).

#### LTR Retrotransposons: A Minor Contributor

LTR retrotransposons, which are often the primary drivers of genome expansion in many large invertebrate genomes, accounted for only 2.35% of the *Plicatula muricata* genome, comparable to the other APP species (0.78% to 2.18%). This relatively low LTR content is also similar to that observed in scallops, where LTRs typically represent a modest fraction of the genome (for example, ~1% in *Mizuhopecten yessoensis*). By contrast, in the limestone mountainsnail *Oreohelix idahoensis*, LTRs comprise 57.73% of the genome, representing the most LTR-dominated repetitive landscape yet documented in a molluscan genome (47). Within LTR elements, Gypsy/DIRS1 was the most abundant family across all APP genomes, ranging from 0.29% in *Anomia chinensis* to 1.46% in *Plicatula muricata*, while Ty1/Copia and BEL/Pao elements were present at much lower proportions. This pattern is consistent with the general dominance of Gypsy elements over other LTR lineages in molluscan genomes, as observed in diverse molluscan species including oysters and gastropods (36, 47). The modest LTR content in *Plicatula muricata* suggests that, unlike many other large genomes where LTR bursts drive expansion (37, 38, 47), its expansion was not primarily driven by LTR retrotransposon activity.

**Unclassified Repeats and Lineage-Specific Expansion**

A particularly striking feature of the *Plicatula muricata* genome is the exceptionally high proportion of unclassified repeats, which account for 48.50% of the genome (1.07 Gb), substantially higher than in *Anomia chinensis* (31.85%, 169.1 Mb), *Placuna vitream* (26.05%, 143.0 Mb), *Heteranomia squamula* (31.02%, 221.0 Mb), and *Pododesmus macrochisma* (19.59%, 150.2 Mb). The unusually high proportion of unclassified repeats suggests the presence of lineage-specific repetitive elements that have not yet been characterized in existing repeat databases. This pattern is consistent with the hypothesis that recent and lineage-specific transposon activity, particularly involving LINE elements, may have driven the rapid genome expansion in *Plicatula muricata*. Indeed, previous studies have noted that unclassified repeats often represent highly diverged or lineage-specific TE families that escape detection by standard homology-based methods (48). The proliferation of unclassified repeats in *Plicatula muricata* may reflect the activity of novel or highly diverged TE families unique to this lineage, potentially representing "dark matter" of the genome—the fraction in which nothing is immediately recognizable as biologically functional but which may harbor ancient TE remnants and other repetitive sequences (36, 49, 50). This observation highlights the need for de novo TE discovery approaches to fully characterize the repetitive landscape of this species, as homology-based methods alone are insufficient for capturing the full diversity of TE content in genomes with extensive lineage-specific repeat expansions (36).

**DNA Transposons and Rolling-Circle Elements**

DNA transposons showed contrasting patterns across APP genomes. *Heteranomia squamula* exhibited the highest DNA transposon content (8.38%, 59.7 Mb), followed by *Placuna vitream* (7.35%, 40.3 Mb), while *Plicatula muricata* (2.33%, 51.4 Mb) and *Anomia chinensis* (3.02%, 16.0 Mb) showed lower proportions. Among DNA transposon families, hobo-Activator elements were most abundant in *Anomia chinensis* (1.00%), while Tc1-IS630-Pogo showed elevated representation in *Plicatula muricata* (0.92%, 20.3 Mb) and Tourist/Harbinger elements were notably expanded in *Heteranomia squamula* (0.67%, 4.81 Mb). Rolling-circle (RC) elements (Helitrons), which replicate via a rolling-circle mechanism rather than the classical cut-and-paste mechanism of other DNA transposons (51, 52), also varied considerably, with *Placuna vitream* (7.95%, 43.6 Mb) and *Heteranomia squamula* (8.34%, 59.4 Mb) showing the highest proportions, while *Anomia chinensis* (0.01%) exhibited a near-complete absence of this TE class. Helitrons are widespread in eukaryotic genomes and have been shown to capture and duplicate gene fragments, contributing to genome evolution and innovation (51, 52). The differential representation of DNA transposons and RC elements across APP genomes suggests that distinct TE classes have undergone lineage-specific expansions, potentially reflecting differences in host defense mechanisms or recombination landscapes among these species (50). The near absence of Helitrons in *Anomia chinensis* is particularly noteworthy, as it suggests either a lineage-specific loss of this TE class or a failure of Helitrons to invade this genome, possibly due to differences in host defense mechanisms or genomic architecture (50).

**Comparison with the relatives**

To contextualize these findings, we compared the TE profiles of APP genomes with those of other Pectinida species. The scallop *Mizuhopecten yessoensis* exhibited the lowest TE content (30.85%), with moderate LINE (3.88%) and LTR (0.99%) content. *Catillopecten margaritatus* showed higher TE content (51.27%), with LINEs at 6.40% and LTRs at 1.01%. The limid *Ctenoides ales*, whose genome is substantially larger than other pteriomorphian genomes largely due to a substantial number of TEs (53), exhibited the highest LINE content among the comparison species (10.29%), comparable to *Plicatula muricata*, along with substantial DNA transposons (10.10%) and rolling-circle elements (17.22%). Notably, the TE profile of *Plicatula muricata*, characterized by high LINE and unclassified repeat content, is distinct from that of other large bivalve genomes where LTRs often dominate the repetitive landscape. For instance, in the limestone mountainsnail *Oreohelix idahoensis*, LTRs account for the vast majority of the repetitive content (57.73%) (47), whereas in *P. muricata*, LTRs contribute only 2.35%. This difference suggests that distinct TE classes can serve as primary drivers of genome expansion in different lineages, even among closely related taxa (38, 39). The diversity of TE landscapes across Pectinida highlights the evolutionary lability of repetitive element dynamics within this order and underscores the importance of lineage-specific factors in shaping genome architecture (36).

**Implications for Genome Evolution in the APP Lineage**

The stark contrast in TE composition between *Plicatula muricata* and its APP relatives highlights the dynamic nature of repeat evolution within this clade. The proliferation of LINEs and unclassified repeats in *P. muricata*, rather than LTR retrotransposons, suggests that different TE classes can serve as primary drivers of genome expansion in different lineages, even among closely related species. This observation is consistent with the emerging view that bivalves host an exceptional diversity of transposons compared to other molluscs, and that multiple diversified transposon lineages contribute to both early and recent bivalve genome evolution (36). The potential relationship between TE dynamics and the extensive chromosomal fusions documented in the APP lineage warrants consideration. It remains unclear whether TE proliferation preceded or followed the chromosomal fusion events that dramatically reduced chromosome numbers in this clade. One possibility is that TE accumulation contributed to increased recombination or chromosomal breakage, facilitating fusion events (L1 insertion intermediates can recombine with distal DNA breaks to generate chromosomal rearrangements) (54). Alternatively, chromosomal fusions may have created conditions, such as reduced recombination rates in fused regions or altered chromatin environments, that favored TE proliferation (transposable elements tend to accumulate in regions with low levels of recombination) (55). TE activity has been shown to drive structural variation in bivalves, with up to 14% of oyster genome base pairs in a hemizygous state attributable to TE-mediated structural variants (42). The exceptionally high unclassified repeat content in *P. muricata* may also be linked to its large genome size and the presence of regions refractory to standard repeat annotation, potentially including ancient TE remnants that have accumulated following chromosomal rearrangements. However, establishing causal relationships between these processes will require more detailed analyses, including examination of TE insertion sites relative to fusion breakpoints and investigation of the timing of TE expansion relative to chromosomal fusion events.

**Gene Family Expansion in Anomiida**

To investigate whether the TE landscape and progressive chromosomal fusions of Anomiida are reflected in nuclear gene family dynamics, we performed comparative gene family expansion and contraction analysis using CAFE5 v5.0 (56) across four Anomiida species with chromosome-level genome assemblies (*Plicatula muricata*, *Placuna vitream*, *Anomia chinensis*, and *Heteranomia squamula*), with two scallop species (*Mizuhopecten yessoensis* and *Ctenoides ales*) serving as outgroups. *Pododesmus macrochisma* was excluded from this analysis due to the lack of a chromosome-level genome assembly. CAFE5 analysis identified nine gene families showing significant expansion across multiple Anomiida species (defined as expansion in at least three of the four species, $p < 0.05$), suggesting possible early expansion events in the Anomiida ancestor or recurrent expansions in independent lineages. No gene families showed significant contraction at this node, indicating that early Anomiida evolution was characterized by gene proliferation rather than large-scale gene loss. The nine expanded families fall into three functional categories: (1) DNA repair and genome stability (OG0000082, OG0000109, OG0000537), (2) transposable element-related (OG0000161, OG0000387, OG0000808), and (3) other functions (OG0000044, OG0000056, OG0000259) (Table S9).

**Association with Transposable Element Content**

The expansion of specific gene families in the APP lineage shows a striking association with transposable element content, suggesting that TE proliferation may have driven compensatory responses in genes involved in genome maintenance. Among the most notable expansions is OG0000082, encoding TatD DNases, conserved metal-dependent nucleases involved in DNA repair, RNA processing, and R-loop resolution (57). This gene family exhibits the most dramatic expansion in *Plicatula muricata* (82 copies) and *Heteranomia squamula* (78 copies), the two species with the highest TE loads (68.60% and 56.62%, respectively), compared to only 11–34 copies in *Anomia* *chinensis* and *Placuna vitream*, and 7–12 copies in scallops (Table S9). TatD DNases function as endonucleases that cleave DNA and RNA, playing critical roles in DNA repair pathways, particularly in the resolution of R-loops, which are three-stranded nucleic acid structures that form during transcription (57). R-loops are known to be stabilized by TE insertions and can cause genomic instability if not properly resolved (58). The expansion of TatD DNases in TE-rich genomes may therefore represent a compensatory response to the increased burden of R-loops and DNA damage induced by TE activity. Similarly, OG0000109, encoding PIF1 helicases, shows extreme expansion in *Plicatula muricata* (57 copies), which harbors massive LINE proliferation (10.22%, 225.5 Mb), but is absent in *Placuna vitream* (0 copies), despite its moderate TE load (50.57%) (Table S9). PIF1 helicases are multifunctional proteins that unwind G-quadruplex (G4) DNA structures, which are non-canonical secondary structures formed by guanine-rich sequences (59, 60). G4 structures are enriched in TE-rich genomes, particularly in the regulatory regions of LINE and LTR retrotransposons, and can cause replication fork stalling and genomic instability if not resolved (61, 62). PIF1 helicases are known to be the primary enzymes responsible for resolving G4 structures in eukaryotic cells, and their overexpression has been shown to suppress G4-induced genome instability (60, 63). The complete absence of PIF1 expansion in *Placuna vitream* can be explained by its TE landscape, which is dominated by DNA transposons (7.35%) and rolling-circle elements (7.95%) rather than LINEs—the latter being the primary source of G4 structures. In *Plicatula muricata*, the high LINE content (10.22%) likely generates abundant G4 structures, creating strong selective pressure for PIF1 expansion. This differential pattern provides compelling correlative evidence that TE family composition, not merely total TE content, determines the host's compensatory gene family expansion response.

Additionally, OG0000161 (encoding P-element transposases) and OG0000387 (containing ASB3, TatD DNase, and retroviral gag-asp protease domains) both show extreme expansion in *Plicatula muricata* (81 and 66 copies, respectively) while being completely absent in *Placuna vitream*. Notably, *Placuna vitream* has the highest DNA transposon content among APP species (7.35%), yet lacks OG0000161 entirely, suggesting that transposase gene expansion is not simply a consequence of DNA transposon abundance but may reflect lineage-specific differences in transposon family composition or host-transposon interaction dynamics. The mosaic structure of OG0000387—containing both nuclease and retroviral domains—suggests that this gene family may have originated from a retrotransposon and been subsequently domesticated for host functions (64, 65). The expansion patterns of these gene families parallel those of TatD DNases and PIF1 helicases, further reinforcing the association between TE load and the expansion of genes involved in genome maintenance.

**Association with Chromosomal Fusion History**

When mapped onto the chromosomal fusion history of the APP lineage (Table 2), the expansion patterns of gene families exhibit an association with the inferred phylogenetic nodes where fusion events occurred. From the ancestral 20 molluscan linkage groups (MLGs) (66), the Anomiida ancestor underwent five independent fusion events, reducing chromosome number from 20 to 15. This ancestral fusion phase was followed by lineage-specific fusion events that ultimately reduced chromosome numbers to as few as six in *Heteranomia squamula* (Table 2). Notably, the expansion of OG0000082 and OG0000387 appears to have occurred at or after the lineage-specific fusion events leading to *Plicatula muricata* and *Heteranomia squamula*, as these species exhibit the highest copy numbers and have undergone the most extensive post-ancestral fusions (Table 2). Chromosomal fusions can generate dicentric chromosomes, leading to breakage-fusion-bridge cycles that produce additional rearrangements and DNA damage (67). The fusion events are associated with increased genomic instability and selection for genome maintenance pathways (53, 67). The expansion of TatD DNases and the multi-domain OG0000387 in species with the most extensive fusion histories may therefore have been driven by selection to mitigate the DNA damage and genomic instability caused by fusion-induced breakage-fusion-bridge cycles.

Importantly, there is no simple linear correlation between chromosome number and DNA repair gene copy numbers. *Placuna vitream* (8 chromosomes) has only 11 copies of OG0000082, while *Anomia chinensis* (7 chromosomes) has 34 copies. *Heteranomia squamula* (6 chromosomes, 78 copies) and *Plicatula muricata* (13 chromosomes, 82 copies) show similar copy numbers despite vastly different chromosome numbers (Table S9). This suggests that the fusion events themselves, rather than the final chromosome number, are associated with repair gene expansion. Species experiencing more fusion events or more severe breakage-fusion-bridge cycles would face stronger selection for repair gene expansion, but this is not directly reflected in chromosome number. The divergence between *Anomia chinensis* and *Placuna vitream*, despite similar fusion counts (13 vs. 12) and chromosome numbers (7 vs. 8), further suggests that factors beyond fusion count, such as TE content and composition, may influence the host response.

Collectively, TE content shows a more consistent association with DNA repair gene copy numbers than chromosome number. *Plicatula muricata* (68.60% TEs) and *Heteranomia squamula* (56.62% TEs) have the highest TE loads and the highest OG0000082 and OG0000387 copy numbers. *Anomia chinensis* (43.85% TEs) and *Placuna vitream* (50.57% TEs) show intermediate TE loads but divergent repair gene copy numbers (34 vs. 11), suggesting that TE family composition influences the host response. In *Placuna vitream*, the TE landscape is dominated by DNA transposons (7.35%) and rolling-circle elements (7.95%), whereas *Plicatula muricata* has a much higher LINE content (10.22%) (Table S5). LINEs are known to be particularly mutagenic, as they can cause insertions, deletions, and chromosomal rearrangements through non-homologous end joining and homologous recombination (36, 43). The high LINE content in *Plicatula muricata* may therefore exert stronger selection pressure for DNA repair gene expansion than the DNA transposon-dominated landscape of *Placuna vitream*. These patterns suggest that both TE proliferation and chromosomal fusions have contributed to the expansion of genome maintenance genes in the APP lineage, but TE-driven selection appears to be the more consistent and stronger driver. The association between TE content and DNA repair gene copy numbers, together with the correlation of these expansions with the timing of chromosomal fusion events, suggests that genomic instability arising from TE activity and chromosome restructuring may have synergistically shaped the evolution of genome maintenance pathways. It remains unclear, however, whether TE proliferation is the cause or the consequence of these genomic changes. The strong correlations we observe provide a foundation for future functional studies to disentangle these relationships. The expansion of TatD DNases, PIF1 helicases, and transposase-derived genes in TE-rich APP species likely represents a compensatory response to increased genomic instability, enabling these species to maintain genome integrity despite high TE loads and extensive chromosomal rearrangements.

#### Reference

1. J. Feng, *et al.*, Novel gene rearrangement in the mitochondrial genome of *Siliqua minima* (Bivalvia, Adapedonta) and phylogenetic implications for Imparidentia. *PLoS ONE* **16**, e0249446 (2021).
2. F. Ghiselli, *et al.*, Molluscan mitochondrial genomes break the rules. *Philos. Trans. R. Soc. B Biol. Sci.* **376**, 20200159 (2021).
3. Y. Zhang, *et al.*, Comparative mitogenomic analyses of the infraclass Pteriomorphia (Mollusca: Bivalvia) provides novel insights into gene rearrangement and phylogeny. *Comp. Biochem. Physiol. Part D Genomics Proteomics* **53**, 101361 (2025).
4. J. Ren, X. Shen, F. Jiang, et al., The mitochondrial genomes of two scallops, *Argopecten irradians* and *Chlamys farreri* (Mollusca: Bivalvia): The most highly rearranged gene order in the family Pectinidae. *J. Mol. Evol.* **70**, 57–68 (2010).
5. A. Marín, T. Fujimoto, K. Arai, The mitochondrial genomes of Pecten albicans and *Pecten maximus* (Bivalvia: Pectinidae) reveal a novel gene arrangement with low genetic differentiation. *Biochem. Syst. Ecol.* **61**, 208–217 (2015).
6. T. Malkócs, *et al.*, Complex mitogenomic rearrangements within the Pectinidae (Mollusca: Bivalvia). *BMC Ecol. Evol.* **22**, 29 (2022).
7. T. A. Rawlings, M. J. MacInnis, R. Bieler, J. L. Boore, T. M. Collins, Sessile snails, dynamic genomes: Gene rearrangements within the mitochondrial genome of a family of caenogastropod molluscs. *BMC Genomics* **11**, 440 (2010).
8. Z. Yu, Z. Wei, X. Kong, W. Shi, Complete mitochondrial DNA sequence of oyster *Crassostrea hongkongensis* – a case of “tandem duplication-random loss” for genome rearrangement in Crassostrea? *BMC Genomics* **9**, 477 (2008).
9. X. Wu, *et al.*, Evolution of the tRNA gene family in mitochondrial genomes of five Meretrix clams (Bivalvia, Veneridae). *Gene* **533**, 439–446 (2014).
10. T. A. Rawlings, T. M. Collins, R. Bieler, Changing identities: tRNA duplication and remolding within animal mitochondrial genomes. *Proc. Natl. Acad. Sci. U. S. A.* **100**, 15700–15705 (2003).
11. M. Passamonti, F. Ghiselli, Doubly uniparental inheritance: Two mitochondrial genomes, one precious model for organelle DNA inheritance and evolution. *DNA Cell Biol.* **28**, 79–89 (2009).
12. E. Zouros, Biparental inheritance through uniparental transmission: The doubly uniparental inheritance (DUI) of mitochondrial DNA. *Evol. Biol.* **40**, 1–31 (2013).
13. F. Ghiselli, *et al.*, Structure, transcription, and variability of metazoan mitochondrial genome: Perspectives from an unusual mitochondrial inheritance system. *Genome Biol. Evol.* **5**, 1735–1754 (2013).
14. A. Gusman, S. Lecomte, D. T. Stewart, M. Passamonti, S. Breton, Pursuing the quest for better understanding the taxonomic distribution of the system of doubly uniparental inheritance of mtDNA. *PeerJ* **4**, e2760 (2016).
15. S. Breton, H. D. Beaupré, D. T. Stewart, W. R. Hoeh, P. U. Blier, The unusual system of doubly uniparental inheritance of mtDNA: Isn’t one enough? *Trends Genet.* **23**, 465–474 (2007).

16. J. M. Serb, C. Lydeard, Complete mtDNA sequence of the North American freshwater mussel, Lampsilis ornata (Unionidae): An examination of the evolution and phylogenetic utility of mitochondrial genome organization in Bivalvia (Mollusca). *Mol. Biol. Evol.* **20**, 1854–1866 (2003).
17. A. Mizi, E. Zouros, N. Moschonas, G. C. Rodakis, The complete maternal and paternal mitochondrial genomes of the mediterranean mussel M*ytilus galloprovincialis*: Implications for the doubly uniparental inheritance mode of mtDNA. *Mol. Biol. Evol.* **22**, 952–967 (2005).
18. B. Smietanka, A. Burzyński, R. Wenne, Comparative genomics of marine mussels (*Mytilus* spp.) gender associated mtDNA: Rapidly evolving atp8. *J. Mol. Evol.* **76**, 123–134 (2013).
19. S. Osawa, T. Ohama, T. H. Jukes, K. Watanabe, Evolution of the mitochondrial genetic code. I. Origin of AGR serine and stop codons in metazoan mitochondria. *J. Mol. Evol.* **29**, 202–207 (1989).
20. R. D. Knight, S. J. Freeland, L. F. Landweber, Rewiring the keyboard: Evolvability of the genetic code. *Nat. Rev. Genet.* **2**, 49–58 (2001).
21. D. W. Schultz, M. Yarus, Transfer RNA mutation and the malleability of the genetic code. *J. Mol. Biol.* **235**, 1377–1380 (1994).
22. S. Sengupta, P. G. Higgs, A unified model of codon reassignment in alternative genetic codes. *Genetics* **170**, 831–840 (2005).
23. Y. Li, *et al.*, Mitogenomics reveals a novel genetic code in Hemichordata. *Genome Biol. Evol.* **11**, 29–40 (2019).
24. Y.-T. Lin, J.-W. Qiu, Reassessment of Pectinida (Mollusca: Bivalvia) phylogenetic relationships and description of a new *Parvamussium* species. *Zool. J. Linn. Soc.* **206**, zlaf200 (2026).
25. Y. X. Li, *et al.*, Phylogenomics of Bivalvia using ultraconserved elements reveal new topologies for Pteriomorphia and Imparidentia. *Syst. Biol.* **74**, 16–33 (2025).
26. L. R. L. Simone, V. S. do Amaral, Phenotypic features of *Dimya cf. japonica* (Bivalvia, Dimyidae) from Niue Island (South Pacific) with accounts on its phylogeny and taxonomic relationships. *Malacologia* **64**, 121–136 (2021).
27. M. Hautmann, Taxonomy and phylogeny of cementing Triassic bivalves (families Prospondylidae, Plicatulidae, Dimyidae and Ostreidae). *Palaeontology* **44**, 339–373 (2001).
28. C. M. Yonge, The status of the Plicatulidae and the Dimyidae in relation to the superfamily Pectinacea (Mollusca: Bivalvia). *J. Zool.* **176**, 545–553 (1975).
29. C. M. Yonge, Functional morphology with particular reference to hinge and ligament in *Spondylus* and *Plicatula* and a discussion on relations within the superfamily Pectinacea (Mollusca: Bivalvia). *Philos. Trans. R. Soc. Lond. B Biol. Sci.* **267**, 173–208 (1973).
30. H. H. Dijkstra, P. Maestrati, Pectinoidea (Mollusca, Bivalvia, Propeamussiidae, Cyclochlamydidae n. fam., Entoliidae and Pectinidae) from the Vanuatu Archipelago. *Zoosystema* **34**, 389–408 (2012).
31. C. M. Yonge, Form and evolution in the Anomiacea (Mollusca: Bivalvia)—*Pododesmus*, *Anomia*, *Patro*, *Enigmonia* (Anomiidae); *Placunanomia*, *Placuna*

(Placunidae fam. nov.). *Philos. Trans. R. Soc. Lond. B Biol. Sci.* **276**, 453–523 (1977).

32. P. M. Mikkelsen, R. Bieler, *Seashells of Southern Florida: Bivalves* (Princeton University Press, 2008).
33. T. R. Waller, Morphology, phylogeny, and systematic revision of genera in the Dimyidae (Mollusca, Bivalvia, Pteriomorphia). *J. Paleontol.* **86**, 829–851 (2012).
34. J.-H. Li, M. Hautmann, Q.-Q. Zhang, H.-C. Zhang, J.-G. Sha, Iteration in the evolution of hinge teeth in the cementing bivalve families Prospondylidae and Plicatulidae: Evidence from Persia (Nyalamia) n. subgen. from the lowest Jurassic of Xizang (Tibet). *Palaeoworld* **35**, 201106 (2026).
35. C. M. Yonge, On the Dimyidae (Mollusca: Bivalvia) with special reference to *Dimya corrugata* Hedley and *Basiliomya goreaui* Bayer. *J. Molluscan Stud.* **44**, 357–375 (1978).
36. J. Martelossi, *et al.*, Multiple and diversified transposon lineages contribute to early and recent bivalve genome evolution. *BMC Biol.* **21**, 145 (2023).
37. S. Liu, *et al.*, The origins and functional significance of bivalve genome diversity. *bioRxiv* 611967 (2024). https://doi.org/10.1101/2024.10.21.611967.
38. S. Farhat, *et al.*, Comparative analysis of the *Mercenaria mercenaria* genome provides insights into the diversity of transposable elements and immune molecules in bivalve mollusks. *BMC Genomics* **23**, 192 (2022).
39. M. G. Kidwell, Transposable elements and the evolution of genome size in eukaryotes. *Genetica* **115**, 49–63 (2002).
40. J. Blommaert, S. Riss, B. Hecox-Lea, D. B. Mark Welch, C. P. Stelzer, Small, but surprisingly repetitive genomes: Transposon expansion and not polyploidy has driven a doubling in genome size in a metazoan species complex. *BMC Genomics* **20**, 466 (2019).
41. O. Simakov, *et al.*, Deeply conserved synteny and the evolution of metazoan chromosomes. *Sci. Adv.* **8**, eabi5884 (2022).
42. J. Martelossi, A. Luchetti, A. Suh, F. Ghiselli, V. Peona, Transposable element activity and polymorphisms drive structural variability within and between individuals in bivalves. *bioRxiv* 705023 (2026). https://doi.org/10.1101/2026.01.15.705023.
43. H. H. Kazazian, Mobile elements: Drivers of genome evolution. *Science* **303**, 1626–1632 (2004).
44. J.-F. Flot, *et al.*, Genomic evidence for ameiotic evolution in the bdelloid rotifer *Adineta vaga*. *Nature* **500**, 453–457 (2013).
45. K. K. Kojima, H. Fujiwara, Long-term inheritance of the *28S rDNA*-specific retrotransposon R2. *Mol. Biol. Evol.* **22**, 2157–2165 (2005).
46. V. V. Kapitonov, J. Jurka, Molecular paleontology of transposable elements in the *Drosophila melanogaster* genome. *Proc. Natl. Acad. Sci. U. S. A.* **100**, 6569–6574 (2003).
47. T. M. Linscott, A. González-González, T. Hirano, C. E. Parent, *De novo* genome assembly and genome skims reveal LTRs dominate the genome of a limestone endemic Mountainsnail (*Oreohelix idahoensis*). *BMC Genomics* **23**, 796 (2022).

48. J. S. Sproul, *et al.*, Analyses of 600+ insect genomes reveal repetitive element dynamics and highlight biodiversity-scale repeat annotation challenges. *Genome Res.* **33**, 1708–1717 (2023).
49. F. Maumus, H. Quesneville, Deep investigation of Arabidopsis thaliana junk DNA reveals a continuum between repetitive elements and genomic dark matter. *PLoS ONE* **9**, e94101 (2014).
50. R. K. Slotkin, R. Martienssen, Transposable elements and the epigenetic regulation of the genome. *Nat. Rev. Genet.* **8**, 272–285 (2007).
51. V. V. Kapitonov, J. Jurka, Rolling-circle transposons in eukaryotes. *Proc. Natl. Acad. Sci. U. S. A.* **98**, 8714–8719 (2001).
52. J. Thomas, E. J. Pritham, Helitrons, the eukaryotic rolling-circle transposable elements. *Microbiol. Spectr.* **3**, MDNA3-0049–2014 (2015).
53. K. E. McElroy, R. Masonbrink, S. Chudalayandi, A. J. Severin, J. M. Serb, A chromosome-level genome assembly of the disco clam, *Ctenoides ales*. *G3 Genes Genomes Genet.* **14**, jkae115 (2024).
54. Y. Sun, et al., L1 insertion intermediates recombine with one another or with DNA breaks to form genome rearrangements. *bioRxiv* 676864 (2025). https://doi.org/10.1101/2025.01.10.676864.
55. T. V. Kent, J. Uzunović, S. I. Wright, Coevolution between transposable elements and recombination. *Philos. Trans. R. Soc. B Biol. Sci.* **372**, 20160458 (2017).
56. F. K. Mendes, D. Vanderpool, B. Fulton, M. W. Hahn, CAFE 5 models variation in evolutionary rates among gene families. *Bioinformatics* **36**, 5516–5518 (2020).
57. L. Balakrishnan, R. A. Bambara, Flap endonuclease 1. *Annu. Rev. Biochem.* **82**, 119–138 (2013).
58. A. Aguilera, T. García-Muse, R loops: From transcription byproducts to threats to genome stability. *Mol. Cell* **46**, 115–124 (2012).
59. M. L. Bochman, K. Paeschke, V. A. Zakian, DNA secondary structures: Stability and function of G-quadruplex structures. *Nat. Rev. Genet.* **13**, 770–780 (2012).
60. K. Paeschke, *et al.*, Pif1 family helicases suppress genome instability at G-quadruplex motifs. *Nature* **497**, 458–462 (2013).
61. P. Sarkies, C. Reams, L. J. Simpson, J. E. Sale, Epigenetic instability due to defective replication of structured DNA. *Mol. Cell* **40**, 703–713 (2010).
62. D. Rhodes, H. J. Lipps, G-quadruplexes and their regulatory roles in biology. *Nucleic Acids Res.* **43**, 8627–8637 (2015).
63. J. B. Vannier, V. Pavicic-Kaltenbrunner, M. I. Petalcorin, H. Ding, S. J. Boulton, RTEL1 dismantles T loops and counteracts telomeric G4-DNA to maintain telomere integrity. *Cell* **149**, 795–806 (2012).
64. E. R. Havecker, X. Gao, D. F. Voytas, The diversity of LTR retrotransposons. *Genome Biol.* **5**, 225 (2004).
65. J. N. Volff, Turning junk into gold: Domestication of transposable elements and the creation of new genes in eukaryotes. *BioEssays* **28**, 913–922 (2006).
66. J. D. Sigwart, Y. Li, Z. Chen, K. Vončina, J. Sun, Still waters run deep in large-scale genome rearrangements of morphologically conservative Polyplacophora. *eLife* **13**, RP10254 (2025).

67. B. McClintock, The stability of broken ends of chromosomes in Zea mays. *Genetics* **26**, 234–282 (1941).

### Supplementary Figures

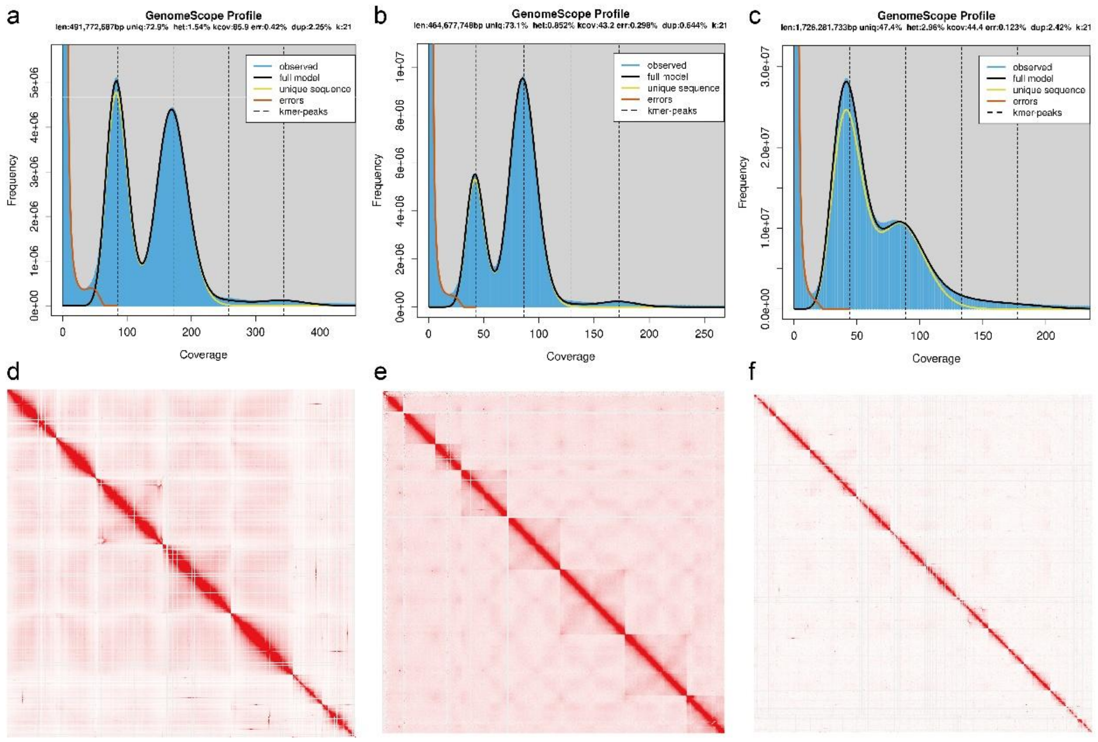


**Fig. S1** Estimation of genome size and inferred chromosome numbers. (a–c) Genome size estimation and heterozygosity profiles for *Anomia chinensis*, *Placuna vitream*, and *Plicatula muricata*, respectively, based on *k*-mer frequency analysis. (d–f) Hi-C interaction heatmaps showing the inferred chromosome-level assemblies for the three species, with each heatmap representing the contact frequency between genomic regions.

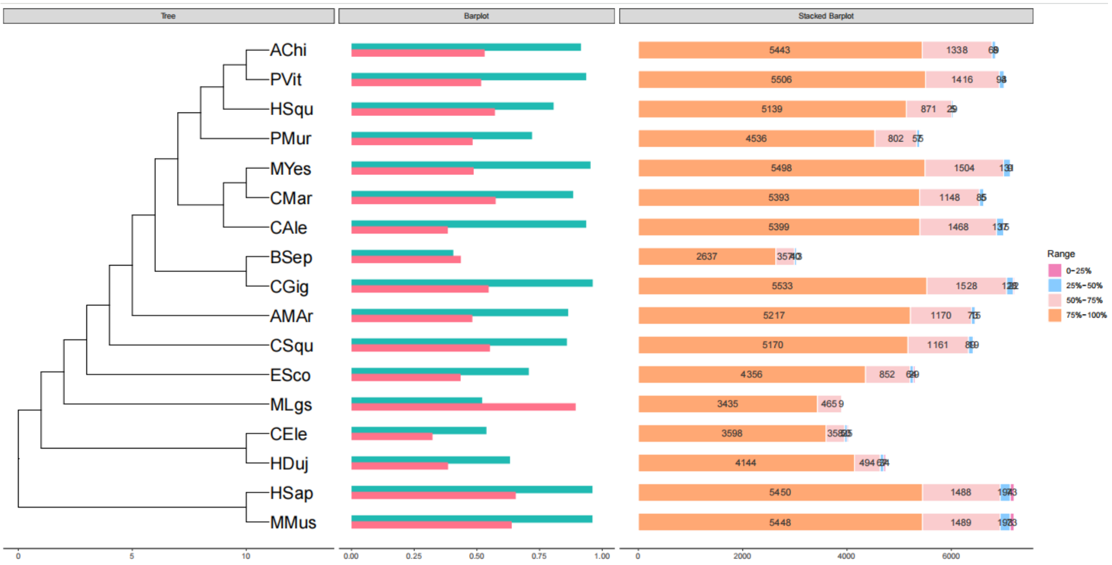


**Fig. S2** Ancestral gene families in the examined genomes. The tree shows the phylogenomic relationships among the species used to reconstruct ancestral gene families. The middle bar plot shows the percentage (x-axis) of ancestral gene families for each species, divided by the total number of ancestral gene families (green bars) or the total number of gene families of a given species (red bars). The stacked histogram shows the number (x-axis) of ancestral gene families in each species, with different colour bars representing the conservation degree (falling within four quantiles) of ancestral gene families in all assayed species.

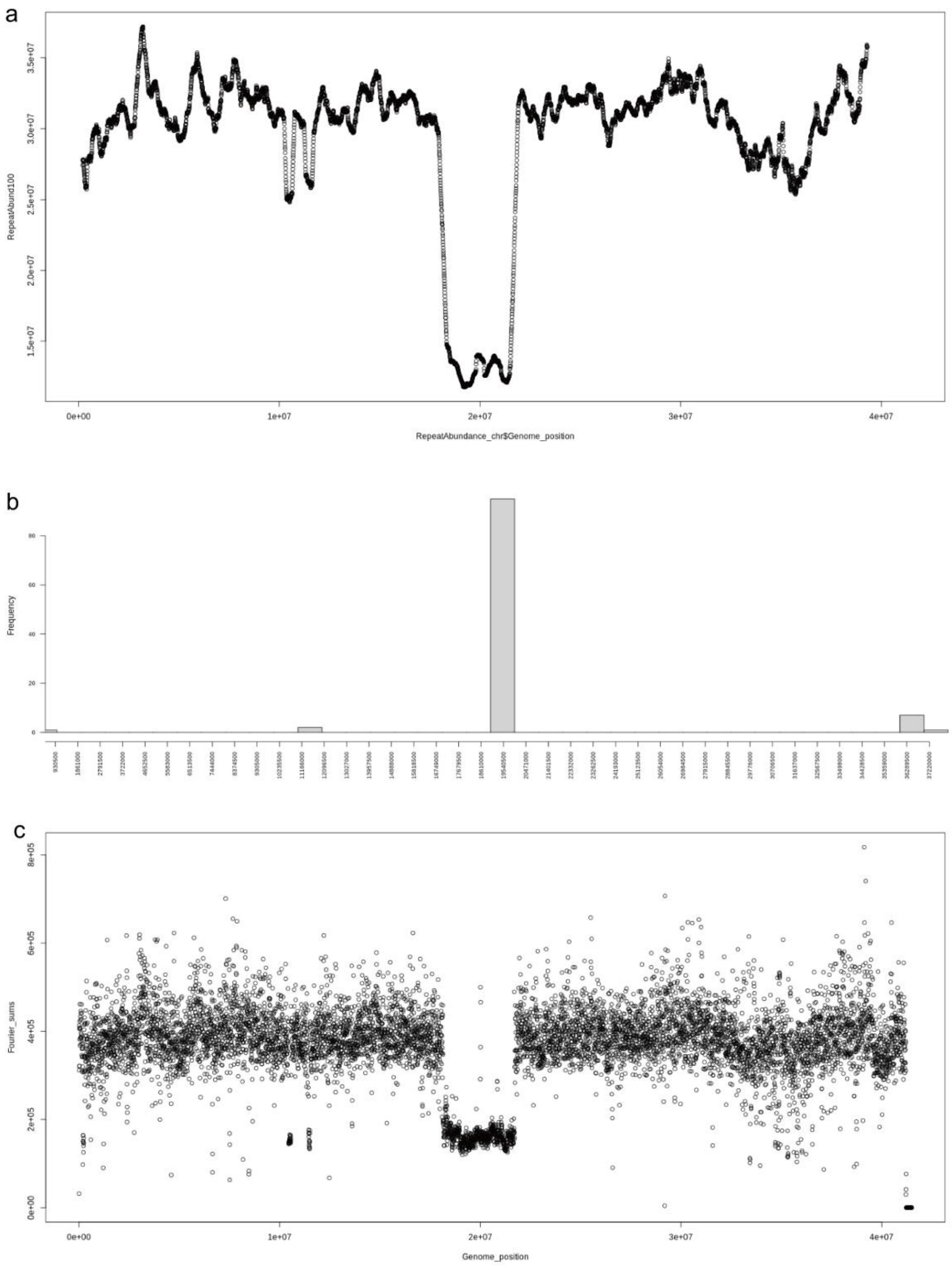


**Fig. S3** RepeatOBserver analysis of centromeric regions on chromosome 7 of *Placuna vitream*. (a) Rolling mean of repeat abundance diversity (window size = 80 bp). (b) Sequence value histogram (POWER_SUM) showing the distribution of a specific repeat family along the chromosome. (c) Distribution of total repeat abundance (sum of all repeat families).

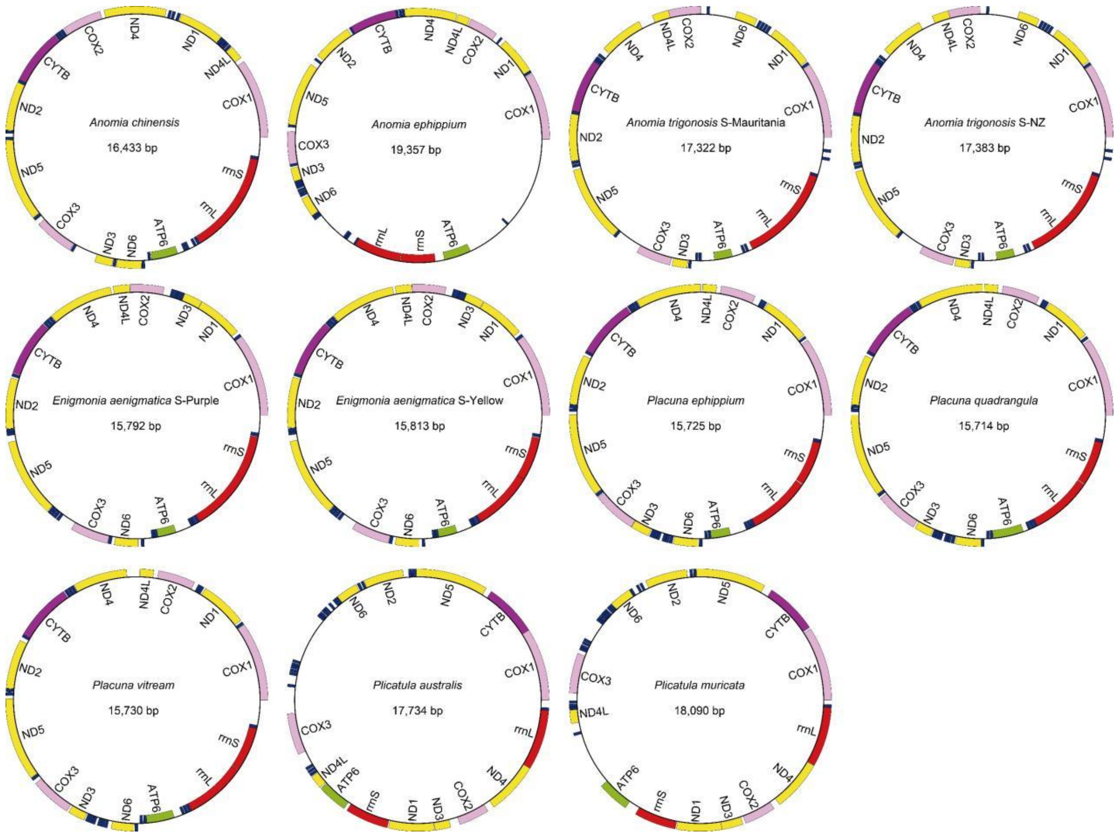


**Fig. S4** Mitochondrial genome maps of species employing the newly identified genetic code with TAA reassigned to Tyr compared to the code 5. For each species, the species name and mitogenome length are indicated at the center of the circle. Genes are color-coded according to functional categories: ATP synthases (ATP6, ATP8), cytochrome c oxidases (COX1–3), NADH dehydrogenases (ND1–6 and ND4L), cytochrome b (CYTB), transfer RNA genes (tRNAs), and ribosomal RNA genes (rRNAs).

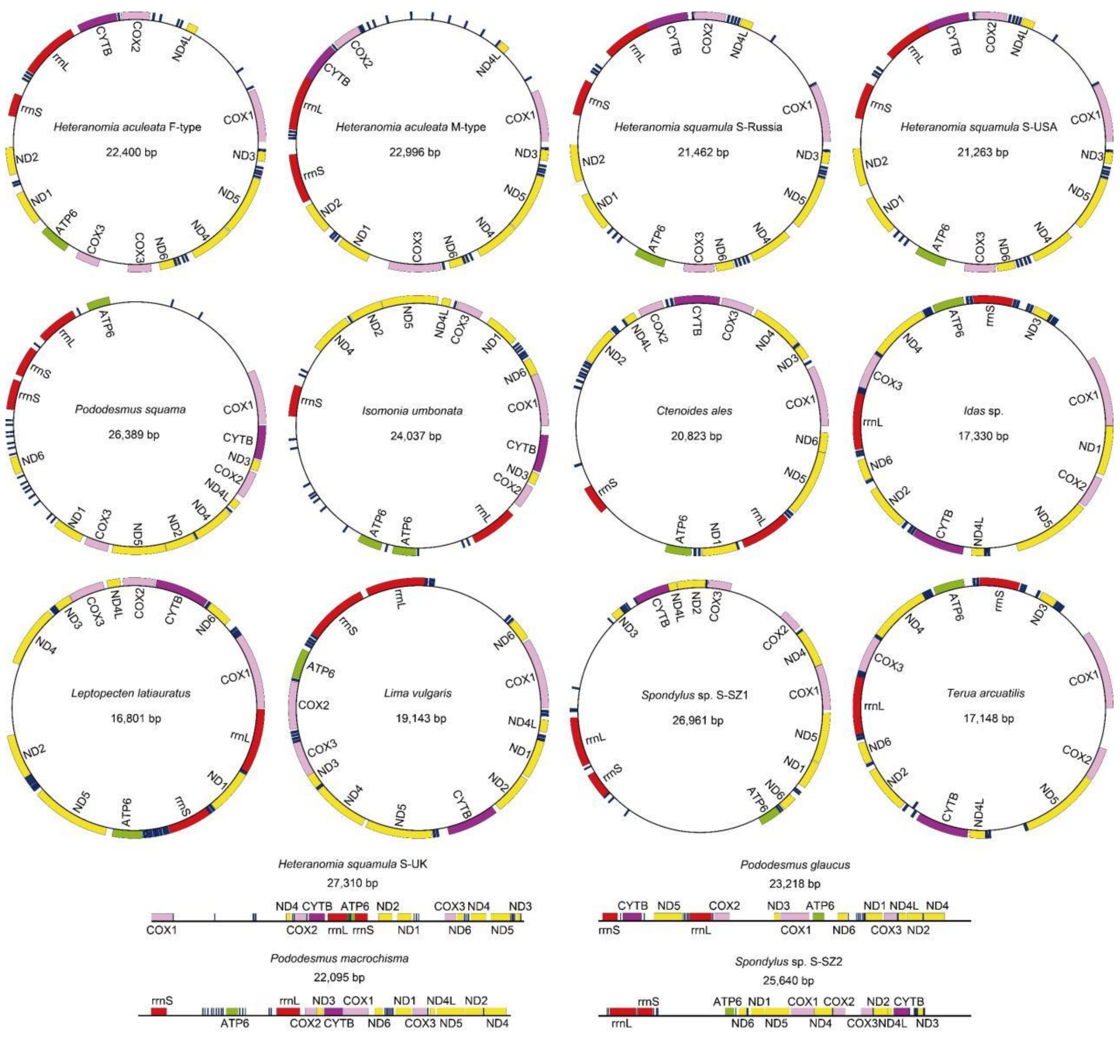


**Fig. S5** Mitochondrial genome maps of species using code 5. For each species, the species name and mitogenome length are indicated at the center of the circle. Genes are color-coded according to functional categories: ATP synthases (ATP6, ATP8), cytochrome c oxidases (COX1–3), NADH dehydrogenases (ND1–6 and ND4L), cytochrome b (CYTB), transfer RNA genes (tRNAs), and ribosomal RNA genes (rRNAs). Uncircular mitogenomes are presented in linearized form.

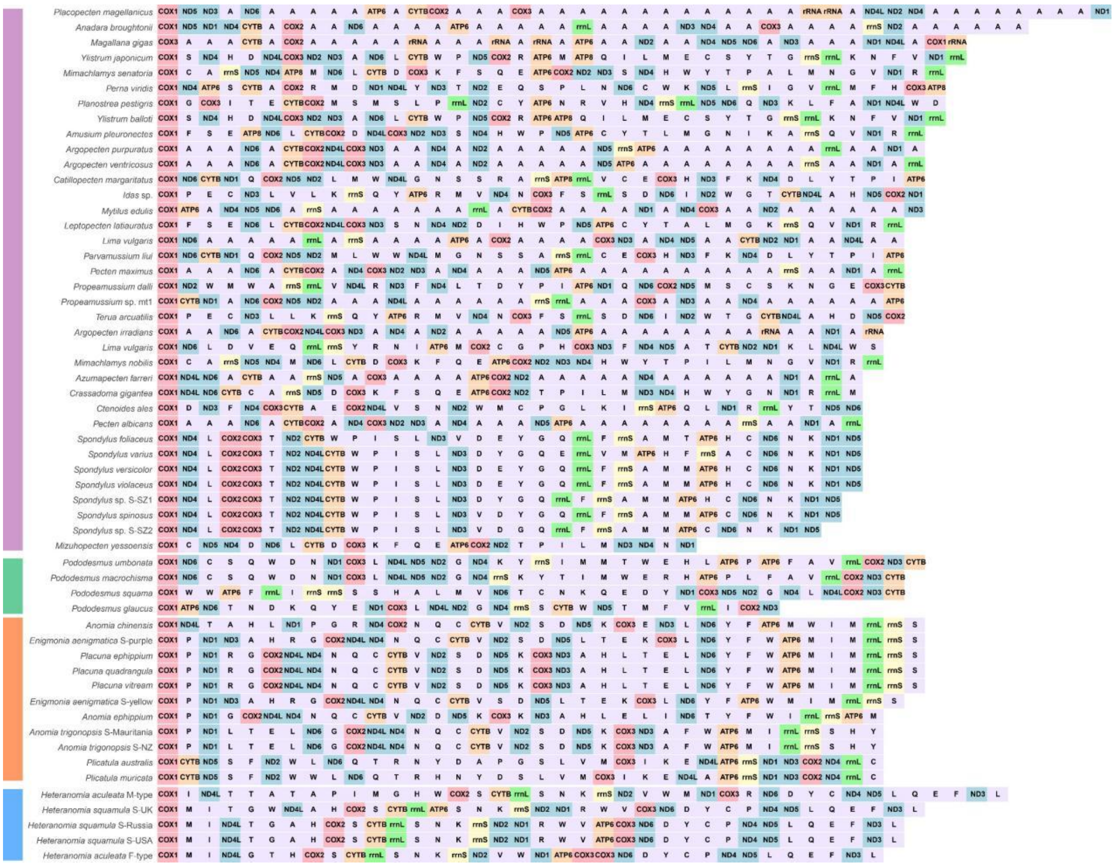

**Fig. S6** Comparison of mitochondrial gene orders among species with different configurations (TAA=Tyr, TAG+1 frameshift, code 5, and Pododesmus-type). Each colored block represents a gene, with colors indicating functional categories: ATP synthases (ATP6, ATP8), cytochrome c oxidases (COX1–3), NADH dehydrogenases (ND1–6 and ND4L), cytochrome b (CYTB), transfer RNA genes (tRNAs), and ribosomal RNA genes (rRNAs). Block lengths do not correspond to gene lengths, and gene orientations are not indicated.

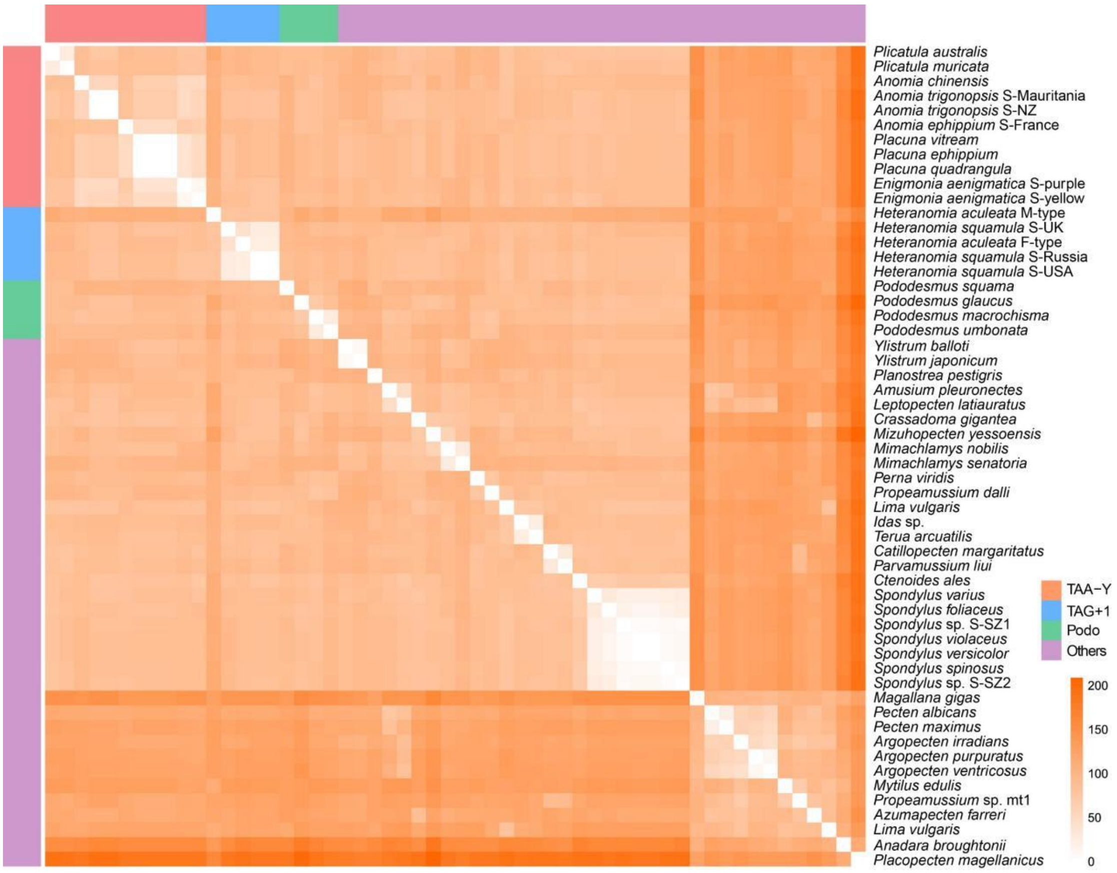


**Fig. S7 Heatmap of pairwise mitochondrial gene-order distances among the** analyzed species. The color gradient represents the degree of gene-order dissimilarity, with warmer colors indicating greater distances and cooler colors indicating greater similarity. Species are grouped according to their mitochondrial genome configurations: TAA=Tyr lineages, TAG+1 frameshift lineages, Pododesmus-type, and code 5 lineages. The gene order distance was calculated as the number of gene rearrangement steps (or breakpoints) between each pair of genomes. This heatmap illustrates the marked structural divergence among major mitochondrial configurations.

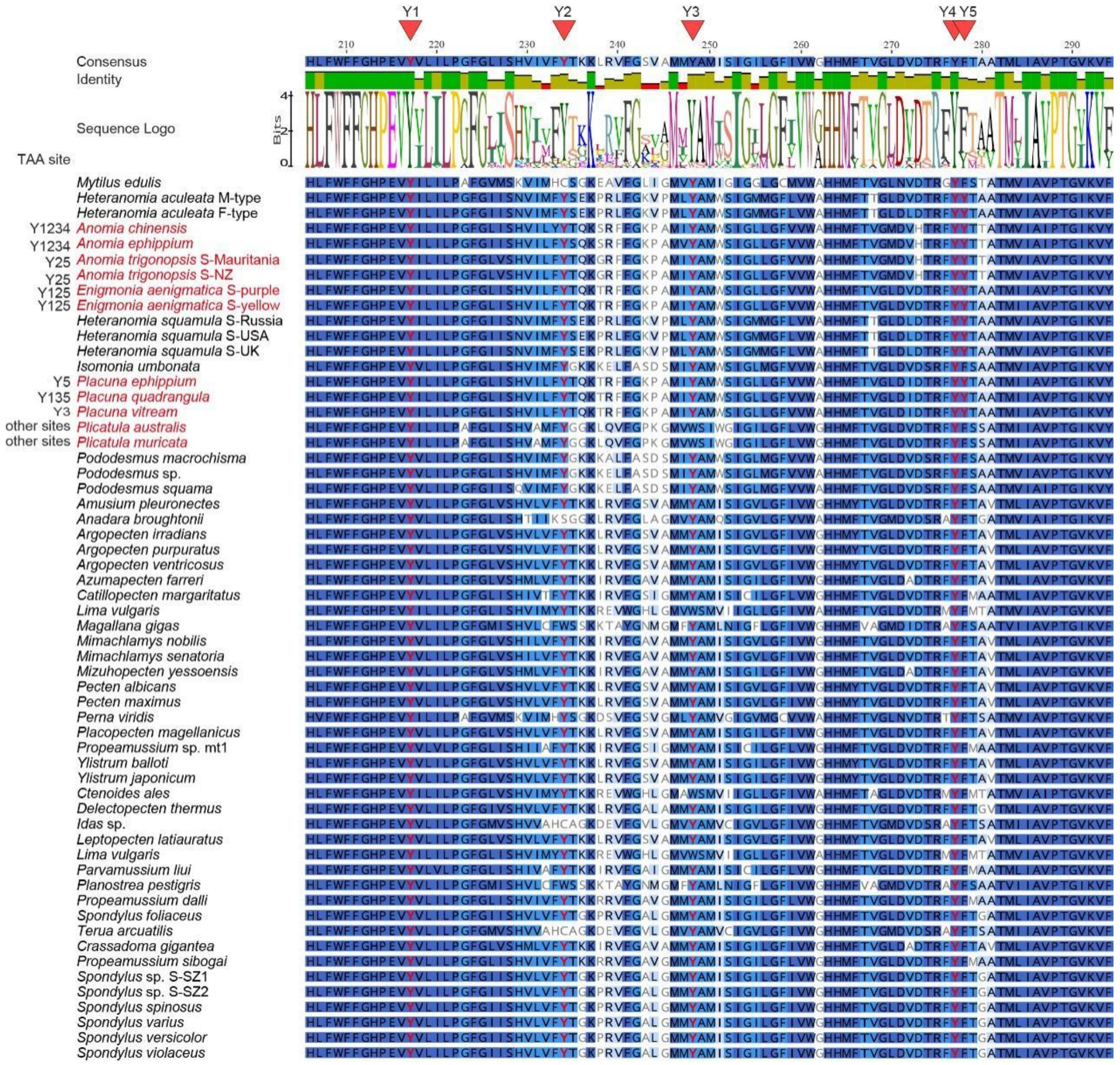


**Fig. S8** Partial multiple sequence alignment and sequence logo of mitochondrial COX1 protein sequences. Red triangles indicate five amino acid positions (Y1–Y5) where the majority of examined species encode tyrosine (Y). For species carrying in-frame TAA codons at these positions, the corresponding Y1–Y5 labels are shown to the left of the species name, indicating the presence of TAA codons rather than alternative tyrosine-encoding codons (TAC or TAT) at these sites.

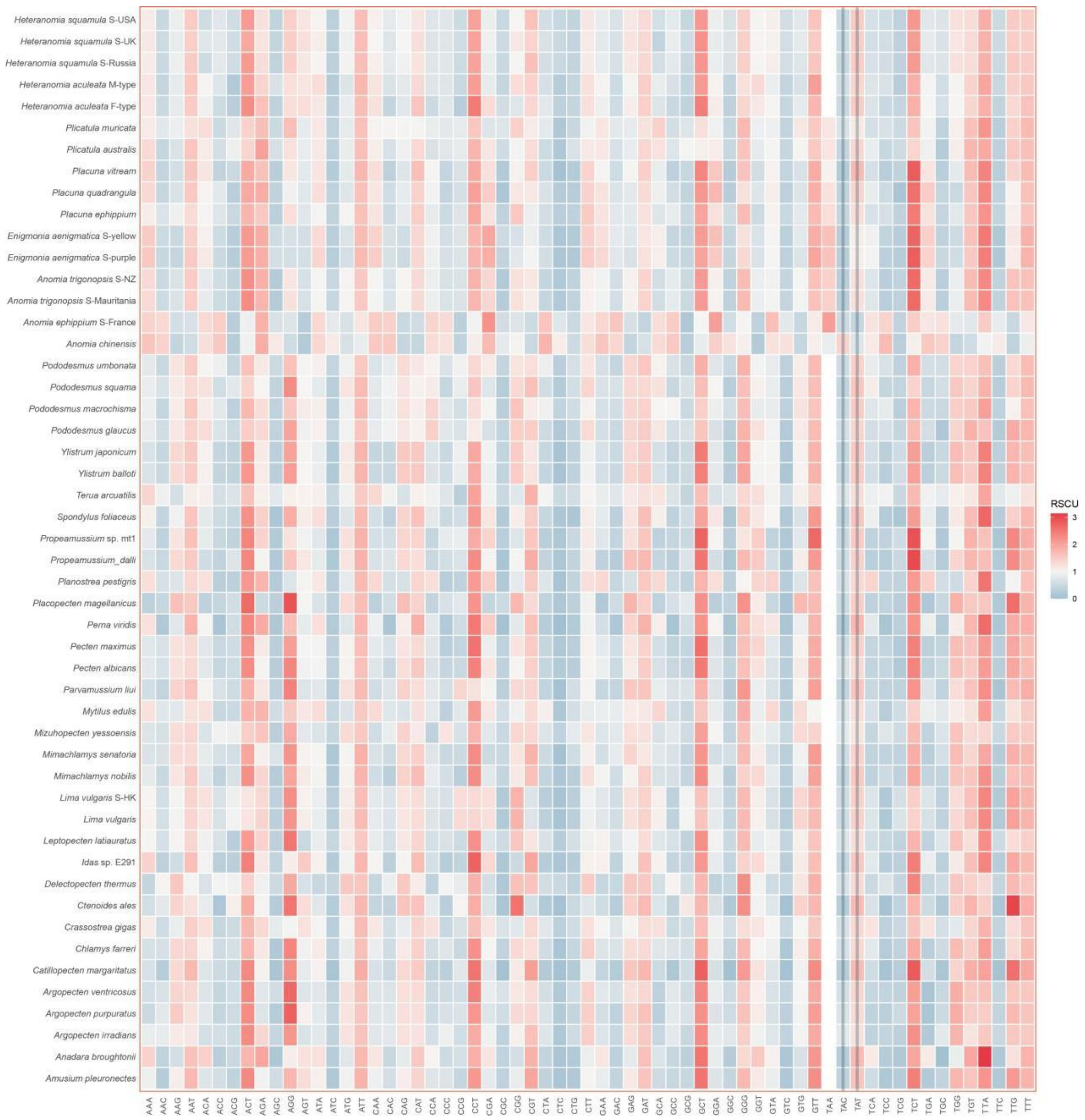


**Fig. S9** Heatmap of relative synonymous codon usage (RSCU) values for all protein-coding genes (PCGs) across the examined mitochondrial genomes. RSCU values indicate the frequency of each synonymous codon relative to the expected frequency under uniform codon usage; values greater than 1 denote preferred codons, whereas values less than 1 indicate codons used less frequently than expected.

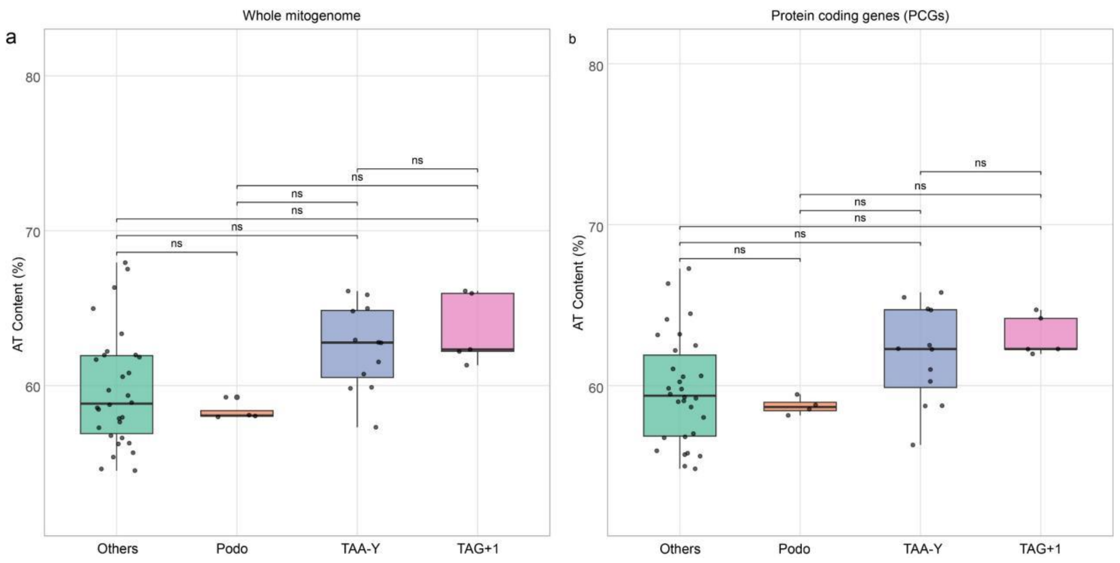


**Fig. S10** Boxplots comparing AT content of whole mitochondrial genomes (a) and protein-coding genes (PCGs) (b) between the APP lineage and related groups. No significant differences were detected among any of the groups (Kruskal-Wallis test, $p > 0.01$). Species are grouped according to their mitochondrial genome configurations: TAA=Tyr reassignment, TAG+1 translational frameshifting, *Pododesmus*-type (code 5), and other code 5 lineages (Pectinidae, Limidae, Ostreidae, Mytilidae).

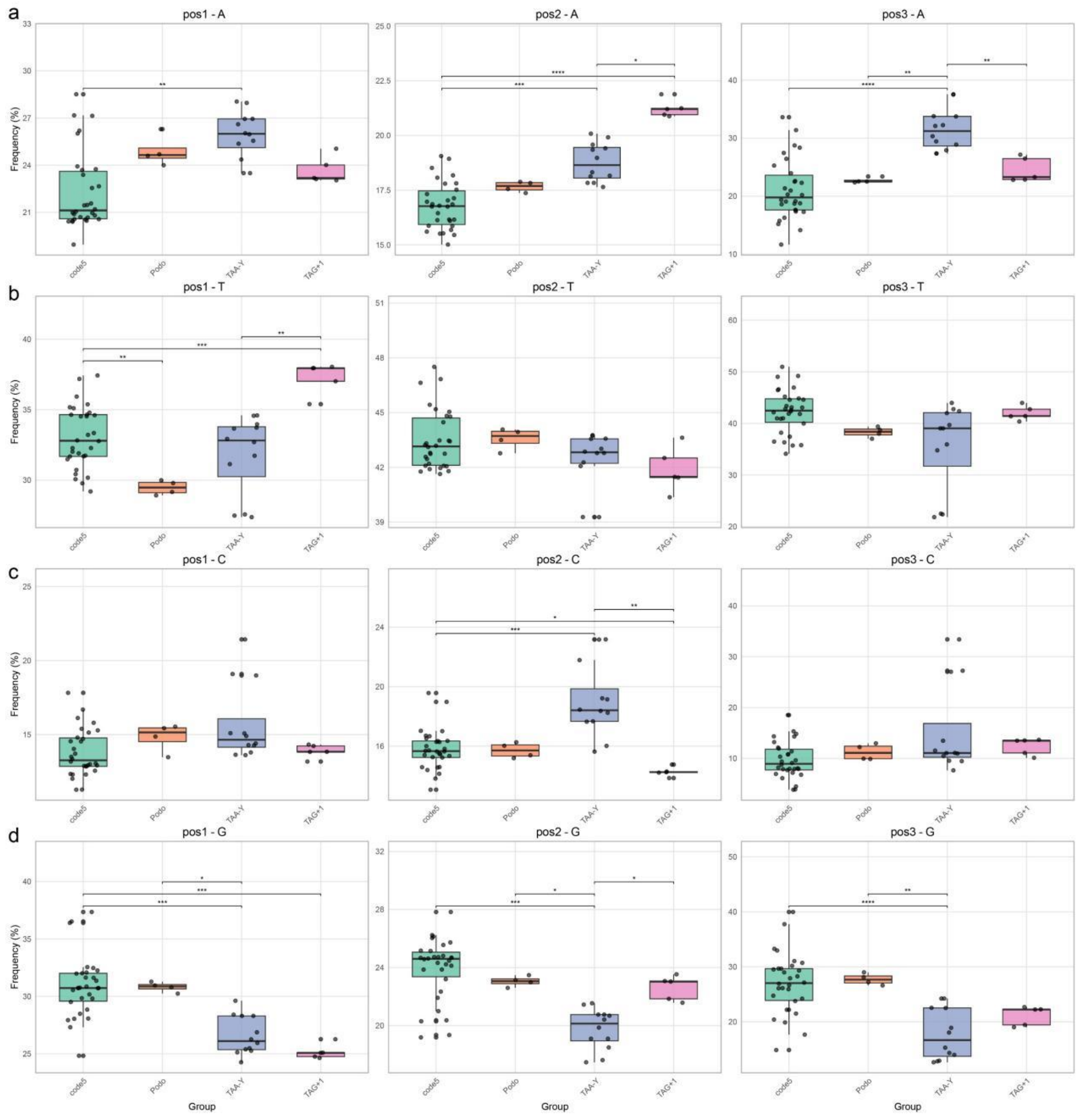


**Fig. S11** Boxplots comparing nucleotide composition at the third codon position of protein-coding genes (PCGs) among species with distinct mitochondrial genome configurations. Pairwise comparisons were performed using the Wilcoxon rank-sum test with Bonferroni correction, with significance levels indicated as follows: $p < 0.01$, $p < 0.001$, $p < 0.0001$. Species are grouped according to their mitochondrial genome configurations: TAA=Tyr reassignment, TAG+1 translational frameshifting, *Pododesmus*-type (code 5 and other code 5 lineages (Pectinidae, Limidae, Ostreidae, Mytilidae).

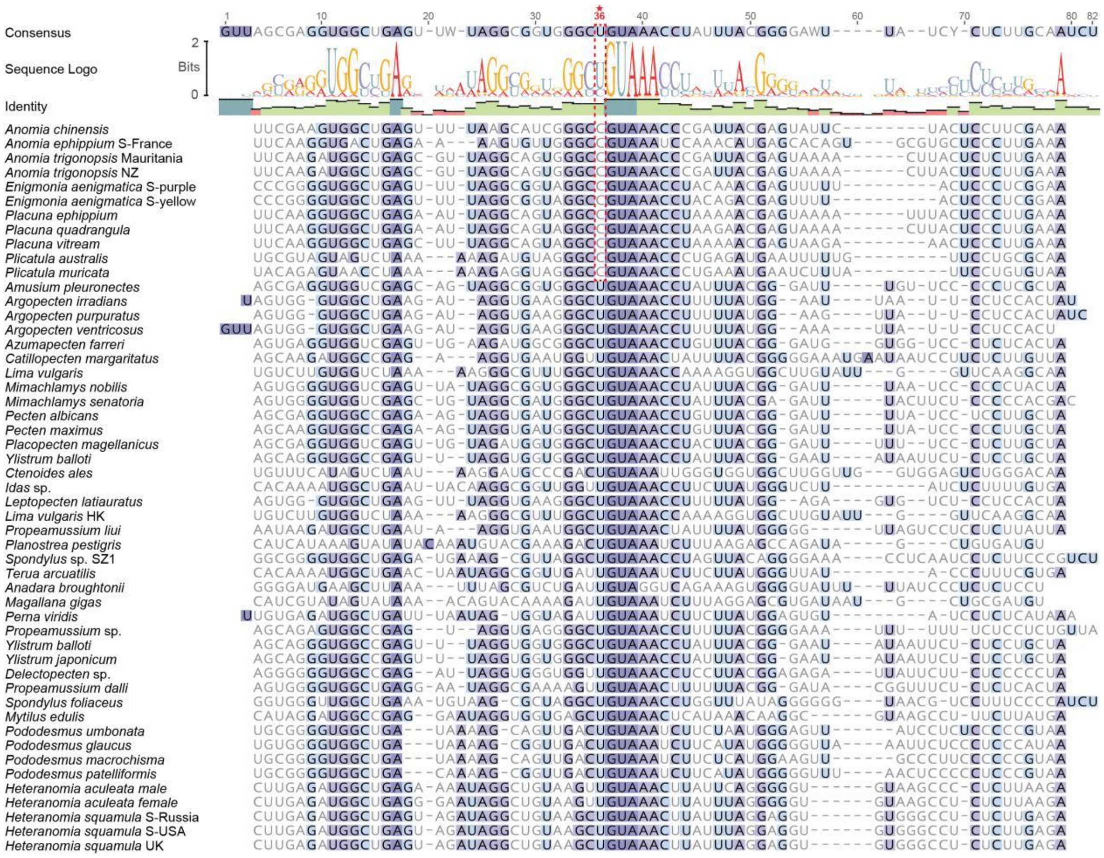


**Fig. S12** Multiple sequence alignment and sequence logo of mitochondrial *trnY* genes across the examined species. An asterisk indicates the nucleotide immediately upstream of the anticodon GUA (position 36). In all TAA=Tyr lineages, this position is occupied by cytosine (C), whereas in species using the invertebrate mitochondrial genetic code (code 5), it is occupied by uracil (U).

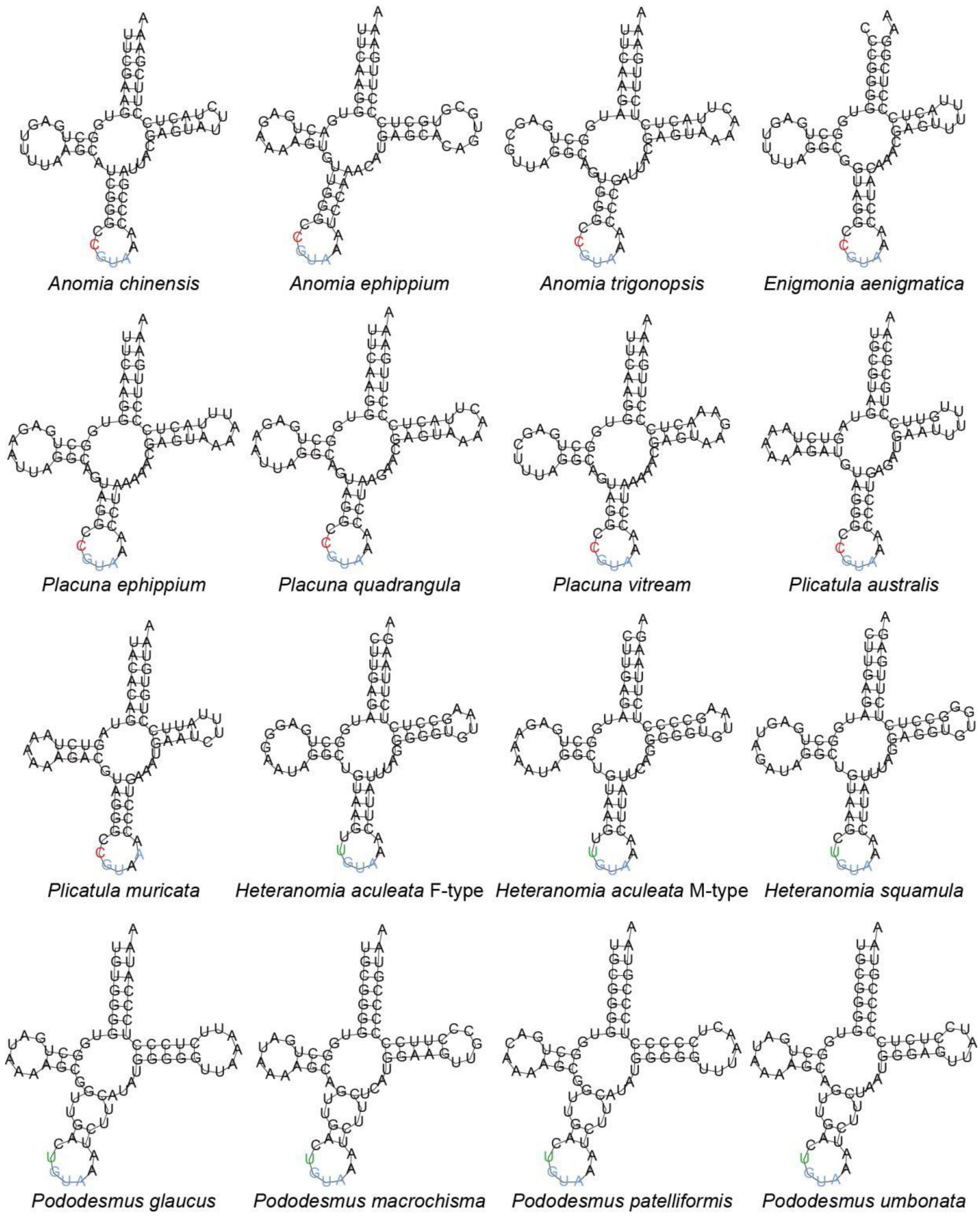


**Fig. S13** Predicted secondary structures of mitochondrial *trnY* in the APP lineage species. The U36C substitution, characteristic of TAA=Tyr lineages, is highlighted in red.

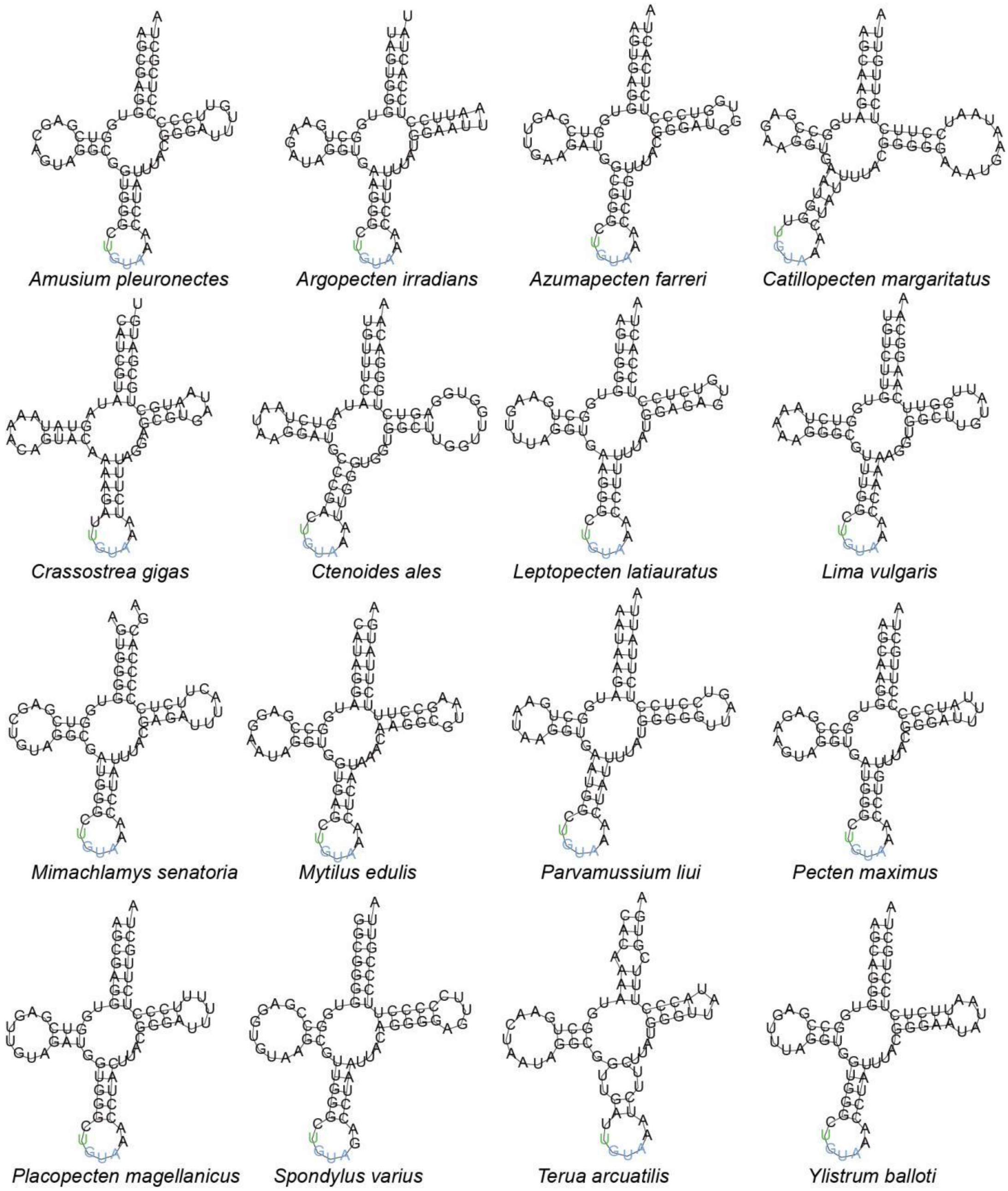


**Fig. S14** Predicted secondary structures of mitochondrial *trnY* in the species of Pectinidae, Limidae, Ostreidae, and Mytilidae that employ the invertebrate mitochondrial genetic code.
